# The interplay between biomechanics and cell kinetics explains the spatial pattern in liver fibrosis

**DOI:** 10.1101/2025.07.31.667562

**Authors:** Jieling Zhao, Seddik Hammad, Mathieu de Langlard, Pia Erdoesi, Yueni Li, Paul Van Liedekerke, Andreas Buttenschoen, Manuel Winkler, Sina W. Kürschner, Philipp-Sebastian Reiners-Koch, Niels Grabe, Björn Hartleben, Stephanie D. Wolf, Johannes Bode, Jan G. Hengstler, Matthias P. Ebert, Steven Dooley, Dirk Drasdo

## Abstract

The formation of liver fibrosis patterns, characterized by excess extracellular matrix (ECM), is a complex process that is difficult to investigate experimentally. To complement experimental approaches, we developed a digital twin (DT) model to simulate the pattern formation of septal and biliary fibrosis, the two common forms of liver fibrosis. This model is based on iterative calibration with experiments from animal models treated with the hepatotoxic substance CCl_4_ (septal form) and Abcb4-knockout mice (biliary form). Septal fibrosis is characterized by ECM accumulation along the connective line between the central veins of neighboring liver lobules, while biliary fibrosis is marked by a scattered ECM pattern within the portal fields. This mechanistic DT model includes the components of hepatocytes (Heps^♠^), hepatic stellate cells (HSCs), macrophages (Mphs), bile duct (BD) cells, collagen fibers secreted by activated HSCs, blood vessels, and cell-cell communication. It allows for the integration and simultaneous modulation of multiple hypothesized mechanisms underlying fibrotic wall formation.

The model simulates the formation of liver fibrosis pattern and demonstrates that ECM distribution results from the pattern of cell death zones and biomechanical compression due to cell proliferation. "Healthy" Heps proliferate to compensate for cell loss. In septal fibrosis, where the cell death zones are several cells thick, the proliferating Heps surrounding a zone mechanically compress the deposited collagen network. After a transient phase of collagen scattered between/around Heps, the ECM eventually adopts a sharp, "wall"-like structure. Whereas, in biliary fibrosis, the pattern of cell death is more scattered, leading to a corresponding scattered ECM pattern. In this case, a pattern of scattered distributed collagen forms without transitioning to a sharp wall. Notably, the failure of fibrotic wall formation in endothelial cell-specific GATA4^LSEC-KO^ mice, due to the disrupted pattern of CYP2E1-expressing Heps, validates our DT model.

In conclusion, the DT model provided a deeper understanding of liver fibrosis pattern formation. It enabled comparison between simulated outcomes of hypothesized mechanisms and experimental data. Additionally, it guided the design of validation experiments and enabled the identification of optimal strategies for drug testing and extrapolation to humans.

## Introduction

Chronic liver diseases (CLD) have diverse etiologies, including viral and other infections, autoimmune disorders, alcohol consumption, malnutrition or overnutrition, cholestasis, drug abuse, and genetic abnormalities. Regardless of the underlying etiological factor, chronicity reflects a prolonged history of parenchymal cell injury and a tissue response characterized by cycles of regeneration, inflammation, and fibrosis^1^. The damage is "repaired" through excessive extracellular matrix (ECM) deposition. However, upon repeated liver injury, this leads to tissue fibrosis and ultimately cirrhosis^2^, a major risk factor for hepatocellular carcinoma^3^. Histological assessment of liver biopsy remains the gold standard for staging CLD.

The prevailing concept of liver fibrogenesis suggests that injured hepatocytes (Heps) release damage-associated molecular patterns (DAMPs), which are recognized by neighboring liver cells, thereby triggering fibrotic signaling cascades. Liver macrophages (Mphs) produce fibrogenic cytokines, such as TGFβ and PDGFβ, which activate and recruit hepatic stellate cells (HSCs) that produce and deposit ECM proteins^4,5^. Mphs include the liver-resident Kupffer cells and recruited monocyte-derived Mphs, and they play diverse roles in the onset, progression, and regression of liver fibrosis by switching between inflammatory and restorative phenotypes^6-8^. HSCs exhibit significant plasticity, transitioning between quiescent (qHSC), activated (aHSC), and reverted (rHSC) subtypes^9-11^. During fibrosis regression in the absence of further injury, aHSCs are depleted, with approximately 50% undergoing cell death and 50% reverting to the qHSC phenotype^10^. The changes in HSC phenotypes are driven by molecular signals and ECM alterations, which have mechanical consequences on tissue stiffness and viscoelasticity^12-14^. Despite these and other investigations^2,15-17^, the orchestration of the processes that lead to the spatial patterns of fibrosis remains poorly understood, limiting the development of improved therapeutic strategies to prevent fibrosis progression toward cirrhosis and hepatocellular carcinoma.

Time-course data on the dynamics of cellular fate changes are primarily generated from experimental models i.e. mice. However, due to the complexity of the system, these models do not permit the simultaneous measurement of all potentially relevant process variables. As a result, they often fail to establish clear cause-effect relationships. In patients, the situation is even more challenging, as insights into the dynamics of liver disease development, progression, and regression are obtained indirectly through longitudinal blood sampling or non-invasive imaging methods such as MRI or ultrasound. However, blood parameters do not exclusively reflect disease activity in the liver, and imaging techniques can currently only provide information on morphological and biomechanical tissue properties at super-lobular scales. Information about the liver micro-architecture is typically obtained from resected or biopsied tissue, which is available only at a limited number of selected time points.

To better understand the complex interplay of processes and components involved in fibrosis initiation and progression, we created a computational digital twin (DT). This DT allows for the simulation of hypothesized mechanisms and supports data interpretation and the design of data acquisition strategies. The DT enables us to overcome the bottleneck of insufficient manipulability and measurability of the in vivo liver system, both in human and animal models, which can lead to misleading conclusions. For example, a consensus model of ammonia detoxification proposed by Häussinger^18^, Gebhardt, and Mecke^19^ has been shown through a DT model to quantitatively disagree with experimental data^20^, prompting revisions to the underlying mechanisms^21^ and the suggestion of a potential therapy for hyperammonemia. Without the DT model, this discrepancy would not have been identified. The DT thus serves as a valuable strategy to explain animal model data, in vitro experiments and can be translated to solve a clinically oriented problem i.e. drug testing.

The presented DT model is capable of resolving the spatial-temporal organization of liver tissue microarchitecture, as determined from histological tissue samples or intravital imaging—both of which require invasive interventions. It achieves this by representing each cell individually in a cell-based (agent-based) simulation approach, either at high resolution to capture cell shapes in regions where interactions with the ECM are critical, or at low resolution in areas sufficiently distant from the tissue damage zone. The basic liver structural unit of our DT is the liver lobule, which represents the smallest functional anatomical unit of the liver parenchyma. It typically has a hexagonal shape, with the central vein (CV) at its center, collecting blood that enters the lobule through branches of the hepatic artery and portal vein (PV) to the portal regions of the liver (**Fig. 1A**). Our DT to study fibrosis focuses on a pair of adjacent liver lobules, the minimal tissue unit required to explain the dynamics of fibrotic sharp wall formation in septal fibrosis or the scattered collagen pattern observed in biliary fibrosis.

**Figure 1.**
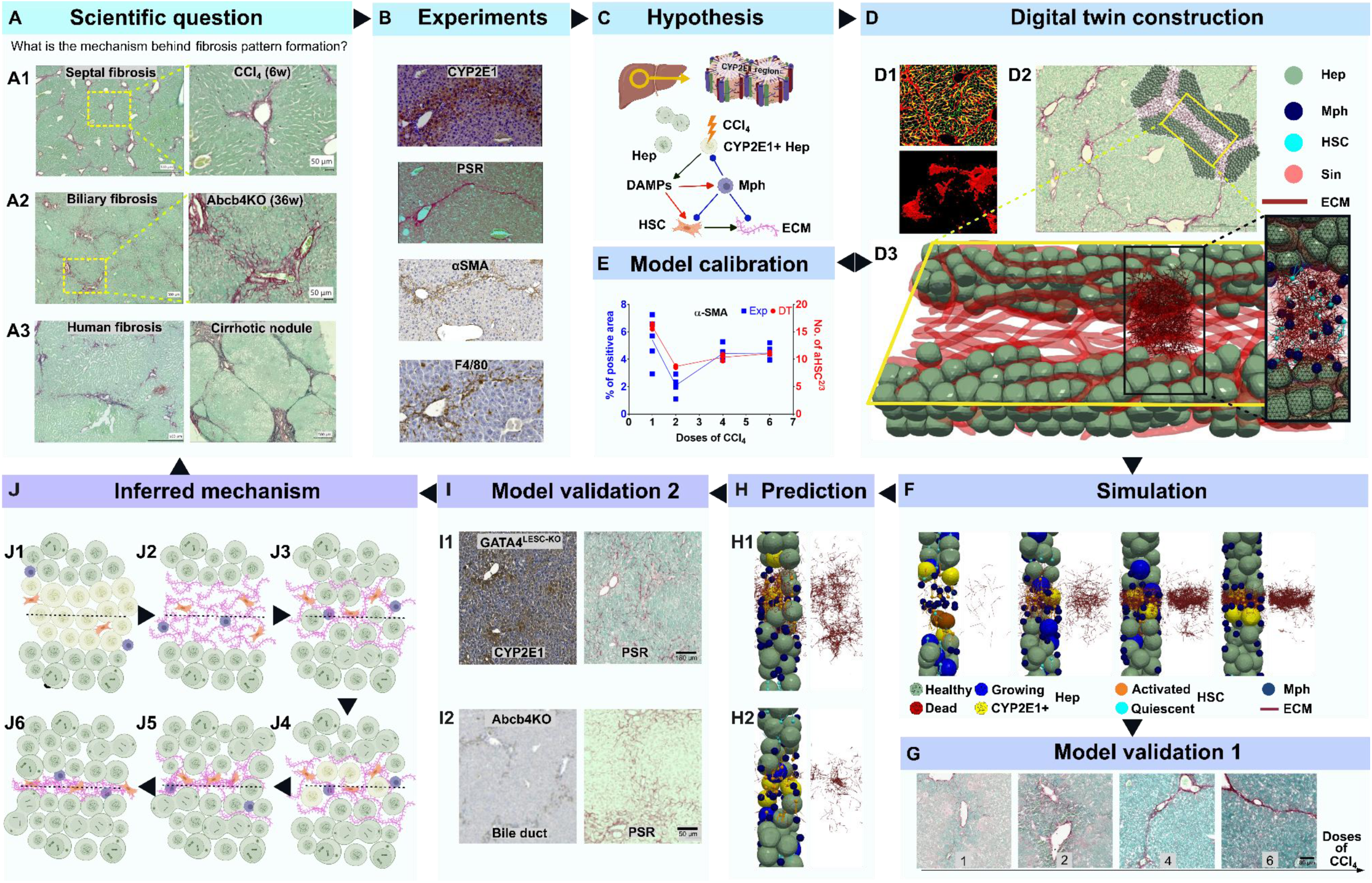
Workflow. (A) Liver fibrosis patterns – Pathomorphology in murine and human specimens (PSR staining of collagen). (A1) Septal fibrosis after hepatotoxic drug CCl_4_ administration (or equivalently, acetaminophen) in mice, (A2) Biliary fibrosis in Abcb4-knockout mice (Abcb4*KO*), (A3) Fibrosis pattern in humans. The scientific question addressed in this study: what is the mechanism behind the formation of liver fibrosis patterns? (B) Based on experimental data, hypotheses were formulated regarding interactions between major components (e.g., cells, signals, ECM) during fibrosis progression (C). (D) These hypotheses were implemented in a digital twin (DT) model. (D1) Staining of the collagen and the view of septal fibrotic wall in 3D (mice). (D2) The DT focuses on the region between the central veins (CVs) of two neighboring liver lobules, where septal fibrosis forms. (D3) The DT includes major cell types: Hep, Mph, HSC, and key structural elements: Sinusoids (Sin) and ECM. (F) The spatial-temporal distribution of cell subtypes in the DT model were calibrated to fit some selected experimental observations (e.g., the quantity of HSCs and Mphs over time). (F) DT simulations of septal fibrosis formation were compared to and validated by experimental observations not used for model calibration (G). (H) Prediction of perturbation tests: (H1) Prediction of the effect of GATA4 knockout (GATA4^LSEC-KO^, leading to perturbed expression and spatial distribution of CYP2E1) on septal fibrosis formation. (H2) Prediction of the effect of FAK inhibition on HSC activity. (I) Experimental validation of predictions: (I1) GATA4^LSEC-KO^ mice experiment to validate (H1). (I2) Abcb4*KO* mice experiment to validate biliary fibrosis pattern formation. (J) Inferred mechanism of septal liver fibrosis formation due to interactions between cells and structural elements. The black dashed line in each scheme represents the CV-CV axis orientation. (J1) CCl_4_ induces death of Heps expressing CYP2E1 (CYP2E1+ Heps, light green). After the first round of regeneration, the CYP2E1+ Heps form a pattern linking an increasing number of CVs of neighboring liver lobules. The damaged Heps attract HSCs and Mphs to the lesion. (J2) Activated HSCs produce collagen fibers (dark pink) in the dead cell region, while Mphs degrade the collagen. (J3) “Healthy” Heps surrounding the lesion proliferate (dark green with multiple points inside) to close the lesion. The newly generated Heps infiltrate the collagen network, compressing it into intercellular spaces, forming a “chicken-wire” pattern (collagen wrapped around hepatocytes). With further rounds of CCl_4_ administration, the death of CYP2E1+ Heps (J4), collagen production, and Hep proliferation lead to a gradual transition (J5) from the chicken-wire pattern to a sharp wall-like structure (J6), formed by proliferating Heps compressing the collagen network. (The mechanism for the formation of biliary fibrosis patterns is shown in Figure 6.) Abbreviations: Hep (hepatocyte), HSC (hepatic stellate cell), Mph (macrophage), ECM (extracellular matrix), CV (central vein), CYP2E1 (Cytochrome P450 2E1), Abcb4 (ATP Binding Cassette Subfamily B Member 4).

The DT is based on consistent computation of biomechanical forces, considering cell and ECM displacements and deformations in realistic and representative 3D liver microarchitectures. These microarchitectures are derived from 3D volume data sets of liver tissue. In the DT, ECM is simplified as the collagen network. Collagen fibers are modeled as linked, semi-flexible polymer-like objects^22-24^. Liver fibrogenesis is simulated as a dynamic, tightly regulated process driven by continuous crosstalk between hepatic cell types and interactions with structural components such as collagen.

The DT integrates HSCs, Mphs, cell-cell communication, collagen production and degradation to realistically mimic fibrosis onset and progression. Earlier DT models focused on acute liver injury, where collagen and largely deformable cells were not as critical. The integration of models for both acute and chronic liver damage marks a fundamental step toward a full DT (**Supplementary Fig. 1**) for liver fibrosis.

Despite the apparent complexity of the DT, its construction and the selection of model components are guided by parsimony (details in SI). In brief, the minimal set of model components was chosen based on their indispensability for explaining fibrotic wall formation. These components and their dynamics were parameterized as much as possible using measurable quantities, enabling direct measurement or identification of their physiological ranges. To account for potential mechanical inhibition of cell-cell rearrangement due to the sinusoidal network, the model had to be three-dimensional (Drasdo and Zhao^25^).

By integrating biomechanical and biological mechanisms, the simulation results from the DT, combined with animal experiments, led to the conclusion that repetitive Hep death in strictly localized regions of the liver tissue triggers collagen deposition to stabilize the dead cell lesions. Proliferation of surviving Heps at the boundary of the dead cell lesions shapes the final distribution of collagen fibers. If the dead cell lesion is several cells thick, proliferating Heps displace the deposited collagen towards the center of the lesion through mechanical forces, eventually generating localized fibrotic walls characteristic of pericentral/septal fibrosis (**Fig. 1A**).

Localized fibrotic "walls" are not exclusive to the liver; they can also be observed in other organs, such as in myocardial^26^ and pancreatic fibrosis^27^. The model predicts that the final collagen distribution pattern reflects the shape of the damage zone. For instance, in biliary fibrosis, where cell death occurs in a scattered pattern, a localized fibrotic wall is not formed. Our DT can capture this etiology as well.

## Results

### (I) Different etiologies of liver damage induce variant patterns of fibrosis

#### In the first step, we compare fibrosis patterns associated with different etiologies to generate potential hypotheses for the formation of pericentral/septal fibrosis, which are then implemented in the DT model

Liver fibrosis can manifest as pericellular (in the Disse space), septal, biliary, and bridging types. In septal fibrosis, liver damage occurs in the pericentral compartment and forms fibrotic walls between the central veins (CV) of adjacent lobules. Chronic CCl_4_ intoxication is a well-established animal model for studying liver fibrosis and testing the possible inhibitors as anti-fibrotic drugs^2^.

Picro-Sirius Red (PSR) staining of the liver after 12 CCl_4_ injections over 6 weeks reveals a fibrotic wall between central veins (CVs), forming so-called "pseudolobules" (**Fig. 1A1**). In biliary fibrosis, as seen in the Abcb4*KO* mouse model, collagen deposition begins periportally and extends towards adjacent portal veins in advanced disease (PV; **Fig. 1A2**^28^). In CCl_4_-induced fibrosis, crosslinked collagen fibers eventually form a thin, sharp wall-like structure, whereas in Abcb4*KO* livers, collagen fibers remain scattered and form a periportal fibrosis area, where a broader network forms between portal fields (**Fig. 1A2**). Early liver fibrosis in humans appears in both pericentral and periportal regions (left, **Fig. 1A3**) or later bridging (connecting fibers) between pericentral and periportal regions, depending on the underlying etiology, ultimately leading to cirrhotic nodules in late-stage disease (**Fig. 1A3**). In conclusion, fibrosis patterns vary based on disease etiology and the type of injury.

### (II) Experiments suggest fibrotic walls emerge from cycles of CYP2E1+ hepatocyte (Hep) death and proliferation of healthy Hep

Currently, our understanding of the mechanisms underlying the distinct ECM deposition patterns is limited. The sharp localization of ECM, forming a narrow wall connecting two lobules, as seen in CCl_4_-induced fibrosis, is absent in all disease stages of the Abcb4*KO* mouse model but is characteristic of advanced fibrosis in human patients, such as those with NASH (Nonalcoholic Steatohepatitis) or MASH (Metabolic Dysfunction-Associated Steatohepatitis) (**Fig. 1A3**). To investigate the dynamics of liver fibrosis, we analyze the spatial distribution of CCl_4_-metabolizing (CYP2E1+) Heps in relation to collagen deposition, using (immuno)staining of serial sections with PSR, CYP2E1, and CK19, the latter marking bile duct epithelial cells typically found in periportal regions. Our observations show that the fibrotic wall in CCl_4_-induced fibrosis forms within the region of CYP2E1+ Heps, which localize around and form stripes connecting CVs following repeated CCl_4_ injections (**Supplementary Fig. 2**). This suggests that the observed septal fibrosis pattern is closely linked to the dynamics of metabolic activity (CYP2E1 expression), Hep damage, and subsequent proliferation in response to toxin-induced injury. This leads to our ***main hypothesis***: the sharp fibrotic wall in septal fibrosis results from cycles of CYP2E1+ Hep death and the proliferation of healthy cells replacing the dead areas, with excess ECM deposition not being sufficiently degraded. Currently, our understanding of the mechanisms underlying the distinct ECM deposition patterns is limited.

### (III) DT-based strategy to uncover the mechanism underlying fibrotic wall formation from cycles of Heps birth and death

To address the complexity of this scenario and continuously integrate and exchange potential components that may explain septal/biliary fibrosis pattern formation, we adopt a systems biology/medicine approach and develop a liver Digital Twin (DT) with minimum lobule units to simulate fibrosis pattern formation. This model simulates the observed and hypothesized process steps leading to fibrotic wall formation, encompassing intercellular signal processing, cell-cell communication, and tissue biomechanics. The key components of the DT include various liver cell types, fenestrated liver capillaries (sinusoids), and collagen fibers (**Fig. 1D**). All components are integrated into a model spanning adjacent hepatic lobules to capture the fibrogenesis process. We focus on a DT that considers two neighbouring liver lobules, as the minimal fibrotic wall unit in septal fibrosis connects two central veins (**Fig. 1D**). The micro-architecture and cell composition within the DT unit are designed to represent the experimental equivalent of the liver’s starting state, prior to the first administration of CCl_4_.

Next, we gather data to construct the spatial-temporal microarchitecture of liver fibrosis in the DT. Our strategy involves calibrating the DT using data from acute liver injury following a single dose of CCl_4_. Once calibrated, the DT is used to predict the remodeling of liver tissue microarchitecture after repeated doses of CCl_4_. These predictions are then compared with experimental findings obtained after repeated CCl_4_ injections (twice a week on days 3 and 7 of each week for 3 weeks).

### (IV) Data-driven generation of a DT model after a single CCl_4_ injection

In the next step the components of the DT are introduced, beginning with the tissue microarchitecture i.e. sinusoidal scaffold. This is followed by the inclusion of Heps in various states, such as CYP2E1+ and CYP2E1-healthy, damaged (or dead), and proliferating. Then, non-parenchymal cells, including HSCs and Mphs, are incorporated, along with the ECM. Each model element is described with its spatial representation and potential temporal dynamics. The selection of these model constituents is based on quantitative imaging of their respective structures.

#### A) Sinusoidal scaffold

Previous research has shown that the sinusoidal network—the fenestrated capillary network connecting the portal field to the central vein within each liver lobule maintains its principal structural framework for liver regeneration after CCl_4_ treatment^29^. Based on this, we use this scaffold structure for the cross-lobule DT. The sinusoidal scaffolds are modeled as a network of semi-flexible chains of small spheres interconnected by springs (depicted in transparent red, **Fig. 1D3**). This model allows the formulation of force balance equations for each sphere within the sinusoidal network, analogous to those governing the Heps. This framework enables the realistic simulation of sinusoidal displacements by incorporating compressive forces exerted by adjacent Heps, which lead to a reduction in the apparent sinusoidal diameter and effectively reproduce physiologically relevant compression dynamics.

#### B) Hepatocytes (Heps)

To experimentally determine the spatial-temporal distribution of different liver cell types, we use mouse livers from healthy controls and those harvested 3 days after a single CCl_4_ injection. This time point serves as a reference state for establishing the initial configuration of the DT. Day 3 is specifically chosen because it marks the onset of fibrosis development, which begins following the second dose of CCl_4_. After a single CCl_4_ injection only, the liver undergoes complete regeneration of the lesion by day 7^29^.

We quantify the proportions of healthy, necrotic, CYP2E1-expressing, and proliferating (BrdU-labeled) Heps in the reference state. Pericentral Heps undergo necrosis due to CCl_4_ metabolism. At the time point of the first dose about 40% of the total area exhibits CYP2E1 positivity, dropping to 8.86 ± 2.45% in the reference state (**Fig. 2A, B**). In contrast, Heps outside the CYP2E1+ zone remain healthy throughout the process. This is demonstrated and quantified on day 3 through HE and IgG staining (**Fig. 2A, B**). Notably, the highest fraction of necrotic cells is observed 1 day after the initial dose, in agreement with prior findings^29^.

**Figure 2.**
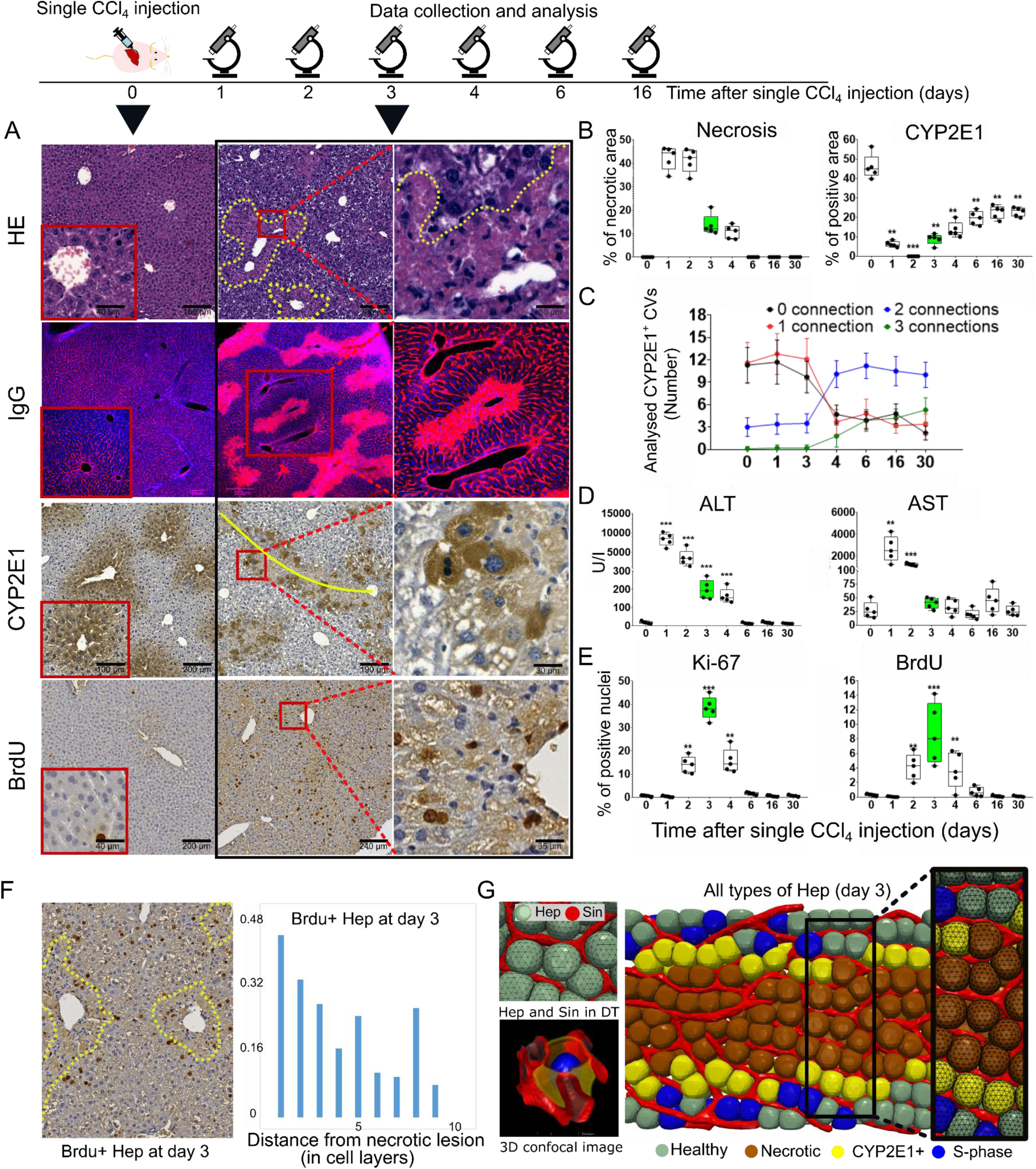
Experimental data of Heps at day 3 after a single dose of CCl_4_ and the Hep representation in the Digital Twin (DT). (A) Histopathological investigations at day 3 after a single injection of CCl_4_ (the right 2 columns in black box) compared to the control (day 0, the leftmost column); HE (light purple, lesion boundary shown as dashed yellow line), IgG for necrosis/cell death, CYP2E1 (the full yellow line marks the connection between two neighboring CVs), the CCl_4_ metabolizing enzyme, and BrdU, for nuclei in S-phase. (B-E) Quantification of necrotic area, CYP2E1, number of CV-CV connections by CYP2E1+ Heps (day 0 corresponds to the healthy case), blood levels of liver enzymes ALT & AST, Ki-67, and BrdU after single CCl_4_ injection, respectively. (F) Example of BrdU stained Hep nuclei (dark brown, lesion boundary shown as dashed yellow line) (left). (Right) The fraction of BrdU-positive Heps versus the distance of the lesion boundary (the yellow line) at day 3 after a single CCl_4_ injection^29^. (G) Components of the DT, including Heps and sinusoidal network (Sin). The left side displays a few Heps in the deformable cell model embedded in a sinusoidal network structure (upper picture), and a reconstructed Hep from confocal microscopy (lower picture). Data is collected from 5 mice per time point. * P <0.05, ** P <0.01, *** P <0.001.

The disappearance of pericentral Heps is evidenced by the complete loss of CYP2E1 immunoreactivity (**Fig. 2A–C**), a marked decline in *CYP2E1* mRNA expression by day 2, and significantly elevated plasma levels of alanine transaminase (ALT) and aspartate transaminase (AST) on days 1 through 3. Liver enzyme levels normalize as regeneration progresses (**Fig. 2D**). Necrotic regions nearly close by day 5. CYP2E1 mRNA and protein expression do not fully return to their original levels and patterns, at least until day 30 in the regenerated liver. Furthermore, a layer of midzonal peri-necrotic Heps begins expressing CYP2E1 on day 3, forming a dumbbell-shaped region connecting adjacent central veins (brownish region surrounding the yellow center line of CYP2E1, **Fig. 2A**). The number of central veins connected by CYP2E1+ Heps remains elevated until day 30 in the regenerated livers (**Fig. 2C**). The remaining Hep fraction on day 3 exhibits an increased number of BrdU/Ki-67-positive cells/nuclei, which are not uniformly distributed within the hepatic lobule but are more prevalent in peri-necrotic areas (**Fig. 2A, B, E, F**; **Supplementary Fig. 3A**).

In the DT, each Hep is individually modeled as a single unit, with the spatial resolution of cell shape depending on its proximity to the emerging necrotic lesion (**Fig. 2G**). Heps located near the lesion, which interact with the collagen network during fibrosis formation, are simulated using a deformable cell model (DCM)^30^ that explicitly captures dynamic cell shape changes (**Fig. 2G**, **Supplementary Movie 1**). In contrast, Heps located further from the lesion and not in contact with newly synthesized collagen fibers are modeled using a center-based model (CBM), in which cellular forces are applied to the cell center. For simplicity, cells in the CBM are represented as spheres, reflecting the shape they would adopt in isolation.

CYP2E1+ Heps (yellow) form connections between two CVs (**Fig. 2G**). In contrast, CYP2E1-Heps located outside the lesion are classified as damage-resistant cells (green, **Fig. 2G**). Following CCl₄ injection, CYP2E1+ Heps undergo cell death and are labeled as necrotic cells (brown). A subset of the remaining Heps is stochastically selected to proliferate (blue) in order to close the necrotic lesion, based on their distance from the lesion and the time point after CCl_4_ injection. The spatial-temporal pattern of Heps proliferation is inferred from the quantification of Ki-67-positive cells, as previously described^29^. BrdU staining is primarily used to highlight the elevated number of proliferating Heps surrounding the necrotic regions.

#### C) Non-parenchymal cells and the extracellular matrix (ECM)

Next, we performed a spatial-temporal analysis of HSCs, which are known to contribute to both liver regeneration and fibrogenesis, following a single CCl₄ injection. This assessment employed a spatial-temporal approach, using α-SMA and desmin to label aHSCs and total HSCs (aHSCs + qHSCs), respectively. In addition, Picro-Sirius Red (PSR) staining was used to visualize collagen deposition, alongside further staining to identify Mphs (**Fig. 3A–E; Supplementary Fig. 4A)**. On day 3 post-injection, α-SMA+ cells accumulate in the region between neighboring CVs, spanning approximately four Hep layers (∼80 µm), which corresponds to the size of the necrotic lesion (**Supplementary Fig. 5**). Quantification shows that the stained area fractions for desmin (11.5 ± 3.4%) and α-SMA (3.86 ± 1.3%) reach their peak on day 3. Consistent with these findings, mRNA expression levels of *Desmin* and *α-Sma* are also significantly upregulated on day 3. At day 0, PSR-positive collagen fibers are primarily detected in the tunica adventitia of larger blood vessels, including CV, PV, and hepatic artery. In contrast, on day 3 following CCl₄ injection, small, isolated, and scattered collagen fibers are observed within the necrotic regions. This spatial pattern coincides with a transient increase in mRNA expression levels of Col1a1 and Col1a2 (**Fig. 3D**), indicating active collagen synthesis and deposition. Importantly, excessive collagen deposition has not been previously associated with acute toxic liver injury, and our findings are consistent with this notion. These results suggest that the isolated collagen fibers appearing within the necrotic zones on day 3 are rapidly degraded following their synthesis and subsequently diminish during liver regeneration.

**Figure 3.**
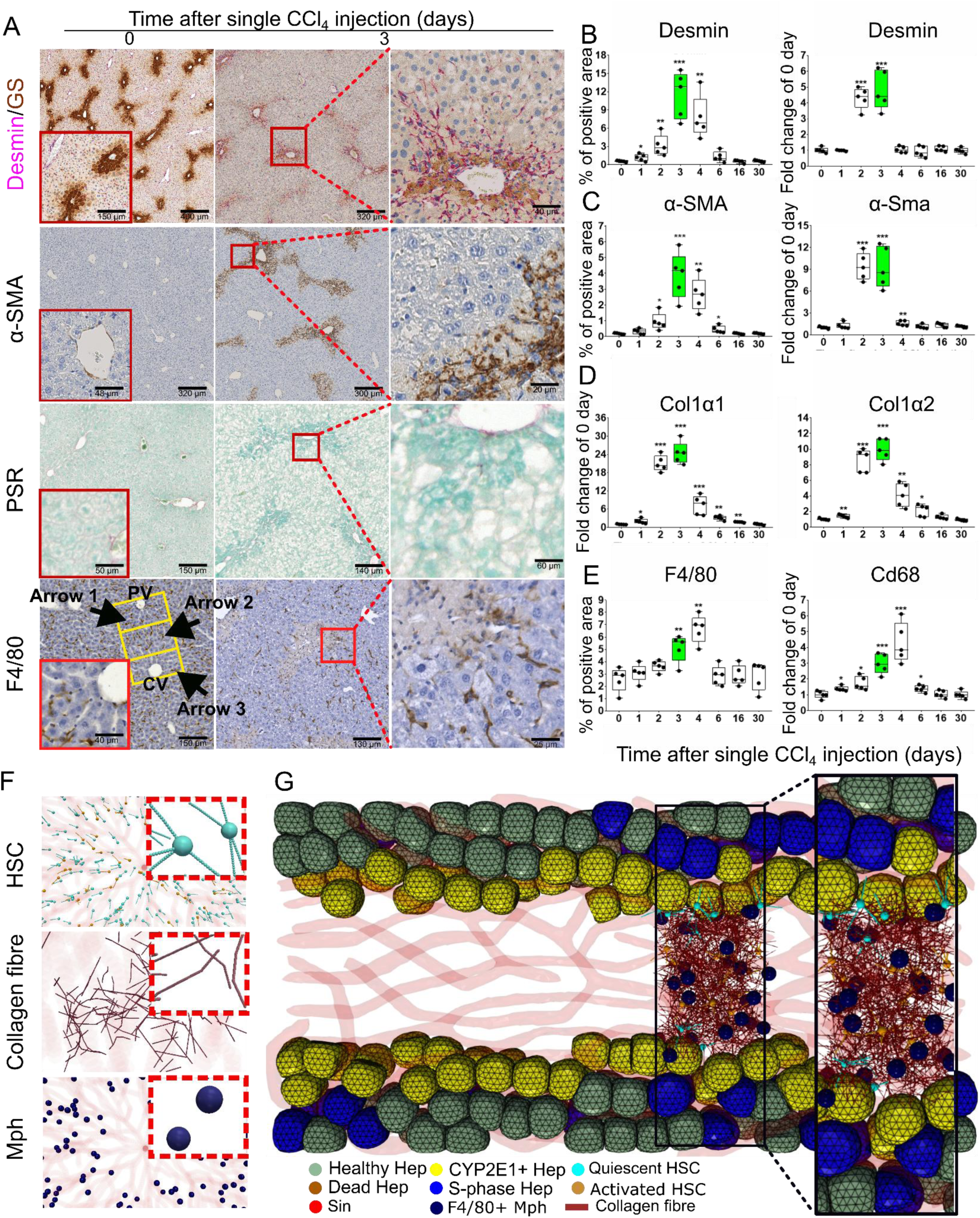
Experimental data on collagen and collagen modulator cells at day 3 after a single injection of CCl_4_ and representation in the Digital Twin (DT). (A) Time-resolved histopathological data after a single CCl_4_ injection, showing PSR (collagen deposition), desmin (quiescent and activated HSC), α-SMA (activated HSC), and F4/80 (stained in brown, resident Mph). Arrows 1, 2 and 3 point to F4/80-positive cells in periportal, midzonal, and perivenous regions, respectively (cf. Supplementary Fig. 4B). (B-E) Quantification of positive staining (left picture) and mRNA levels (right picture) for desmin (B), α-SMA (C), mRNA levels of Col1α1 and Col1α2 (D), and F4/80 (positive staining)/Cd68 (mRNA level) (E). (F) Components of the DT include HSC, Mph and collagen fibers (shown are small spatial tissue regions in the model and the model component in the inset). (G) A DT spanning two liver lobules that comprises all listed components. Data is shown from 5 mice per time point. * P <0.05, ** P <0.01, *** P <0.001.

In the DT, each HSC is modeled as a central spherical nucleus connected to multiple chains of elastic springs, representing the cell’s elongated protrusions. qHSCs are depicted in cyan, while aHSCs are shown in light brown (highlighted in dashed red boxes, **Fig. 3F**). Within the model, only aHSCs possess the ability to synthesize collagen fibers. These fibers are deposited into the extracellular space, where they progressively accumulate and interconnect to form a complex collagen network (**described in detail in the SI**).

The ECM is modeled as a network of semi-flexible chains of collagen fibers that require energy to bend and compress along their length, effectively capturing the biomechanical behavior of collagen bundles (depicted in purple, **Fig. 3F**; **Supplementary Fig. 6A**). The density of the collagen network is determined by the local concentration of collagen fibers. When the minimum distance between two fibers falls below a defined threshold, crosslinking occurs, represented by the formation of an additional connecting node (**Supplementary Fig. 6B**). The biomechanical properties of the collagen network are calibrated using data from in vitro bending^31^ and compression^32^ experiments. This calibration ensures that the simulated collagen network reproduces the experimentally observed biomechanical responses to deformation (**Supplementary Fig. 6C–G**). In the next stage, we hypothesize that cumulative collagen deposition is modulated not only by synthesis but also by degradation processes, such as those mediated by matrix metalloproteinases (MMPs) secreted by resident Mphs (Kupffer cells). To assess this, we analyze the spatial-temporal distribution of Mph infiltration using F4/80 immunostaining. In healthy livers, F4/80+ cells are primarily located in periportal (arrow 1, **Fig. 3A**, day 0 of F4/80) and midzonal (arrow 2) regions, with markedly fewer cells in pericentral zones (arrow 3) (**Fig. 3A**; **Supplementary Fig. 4A, B**). Following CCl₄ injection, a pronounced increase in F4/80-positive Mphs is observed throughout the entire lobule by days 2 and 3 (highlighted by black arrows, **Supplementary Fig. 4A**). This injury-induced redistribution persists for over 16 days before returning to a normal pattern. Gene expression analysis further supports this response, showing elevated levels of Cd68 (a Mph marker), as well as key ECM modulators such as Tgfβ1, Mmp9, and Timp1 within the first four days post-injury. In addition to engulfing necrotic Hep debris^33^, Mphs contribute to ECM remodeling by degrading collagen fibers^34^.

In the DT, two Mph populations are represented: resident and infiltrating ones. Resident Mphs are present in the liver under homeostatic conditions, whereas infiltrating ones enter the lobule only after each CCl₄ injection and undergo apoptosis thereafter^7^. In the model, Mphs are depicted as elastic spheres (dark blue, Mph, **Fig. 3F**). Upon contact, they phagocytose necrotic Heps and contribute to ECM remodeling by degrading collagen fibers. The experimental findings suggest a complex interplay involving Heps (healthy; damaged, therefor source of DAMPs; and proliferating), qHSC, aHSC, ECM, and Mph, forming a minimal set of building blocks to construct a dynamic model for studying the cycles of damage, regeneration/repair, and fibrogenesis after repetitive CCl_4_ injections (**Fig. 3G**). Notably, a subset of aHSCs is capable of reverting to a quiescent state through interactions with Ly6C^low^ Mph (see the SI). These reverted HSCs exhibit a “memory” phenotype, responding differently upon subsequent injury challenges compared to HSCs with original quiescent phenotype, as documented following the second CCl₄ injection^9-11^.

### (V) Implementation of processes pertaining to septal fibrosis formation dynamics and prediction of disease scenario

In the next step, we employed the DT to simulate the effect of six CCl_4_ injections over a three-week period, in alignment with the experimental protocol. CCl_4_ injections are scheduled twice a week, on days 0 and 3 of each week, with weekdays enumerated from 0 to 6. Accordingly, CCl_4_ injections are administered on days 0, 3, 7, 10, 14, and 17, followed by a final histological assessment on day 21. Each injection elicits damage primarily within the CYP2E1+ Hep population (**Supplementary Fig. 7A**). This damage is consistently accompanied by elevated plasma levels of AST and ALT (**Supplementary Fig. 7B**), compared to baseline levels observed in healthy animals (see time point zero in **Fig. 2D**). In the experimental observations, only the Hep layer directly adjacent to the necrotic lesion expresses CYP2E1 at day 3 following a single CCl_4_ injection. These necrotic areas are subsequently repopulated through Hep proliferation. Notably, the newly formed Heps uniformly express CYP2E1, indicating a reestablishment of pericentral metabolic zonation—and thus renewed sensitivity to subsequent CCl_4_ exposure. Over time, the CYP2E1+ Heps become spatially organized in a continuous stripe that connects two neighboring CVs. Importantly, the extent of this CYP2E1-expressing region remains consistent in size after 2, 4, and 6 CCl_4_ injections (**Supplementary Fig. 7A, C, D**).

In the DT, Heps that repopulate the necrotic lesion are assumed to express CYP2E1. This regenerated CYP2E1+ pattern drives the progression of tissue damage spatially-temporally, thereby influencing the distribution of DAMPs and subsequent cellular responses. These responses include the recruitment of HSCs and Mphs, deposition of collagen, and compensatory Hep proliferation, all of which contribute to the repeated restoration of the lesion.

The Hep proliferation profile is parameterized based on Ki-67 staining quantified on day 3 following the first CCl_4_ injection, in line with previous studies^29^. Experimental data on both the extent and spatial distribution of proliferation after repeated CCl_4_ administration indicate that healthy Heps surrounding the lesion re-enter the cell cycle. The proliferation rate is highest in Heps immediately adjacent to the lesion and gradually declines with increasing distance from the lesion boundary (**Fig. 2F**).

#### A) DT predicts intermediate chicken-wire pattern

In the DT, compromised Heps expressing CYP2E1 are assumed to serve as a source of DAMPs. Among other effects—DAMPs attract HSCs (**Supplementary Fig. 8A, B**), triggering their activation and migration into the lesion. These aHSCs subsequently produce collagen fibers, initially forming a characteristic "chicken wire" pattern that is recapitulated by the DT model (**Supplementary Fig. 8C, D**). This pattern emerges because, after the first CCl_4_ dose, collagen deposition is sparse and loosely organized. As proliferating Heps repopulate the lesion, they push the nascent collagen fibers into the intercellular space. With repeated CCl_4_ administration, collagen density and fiber connectivity progressively increase (**Supplementary Fig. 11D**), and proliferating Heps instead begin to push against an increasingly consolidated collagen network. Experimentally, scattered collagen fibers are observed one week after the second CCl_4_ injection, which progressively organize into a well-defined fibrotic wall after 4–6 doses. The evolving collagen deposition pattern parallels the spatial distribution of desmin- and α-SMA-expressing cells (**Supplementary Fig. 9A**). Temporally, mRNA expression levels of ECM-related genes such as *Col1a1* and *Col1a2* exhibit similar upregulation as those of *Desmin*, *α-Sma*, *Cd68*, *Mmp9*, and *Timp1*, observed at both transcription and protein levels (**Supplementary Fig. 9B–D**). Notably, no further accumulation of HSCs is detected following 4–6 doses of CCl_4_, as indicated by plateauing α-SMA expression at both mRNA and protein levels.

The mechanism underlying the spatial organization of secreted collagen fibers—and its dynamic transformation leading to the formation of fibrotic walls—remains poorly understood. We hypothesize that the enzyme CYP2E1 plays a central role in shaping the overall spatial patterning of the collagen and in regulating the localization of collagen-producing aHSCs, as indicated by desmin and α-SMA staining, despite their relatively dispersed appearance (**Supplementary Fig. 9A**).

To test this hypothesis, we simulate the full course of repeated CCl_4_ injections and the subsequent cascade of events within the DT. Due to uncertainty regarding the precise structural form of collagen within the lesion, we explore six alternative scenarios (S1–S6) of collagen deposition. These scenarios differ in terms of fiber structure, sinusoidal anchoring, and crosslinking dynamics (see the SI and **Supplementary Fig. 10A** for detailed descriptions). Among these, the scenario in which collagen is initially deposited as individual fibers that gradually crosslink into a network ("Scenario 5" in the SI) most closely aligns with the experimental observations. In this model, aHSCs produce and deposit collagen fibers, which progressively crosslink to form an interconnected collagen network.

#### B) Repeated rounds of CCl_4_-injection result in fibrotic wall formation in DT simulations through mechanical compression by Hep proliferation

Based on its consistency with experimental data, Scenario 5— was chosen as a collagen deposition scenario. Mph was incorporated into the DT model based on the spatial-temporal dynamics of F4/80-positivity, as observed through immunostaining (dark blue; **Fig. 3F, G**). In the model, these Mphs are assumed to degrade collagen fibers upon direct contact.Accordingly, collagen fibers begin to be produced by aHSCs after the second CCl_4_ injection, and are subsequently degraded by infiltrating Mph within the lesion. After six CCl_4_ doses, the total amount of collagen deposition is assessed both in the DT model and experimentally, using quantification of PSR-positive areas over time (**Fig. 4A, B**). The thickness of the collagen network in the DT model is regulated by the interplay between collagen production (by α-SMA-positive cells), collagen degradation (by F4/80-positive Mph), and biomechanical forces generated by proliferating Heps. This regulation is captured by the gyration diameter, which quantifies the spatial spread of collagen fibers. Initially, this spread aligns with the developing fibrotic wall connecting neighboring CVs. The gyration diameter decreases from approximately 46 µm after the second CCl_4_ injection to about 20 µm after six doses, reflecting the compaction and structural consolidation of the collagen network into a well-defined fibrotic wall (**Fig. 4C, D**). This intricate interplay gives rise to the characteristic fibrotic patterns observed experimentally (**Fig. 4E**; **Supplementary Movie 2**).

**Figure 4.**
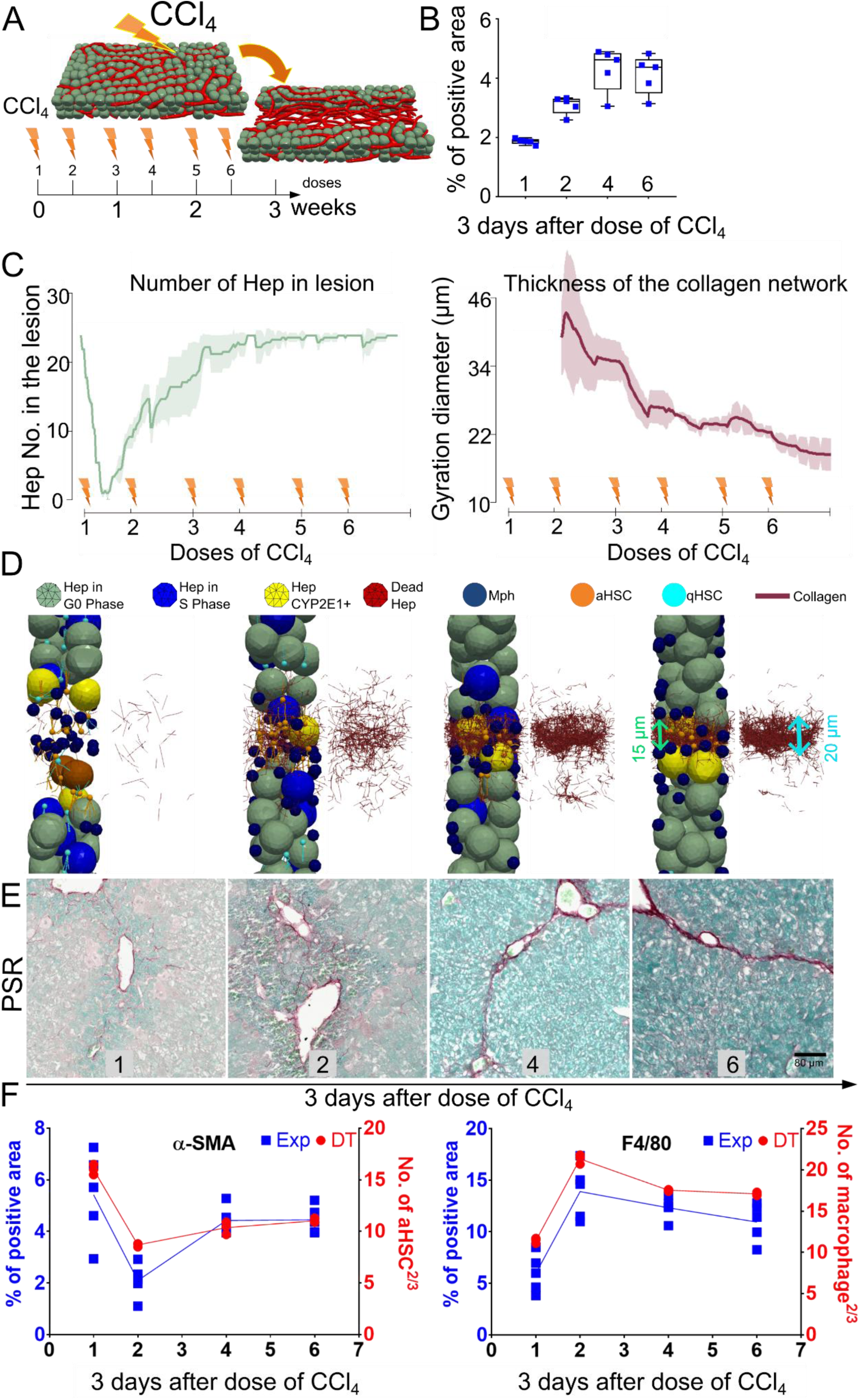
DT reference simulation predicts formation of wall-pattern of liver fibrosis upon repetitive doses of CCl_4_. (A) We run the simulation of repetitive doses of CCl_4_ on the DT cross-lobule system for 3 biological weeks. (B) The experimentally determined PSR positive area, resulting from different doses of CCl_4_, as indicated. (C) Number of Heps in the lesion and thickness of the collagen network over time. Thickness is measured by the gyration diameter of the collagen network corresponding to the standard deviation of the collagen distribution. The shaded area depicts the standard deviation of three independent simulations. (D) Snapshot of DT components (in the striped regions as shown in Fig. 1E; **Supplementary Movie 2**). The width of the collagen network buried inside the Heps (to mimic the PSR staining in experiment, spanned by green arrows, about 15 µm) and width of the concentrated part of the collagen network (spanned by cyan arrows, about 20 µm) are displayed. (E) PSR staining of collagen fibers in the fibrotic mouse liver at repeated doses of CCl_4_, as indicated (see details in Materials and Methods section, scale bars are 80μm). The size of the fibrotic wall after 6 doses of CCl_4_ is about 15μm. (F) Quantification of α-SMA and F4/80 positive areas, and the numbers of aHSCs and Mphs in the simulated lesion volume to the power of 2/3 (to compare the simulation result to the experimentally analyzed area *A* ∼ *V^2^*^/*3*^) as predicted from the DT. Experimental data are shown as the means ± SD of 5 mice per group.

#### C) Repeated rounds of CCl_4_-injection in the experiments support DT-predicted scenario

In a simulation replicating chronic liver injury over a 3-week period—with two CCl_4_ injections per week—each injection induces cell death of CYP2E1-expressing Heps localized within the lesion (**Supplementary Fig. 7A**). PSR staining shows that individual collagen fibers begin to appear by day 3 following the first CCl_4_ dose, gradually accumulating within the necrotic area after two doses, and coalescing into thick collagen bundles after four doses (**Fig. 4E**). By six CCl_4_ doses, a distinct wall-like fibrotic structure forms, extending between and enclosing CVs. The DT model successfully recapitulates this sequential pattern of collagen fiber deposition, as well as the spatial-temporal distribution of collagen and aHSCs (**Fig. 4D, E**). aHSCs begin migrating into the lesion following the first injection and progressively accumulate in the region where the collagen wall eventually forms after six doses (**Supplementary Fig. 9A**). This accumulation is also reflected in the temporal increase of α-SMA-positive cell fractions, as quantified in **Fig. 4F**.

As indicated by HE staining (**Supplementary Fig. 7A**), cellular infiltration—characterized by cells with small nuclei—is observed within and around fibrotic areas. A subset of these infiltrating cells are Mphs, whose spatial distribution closely follows that of aHSCs and collagen, as demonstrated by F4/80 immunostaining. Accumulation of F4/80-positive Mphs in the fibrotic region is evident after two CCl_4_ injections. By the fourth dose, Mphs begin to organize into wall-like structures, which by the sixth dose appear as walls linking neighboring CVs (**Supplementary Fig. 9A**). Quantification of F4/80-positive cells indicates that Mph numbers remain relatively stable following the second CCl_4_ dose (**Fig. 4F**). This apparent stabilization is further supported by RNA expression data, which show no additional upregulation of *Cd68*, *Mmp9*, or *Timp1* transcripts (**Supplementary Fig. 9D**).

In the DT model, Mphs are guided by the gradient of DAMPs released from CYP2E1-expressing Heps following the CCl_4_ injection. These Mphs migrate toward the lesion and remain distributed in a scattered pattern around it, even after six doses (**Fig. 4D**). The number of Mphs within the lesion is defined by the fraction of F4/80-positive cells, which increases after the first CCl_4_ dose (**Fig. 4F**).

In summary, the DT accurately recapitulates the spatial-temporal distribution patterns of collagen deposition, aHSCs, and Mphs, demonstrating concordance with experimental data. Model simulations predict that a gradient of DAMPs, released from CYP2E1-expressing Heps, orchestrates the spatial localization of collagen deposition around and between adjacent CVs.

### (VI) Spatial distribution of cells expressing CYP2E1 drives collagen deposition through the release of DAMPs, while the division of Heps shapes the resulting pattern of collagen deposition

To elucidate the origin of the observed collagen deposition patterns, CYP2E1-expressing Heps are identified as the spatial-temporal source of DAMPs. In cases of acute liver injury induced by CCl_4_, the number of CVs connected by CYP2E1-positive Heps increases from day 3 post-injection to an average of two connections per CV, and remains stable until at least day 16 (**Fig. 2C**; **Supplementary Fig. 3B**). Notably, under repetitive injury conditions, the majority of CVs exhibit 3–4 such connections after six doses—representing a marked increase compared to just 3 days (**Supplementary Fig. 7D**) and even 4 days (**Fig. 2C**) after the first CCl_4_ injection. This spatial pattern of CYP2E1+ Heps is incorporated into the DT model as the source of DAMPs for subsequent CCl_4_ injections (**Supplementary Figs. 3A, 8A**). Within this framework, the model predicts that the resulting spatial DAMP gradients drive collagen deposition by recruiting aHSCs and Mphs into the lesion site (**Supplementary Figs. 3A, 4A, 8B**; **Supplementary Movie 3**).

To investigate whether the spatial distribution of dying or damaged Heps influences the morphology of the resulting collagen network, we performed perturbation simulations (**Fig. 5A**). In one scenario (**Perturbation 1; P1, Fig. 5A**), we simulated a randomized distribution of CYP2E1+ Heps after the second CCl_4_ injection, in contrast to the reference case where CYP2E1 expression is restricted to the lesion area. This perturbation led to the formation of a collagen network with a broader spatial extent and a more homogeneous fiber distribution, lacking the characteristic wall-like structure observed in the reference case (**Fig. 5B, C**; **Supplementary Movie 4**).

**Figure 5.**
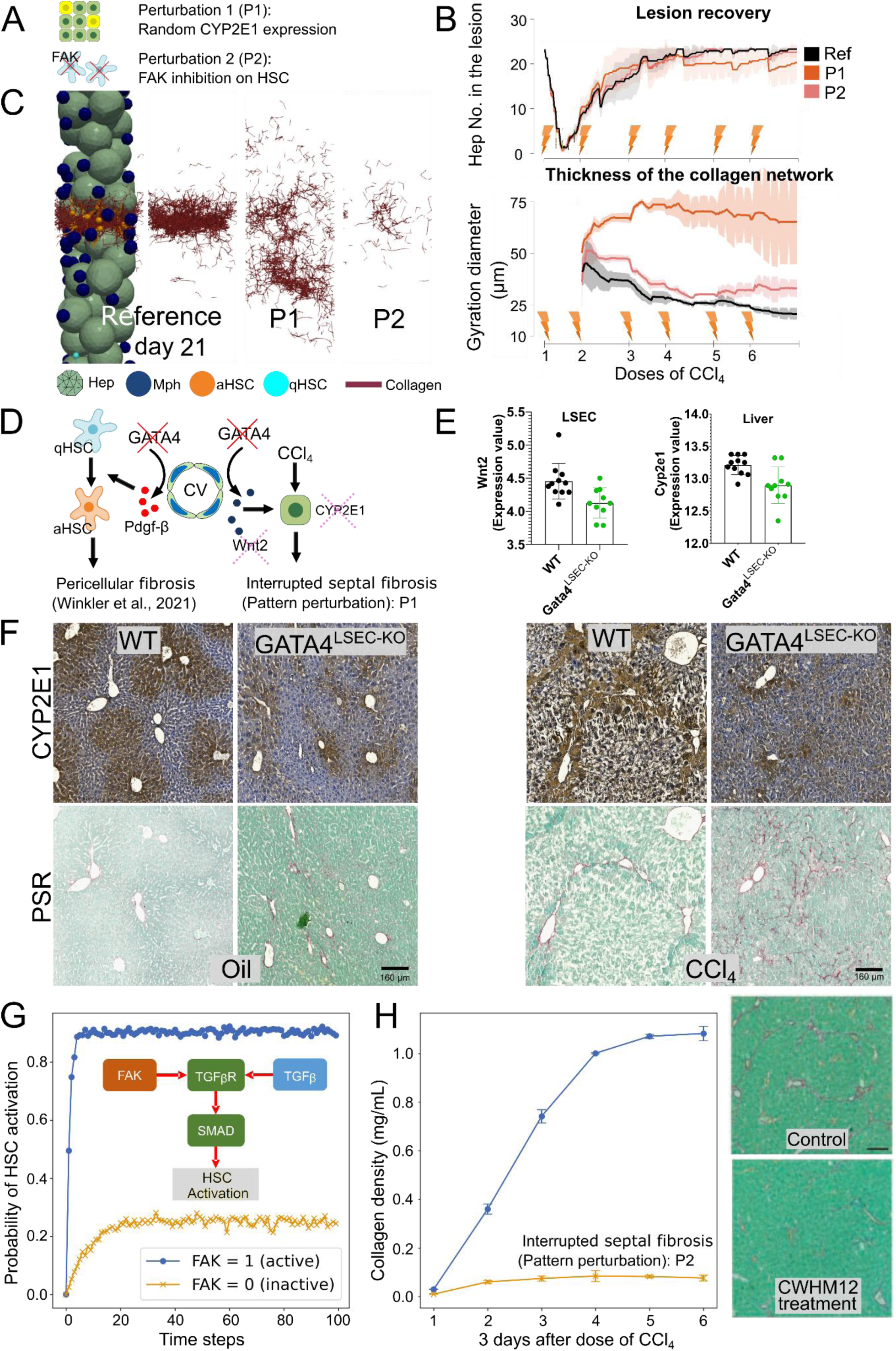
Perturbation simulations of random distributed CYP2E1 distribution and FAK inhibition and corresponding experimental validation. (A) Two different perturbation scenarios are tested. P1, randomly distributed CYP2E1 expressing Heps; P2, Inhibition of FAK activity on HSCs. (B) Number of Heps in the lesion and thickness of the collagen network during 3 weeks of chronic CCl_4_ injections. Thickness is determined by the gyration diameter of the network. The error bars represent the standard deviation of 3 simulation runs. (C) Snapshots of the collagen fiber network, resulting from the different perturbations, after 6 CCl_4_ injections each (highlighted is the section as shown in Fig. 1E). (The corresponding time-course simulations are shown in Supplementary Movies 4 and 5). (D) Endothelial cell specific deletion of the transcription factor GATA4 (Winkler et al.^34^) leads to decreased availability of the WNT2 ligand and a disturbed pattern of CYP2E1+ Heps. The model is used to experimentally validate perturbation scenario 1 (random CYP2E1 expression). (E) mRNA expression data (GSE141004) in endothelial cells isolated from GATA4^LSEC-KO^ mice, showing downregulation of WNT2. (F) CYP2E1 and PSR staining of wild type (WT) and GATA4^LSEC-KO^ mouse livers after 6 CCl_4_ injections. Data are shown as the means ± SD of 6-8 mice per group. (G) The intracellular network of FAK-promoted HSC activation and the Boolean network model to predict the long-term probability of HSC activation under the cases of active and inactive FAK expression. (H) PSR staining of control and CWHM12 treatment (inhibition of integrin) and the curve of collagen density over doses of CCl_4_ injection in DT simulation. Copyright of the experimental PSR staining of control and CWHM12 treatment: ^©^Henderson et al.^39^.

To experimentally validate the DT simulation for P1, we tested mice with altered spatial and absolute expression of CYP2E1. This was achieved by deleting a transcription factor GATA4 from endothelial cells^35^, resulting in a GATA4^LSEC-KO^ mouse model. In these mice, GATA4 deletion from endothelial cells leads to a decrease in Wnt2 expression, a well-known upstream regulator of CYP2E1 expression^36^. Consequently, these mice exhibit a reduced fraction of CYP2E1-positive Heps, attributed to the reduced Wnt2 expression from LSECs (**Fig. 5D, E**). The mice were subjected to repetitive CCl_4_ intoxications, receiving two injections per week over three consecutive weeks. As expected, the CYP2E1+ Hep distribution in the chronic CCl_4_-treated GATA4^LSEC-KO^ mice deviates from the control pattern, becoming more randomly distributed (**Fig. 5F**). Furthermore, collagen deposition was compromised in 75% (6 out of 8) of the GATA4^LSEC-KO^ mice, showing a more uniform distribution of perisinusoidal fibrosis compared to the wild type mice, which exhibited a characteristic septal fibrosis pattern (**Fig. 5F**). This perturbation experiment confirms that the spatial arrangement of CYP2E1+ Heps plays a key role in the development of septal fibrosis patterns in the liver following chronic CCl_4_ treatment.

### (VII) Test the effect of drug treatment on the activity of FAK on HSC

As a further challenge for the DT we conducted a perturbation test in the DT to predict the outcome of drug treatment on the activity of focal adhesion kinase (FAK) in HSCs. According to the study by Chen et al.^37^, FAK promotes TGFβ/TGFβ receptor-induced SMAD phosphorylation, which triggers HSC activation. To model this process, we constructed a simplified intracellular network of FAK-mediated HSC activation (**Fig. 5G**). Then, we applied a Boolean network model^38^ to estimate the long-term probability of HSC activation under conditions of active and inactive FAK expression. As shown in **Fig. 5G**, inhibiting FAK in HSCs reduces the probability of activation from 90% to 20% for the chosen parameters (up-/down-regulation rate, see Supplementary Information). We then integrated this Boolean network model into the DT to simulate fibrosis progression under drug treatment inhibiting FAK in HSCs. As shown in the collagen density curve (**Fig. 5H**), collagen expression was significantly reduced when FAK activity was suppressed upon treatment. This prediction aligns with experimental findings by Henderson et al.^39^, who demonstrated that inhibiting integrin (a binding partner of FAK) activity in HSCs during CCl_4_-induced fibrosis progression reduced collagen expression. Our findings underscore the utility of the DT model as a platform for virtual drug testing and hypothesis generation for antifibrotic targets.

### (VIII) Validation of DT model by simulating the pattern formation of biliary fibrosis

To further demonstrate the applicability of the DT, we applied the DT to explain the mechanism underlying the development of biliary fibrosis patterns. We analyzed the distribution of bile ducts (Fibrogenic niche)^40^ in Abcb4*KO* mice, a well-established model for biliary fibrosis, at age 8, 16, and 28 weeks. Using this data, we generated a heatmap of the spatial-temporal distribution of bile ducts along the PV-PV region (**Fig. 6A and B**). The DT was then set up in the region between two PVs, and bile ducts were generated based on the heatmap (**Fig. 6C**). Bile leakage from the bile ducts induces cell death in surrounding Heps, leading to the release of DAMPs, which attract HSCs and Mphs to produce and degrade collagen (similar to septal fibrosis). The same number of Heps proliferate from adjacent healthy cells to replace the dying ones (for details see the SI). As shown in **Fig. 6D**, the simulation results for collagen distribution align with experimental observations in Abcb4*KO* mice, where collagen is more broadly and randomly distributed along the PV-PV axes in biliary fibrosis. Unlike septal fibrosis, where collagen tends to finally form "wall-like" thin bundles, the collagen in biliary fibrosis surrounds Heps in a more dispersed pattern. The very low rate of Hep division in biliary fibrosis (approximately 4%) may contribute to the broader fibrotic pattern observed due to loss of biomechanical compression from Hep proliferation in septal fibrosis (see **Supplementary Fig. 8D**). Interestingly, if such bile ducts along the PV-PV axis do not form, a layer-by-layer “onion”-like pattern forms as observed in onion-skin biliary fibrosis in for example human Primary Sclerosing Cholangitis (**Fig. 6F**). Here, collagen secreted by aHSCs is initially pressed into inter-hepatic spaces, subsequently increasingly replacing dead hepatocytes layer-by-layer.

**Figure 6.**
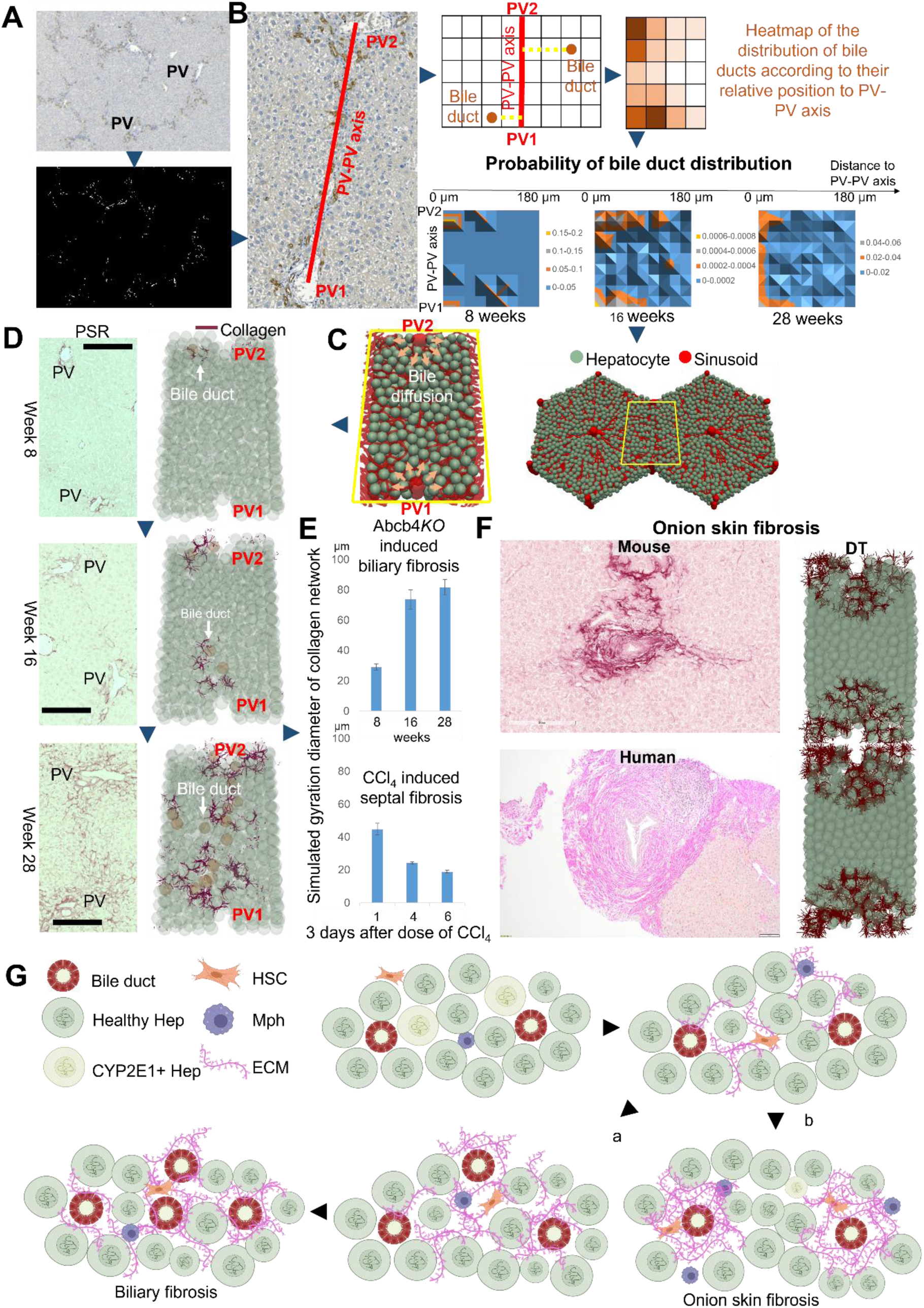
Simulation of the biliary fibrosis and experimental validation. (A) Visualization and quantification of bile duct (CK19, brown in the top figure) in Abcb4*KO* mouse liver. The binary of the original staining locates the coordinates of the bile ducts (white in the bottom figure). (B) Each pair of PV-PV is taken out and one line is added to represent the PV-PV axis (as shown in red). The heatmap for three different time points (8, 16, and 28 weeks old mice) were generated (see more details in the **SI**). (C) A DT in a section between two PVs was set up to simulate the process of biliary fibrosis formation. The bile ducts were assumed to induce the cell death of neighboring Heps. The spatial-temporal distribution of bile ducts (represented as brown spheres pointed by white arrows) was according to the heat maps obtained in B. (D) The simulation results of the distribution of collagens (right, the width of the section: 240 µm) and compared to the experimental data (left, PSR, scale bar: 200 µm) during the progression of biliary fibrosis. (E) The simulation results of gyration diameter of the collagen network of both biliary and septal fibrosis. (F) In case no bile ducts form along the PV-PV axis, the collagen forms rings around the portal tract as observed in mouse and human in biliary fibrosis for example in Primary Sclerosing Cholangitis. The DT displays the onion-like pattern. Shown are DT simulations with smaller (upper picture) and larger (lower picture) collagen production rates. (G) Inferred mechanism of formation of biliary fibrosis pattern in (D), and in onion skin fibrosis pattern (F): the bile leaking out from the duct induces the death of surrounding Heps. The dead Heps attract aHSCs and Mphs to generate and digest collagen fibers. The collagen fibers wrap Heps to form a chicken-wire pattern. As more bile ducts are generated in a broad area (a), more collagen fibers are produced. Since there is less mechanical pressure due to minimal Hep proliferation, the distribution of collagen fibers is much more scattered. Otherwise, if no more bile ducts are generated in the region between PVs (b), collagen fibers form layers surrounding the bile ducts like onion skins.

In conclusion, the fibrosis pattern is primarily driven by the spatial distribution of Hep death, HCS activation and migration, and mechanical pressure resulting from the Heps. In septal fibrosis, the distribution of dead Heps is determined by the location of CYP2E1+ Heps, in biliary fibrosis, by the localisation of leaking bile ducts (Fig. 6E).

## Discussion

Liver fibrosis is characterized by the accumulation of ECM, which in the CCl_4_ mouse model forms fibrotic walls linking the CVs of adjacent lobules due to repeated CCl_4_ injections. A key concept in understanding liver fibrosis is determining why and where ECM is deposited in specific patterns, which ultimately progress to cirrhotic nodules (a major risk factor for developing HCC). Characteristic patterns include fibrotic walls, observed in septal fibrosis, or more dispersed collagen patterns, such as those observed in biliary fibrosis. In this work, we study the septal fibrotic pattern linking the CVs of adjacent lobules in response to chronic CCl₄ exposure, comparing it to the pattern observed in biliary fibrosis. However, a comprehensive understanding of these patterns has been hindered by their inherent complexity, involving numerous components and subprocesses that cannot currently be simultaneously monitored through experimental approaches.

To complement single- or dual-target experimental approaches, we have developed a DT of a cross-lobule liver system. This DT model represents the fundamental cellular and structural unit underlying the liver fibrosis pattern. It is built upon quantifiable experimental parameters to explore the potential sequence and interplay of subprocesses involved in fibrotic wall formation. The iterative process includes refining the DT model based on experimental data, making predictions, and validation through further animal experimentation.

Previous computational models have been applied to predict cellular and tissue responses of both acute and chronic liver injury in mice and patients, integrating complex interactions and system components^41-48^. Some models resolve tissue microarchitecture by representing individual cells^41,43,44,46-48^. Some continuum models study spatial cell densities and molecular concentrations where the individual organization of cells can be neglected^45^. Whereas others use statistical models that disregard spatial structure entirely^42^. At the tissue microarchitectural level, some models employ a rigid scaffold structure leading to cellular automaton models having short computational time. However, gradual cell displacements or deformations are not allowed by these models^43,44,46^. Other models can closely mimic 3D histological reconstructions by resolving biomechanical properties of cells and tissues. However, the computational time is long^41,47,48^.

The DT presented in this work belongs to the latter class of models, encompassing a minimal selection of critical liver cell types (Hep, HSC, and Mph) as well as key tissue elements (sinusoidal networks and collagen fibers). It serves as a platform to study the development, progression, and pattern formation of septal fibrosis following CCl_4_ exposure and biliary fibrosis induced by deleting the Abcb4 gene. The DT also provides insights into how mechanical interactions and molecular crosstalk between these tissue components contribute to the observed septal/biliary fibrosis pattern in experiments. To achieve this, the DT captures the spatial-temporal dynamics of tissue microarchitecture. Comparing to earlier liver DTs addressing other pathologies (**Supplementary Fig. 1**), this DT directly simulates cells and the collagen network at subcellular resolution, allowing for the modeling of their rearrangements and remodeling in response to repetitive CCl_4_ exposure and bile leakage from biliary ducts. Modeling at subcellular resolution introduces substantial technical and computational complexity, which is crucial to ensure accuracy and to minimize the risk of introducing artifacts into the simulation.

To demonstrate the interplay of intracellular mechanisms governing cellular decisions and cell-cell communication we studied CWHM12-treatment of septal fibrosis. However, at the current stage of the DT model, the most intricate intracellular mechanisms governing cellular decisions and cell-cell communication have not yet been explicitly incorporated but are planned as follow-up steps in the integration of molecular and cellular scale models. A simulation can be viewed as the *in-silico* equivalent of observing a biological process under a microscope—offering a controlled, well-defined environment similar to intravital imaging. However, intravital imaging of the fibrotic wall formation process remains out of reach due to the extended duration of this process. Repetitive CCl_4_ exposure leads to cycles of tissue damage of CYP2E1-positive Heps, followed by incomplete regeneration through the division of surviving Heps. In the DT model, the region of CYP2E1 positivity is determined based on experimental data linking adjacent CVs from the second CCl_4_ injection onward.

The dynamic process of Hep growth and division is stochastically sampled from the spatial-temporal distribution of Ki-67-positive cells surrounding the lesion over time. The migration of HSCs and Mphs is modeled by applying a chemotactic force that directs cells along the gradient of DAMPs^49,50^. We model active cell migration following the approach outlined in a previous study^29^, omitting explicit representation of cell protrusions, as it has been shown that these do not alter the overall movement dynamics^51^.

DAMPs are released by damaged Heps. After the initial exposure to CCl_4_, DAMPs concentrate around CVs. However, during subsequent regeneration phases, observations align with previous studies^52,53^, showing that CYP2E1-expressing cells recover and form a wedge-shaped structure connecting adjacent CVs. Analysis of the connections between CYP2E1-expressing Heps and adjacent CVs reveals that shortly after fibrosis establishment, one CV is linked to 3-4 neighboring CVs. This occurs when a “mother” CV associated with one lobule splits into two “daughter” CVs, which are then associated with two other lobules. Before branching, there is one circular CYP2E1-positive area surrounding the mother CV, and each daughter CV is similarly associated with a circular CYP2E1-positive area. Through the tree-like hierarchical branching structure, the CYP2E1-positive zones of both daughter CVs remain connected until they are sufficiently distant from each other (the branching concept is explained in **Figs. 3 and 4** in ref^54^). Consequently, Heps die along these CV-CV connections after subsequent CCl_4_ doses, altering the spatial pattern of DAMP sources from around the CV after the first dose to along the CV-CV connections in later doses. Therefore, the time point of the second dose, three days after the first CCl_4_ exposure, is selected as the model’s reference point. This time point includes critical fibrogenic factors such as the earliest signs of metabolic activity in midzonal Heps, pronounced cell division, maximal HSC activation, and Mph infiltration^29,55^. Most model parameters are calibrated based on data collected at this time point. Using the calibrated DT, simulations are performed for six consecutive CCl_4_ injections, with the results compared to experimental outcomes.

The DT simulations faithfully reproduce the experimentally observed scenario, including the precise pattern formation of fibrotic walls. This supports the hypothesis that the spatial distribution of CYP2E1-expressing Heps drives fibrosis patterning through DAMP gradients released by damaged cells. While spatial distribution of CYP2E1+ Heps explain where fibrosis occurs, it does not explain the sharp localization of fibrotic walls between two adjacent CVs. The DT suggests that this focal consolidation results from successive waves of Hep proliferation following repeated CCl_4_ administrations. As necrotic lesions resolve, proliferating Heps in the surrounding area exert biomechanical pressure, displacing the collagen network. When collagen density is still low, the model predicts that the collagen network is pressed into inter-cellular spaces, forming a characteristic “chicken-wire” pattern—an observation confirmed by careful analysis of experimental images (**Supplementary Fig. 8C**). However, after 3–4 rounds of CCl_4_ administrations, when collagen density is high, the DT predicts that the collagen network is shifted towards the center of the necrotic zone. This prediction is supported by the experimental findings showing that initially, collagen fibers are diffusely distributed within the CYP2E1+ zone deposited by activated HSCs, but over time, the collagen network consolidates into a sharply localized fibrotic wall.

To test this concept, we performed a mathematical analysis using the DT by simulating a homogeneous distribution of CYP2E1-expressing Heps. Under this condition, the DT predicted a dispersed DAMP concentration, which resulted in homogeneous collagen deposition rather than the sharply localized fibrotic walls observed in the reference scenario. This hypothesis was experimentally validated using GATA4^LSEC-KO^ mice, in which deletion of the GATA4 transcription factor—essential for LSEC differentiation—leads to a disrupted and more scattered expression of CYP2E1. Upon repeated CCl_4_ administration, these mice exhibited a correspondingly scattered fibrosis pattern, lacking the characteristic fibrotic walls observed in the reference case, which was in agreement with DT predictions. These findings highlight that the spatial distribution of dead Heps, as the primary DAMP source in chronic liver injury, plays a critical role in influencing the overall fibrosis pattern.

To further support the role of biomechanical pressure in the localization of the collagen network, we used the DT to simulate biliary fibrosis based on experimental data from Abcb4*KO* mice. In these mice, knockout of the Abcb4 gene leads to the formation of numerous bile ducts along the PV–PV axis. Leaking bile induces localized Hep death, which in turn activates HSCs to produce collagen fibers. The simulation reveals that, unlike the sharply defined fibrotic walls observed in septal fibrosis, collagen fibers in biliary pattern form a much more scattered network. This difference arises from two key factors: (1) In biliary fibrosis, the regions of Hep death are less spatially connected, leading to a more diffuse pattern of collagen deposition—similar to the scattered fibrosis pattern seen in GATA4^LSEC-KO^ mice. This reinforces the idea that the shape and connectivity of the dying Hep regions critically influence fibrosis patterns. (2) The extent of Hep death and proliferation is significantly lower in biliary fibrosis compared to the CCl_4_-induced fibrosis. Consequently, the mechanical displacement of collagen fibers by proliferating Heps, which transiently generates the chicken-wire pattern at low collagen densities in septal fibrosis (as seen at days 7 or 10 in **Supplementary Fig. 8E**), is less pronounced. In biliary fibrosis, the absence of large, connected regions of Hep death allows the chicken-wire pattern to persist over extended periods (**Fig. 6F**).

However, our DT simulations suggest that a chicken-wire pattern should transiently emerge in other liver damage contexts as well, not limited to those explicitly studied in this work. Supporting this statement, such patterns are also observed in steatotic and alcohol-related liver diseases (**Supplementary Fig. 8F**) in human livers. These findings highlight the model’s adaptability across different disease etiologies and reinforce its potential clinical relevance.

In summary, the DT presented here successfully explains the formation of fibrotic walls in septal fibrosis and the scattered collagen distribution characteristic of biliary fibrosis. While previous studies have addressed specific aspects of fibrosis development, they have often neglected the role of tissue microarchitecture and the contribution of mechanical forces—both of which are central to the DT model’s predictive power.

Friedman and Hao^45^ developed a mathematical model to investigate liver fibrosis by representing interactions among various cell types and cytokines. However, their model adopts a continuum approach that averages cellular behavior over multiple cell diameters, thereby lacking the spatial resolution necessary to capture tissue microarchitecture and permit direct comparison to histology. Furthermore, their focus is restricted to cellular and molecular kinetics, omitting mechanical components entirely. In contrast, Dutta-Moscato et al.^43^ and Yoshizawa et al.^44^ utilized agent-based models (ABMs) to simulate aspects of fibrotic wall formation. Both approaches position cells on a fixed two-dimensional lattice and entirely omit the sinusoidal network. However, implementing a space-fixed lattice restricts the ability to simulate realistic cell and tissue biomechanics, as demonstrated by Van Liedekerke et al.^51^. This omission is particularly problematic since biomechanical interactions are essential to fibrotic wall formation, as shown in the current DT model. Additionally, the absence of the sinusoidal network—an essential scaffold for cell movement and division—is a critical limitation. Drasdo and Zhao^25^ and Hoehme et al.^29^ have shown that omitting this network can fundamentally alter simulated disease dynamics.

Moreover, both Dutta-Moscato et al.^43^ and Yoshizawa et al.^44^ model septa between liver lobules and compare their simulation results with rodent (rat) data, where connective tissue septa are not observed under physiologically normal conditions in rodents. Their models are rule-based, with parameters expressed as unitless numerical values. This approach raises concerns about realism of simulated dynamics, as many parameters cannot be directly linked to measurable physical or biological quantities. This makes it difficult to know whether they are within physiological parameter ranges. Some modeling assumptions appear biologically implausible—for instance, in Dutta-Moscato et al.^43^, Heps can die and proliferate but are not allowed to move, which contradicts well-established tissue biomechanics.

In addition, the model proposed by Dutta-Moscato et al.^43^ predicts the formation of fibrotic walls along the portal field, which contradicts experimental observations^56^. It does not correspond to the spatial distribution of CYP2E1-positive Heps known to drive fibrosis in the CCl_4_ model. Consequently, their model identifies fibrotic structures at locations where such formations are not observed in rodent livers following CCl_4_ intoxication. In contrast, the current DT demonstrates that fibrotic walls consistently form between CVs, which is in agreement with experimental data. While Yoshizawa et al.^44^ correctly interpreted histological images suggesting fibrotic walls between CVs, their model fails to reproduce this pattern, despite incorporating spatial-temporal dynamics. This suggests that the underlying mechanisms implemented, or their technical execution, are likely incomplete.

The main differences between the simulation outcomes of the ABM models by Dutta-Moscato et al.^43^ and Yoshizawa et al.^44^ and our proposed DT model are due to two factors: the biological realism of the model and the spatial dimensions used. The current DT model works in a fully 3D spatial setting and includes key structural elements, like sinusoids and biomechanical interactions, which are crucial for accurately simulating fibrosis patterns.

We believe the present mathematical model for septal fibrosis marks a substantial advancement toward a comprehensive DT of the CLD. Furthermore, the DT’s architecture supports the progressive integration of additional cell types and molecular processes, ultimately enabling simulations of disease progression toward decompensated cirrhosis and hepatocellular carcinoma—key endpoints in CLD that go beyond the fibrosis stage.

The DT approach offers a wide range of opportunities for both basic and translational research applications as indicated by the simulated examples of septal fibrosis (**Figs. 1-4**), and fibrosis development in Abcb4*KO* mice and human (**Fig. 6**). As demonstrated by Drasdo and Zhao^25^, it can be used to guide the selection and design of preclinical experiments including drug testing (**Fig. 5**), by enabling hypothesis testing within a rigorously defined *in silico* setting. The DT allows investigation of complex interactions across multiple biological scales—from intracellular signaling pathways to tissue-level organization and even organism-wide dynamics, and holds the potential to extrapolate findings from in vitro experiments to predict in vivo behavior as illustrated by Dichamp et al.^47^. Moreover, as shown in the study on resected liver by Hoehme et al.^41^, the DT permits extrapolation across species boundaries, and, as demonstrated for biliary fibrosis in this work (**Fig. 6**), to identify possibly critical mechanistic differences in disease progression between species. This predictive capacity underscores the DT’s potential as a powerful tool in both mechanistic research and preclinical modeling.

Although the DT has primarily been applied in non-human settings, its potential translation to human contexts holds significant promise. It could help bridge the gap between human tissue cultures, bioengineered systems, animal disease models, and patient-specific scenarios. Initial setbacks are to be expected. However, these challenges are valuable for uncovering critical gaps in current knowledge regarding underlying mechanisms and parameterization, ultimately advancing a more comprehensive understanding of human liver disease and fostering drug testing.

Personalizing the DT for individual patients remains a major challenge. This involves extrapolating in vivo dynamics from in vitro systems, such as patient-derived cells, organoids, or slices. Additionally, findings from murine models need to be translated to human pathophysiology, and the DT could also be directly applied in clinical environments. By integrating molecular data collected at various disease stages—particularly biomarkers identified through transcriptomics, proteomics, or histopathology^57^— the DT could serve as a complementary tool alongside emerging high-resolution imaging technologies, such as high-field magnetic resonance imaging (MRI) or temporal diffusion spectroscopy.

While artificial intelligence (AI) approaches offer powerful tools for pattern recognition and data-driven predictions, mechanistic DTs provide unique predictive capabilities for novel therapeutic interventions and uncover causal relationships where empirical data is lacking. Rather than being mutually exclusive, AI and mechanistic modeling approaches are inherently complementary.

In conclusion, the presented DT offers a robust and mechanistically grounded framework for understanding fibrotic wall formation in septal and biliary fibrosis. It represents a substantial advancement in the modeling of liver disease, enabling the integration of experimental observations with spatial and biomechanical dynamics. This DT paves the way for broader applications across diverse etiologies and disease stages, ultimately fostering a deeper, more predictive understanding of CLD and supporting translational efforts from bench to bedside.

### Limitation

Mphs produce matrix metalloproteinases (MMPs) to digest the collagen fibers. They further have been reported to contribute to the survival of HSCs^58,59^, and they phagocytize dead cells^7^. We here focus on the phagocytizing function of Mphs and do not consider a survival signal to aHSC provided by Mphs or other immune cells. In our DT, aHSC autonomously does not revert or die before either the lesion is closed, or a certain collagen density is locally reached. Further refinements such as including the possible influence of Mphs on aHSC survival will be addressed in future versions of the DT along with further applications in disease progression, as e.g. until HCC.

## Methods

### Animal experiments

To investigate septal fibrosis patterns, adult C57Bl/6N male mice (Janvier labs; France) are housed three per cage in a temperature-controlled (24°C) room with a 12-h light/dark cycle. Mice are given *ad libitum* access to water and laboratory diet (Ssniff, Germany). Mice are exposed to carbon tetrachloride (CCl_4_; Sigma-Aldrich, Cat. no. 319961) in a dose of 1.6 g/kg body weight^56,60^, either at a single (acute) or repeated doses. To track regeneration and wound healing parameters upon acute intoxication, mice are exposed to a single dose of CCl_4_. Blood and liver samples are harvested at 0, 1, 2, 3, 4, 6, 16, and 30 days post CCl_4_ injection. Liver fibrosis is induced by two doses of CCl_4_/week for 3 weeks^61^. Three days after the last CCl_4_ injection, blood and livers are harvested. To study biliary fibrosis patterns, Abcb4*KO* mice on a BALB/c background are maintained under pathogen-free conditions. Blood and liver samples from Abcb4*KO* mice for 8, 16 and 28 weeks^28^. All experimental protocols with animals are carried out in full compliance with the guidelines for animal care and are approved by the Animal Care Committee from the German government (Animal permission number: 35.9185.81/G87/10, 35.9185.81/G-216/16 and 35.9185.81/G-22/20).

#### Histological staining and quantification

Stainings are performed on paraffin slides (3µm) for hematoxylin and eosin (H&E) and Sirius red staining and with fast green staining^62^. Further, immunohistochemistry (IHC) using anti-CYP2E1 (Sigma-Aldrich, HPA009128, raised in rabbit, 1:50), anti-mouse F4/80 (BioRad, MCA497, raised in rat, dilution 1:2000), anti-Ki-67 (Cell Signaling Technology, CST#12202, raised in rabbit, dilution 1:200), anti-CK19 (ProteinTech, 14965-1-AP, raised in rabbit,1:100), anti-BrdU (BioRad, MCA2060, raised in rat, dilution 1:250) and anti-α-smooth muscle actin (α-SMA; Abcam, Ab5694; raised in rabbit, dilution 1:100) is performed. Image acquisition is performed using an Aperio 8 slide scanner (Leica). A minimum of 10 high-power fields is captured for image analysis. The positive areas are quantified as a brown area percentage or nuclei number by Image J or manually, respectively (5-7 fields at least per slide, 10X).

### Mathematical model: the digital twin

The digital twin considers each individual Hep, HSC, and Mph as individual model units, and considers sinusoids, and extracellular matrix bundles both in space and time. Each Hep and Mph is represented by one component, while HSCs, sinusoids, and ECM bundles are mimicked as composed of many model components. The movement and interaction of each model component is modeled by a force balance equation. The force balance equations are updated simultaneously for all model components (e.g. Hep, HSCs, Mphs and the collagen fibers) by solving a system of coupled equations, one for each component, to represent all forces exerted on each component: ∑ *F*_*fri*_ + *F*_*def*_ + *F*_*adh*_ = 0. Here *F*_*fri*_, *F*_*def*_, *F*_*adh*_are the friction force with the environment, deformation force, and adhesive force, respectively (see details in SI). In total, half a million equations (total number of model components) had to be solved at each time step and were simulated over three weeks of real time. Due to the long simulation times with one time step of the order of 10^-3^ s, we have to limit it to one single part of the damage region along the CV-CV axis of two neighboring lobules sections. In addition to direct interactions by physical forces, cells can communicate via diffusive signals. For these, reaction-diffusion equations are solved.

*Detailed Methods are described in the **Supplementary Information***.

## ♠ Abbreviations and definitions

Hepatocyte (Hep): The main functional cell type in the liver, responsible for metabolism, detoxification, and protein synthesis among other function
Sinusoids (Sin): Fenestrated liver capillaries at which the exchange of nutrients and waste occurs between blood and liver cells
Portal Vein (PV): The portal vein within the liver that supplies nutrients- containing blood
Central Vein (CV): The vein collects blood from sinusoids to drain into the hepatic vein
Kupffer Cell (KC): Specialized macrophages within the liver that help remove pathogens and debris
Macrophages (Mph): 
Pericentral Hepatocyte (PH): Hepatocytes located near the central vein, often with distinct metabolic and detoxification functions
Hepatic Stellate Cell (HSC): Liver cells that store vitamin A and contribute to fibrosis in liver injury
Extracellular Matrix (ECM): Network of proteins and molecules outside cells that provide structural support to the liver
Endothelial Cell (LSEC): Cells lining the blood vessels in the liver, including sinusoids, facilitating nutrient and waste exchange
Cytochrome p450 2E1 (CYP2E1) region: pericentral hepatocytes are able to metabolize the toxic substance i.e. CCl4 to the active form
Carbon tetrachloride (CCl4): toxic molecule to induce liver damage
Damage associated molecular pattern molecules (DAMPs): molecules expressed and released upon tissue damage, resulting in activation of the immune system
Picro-sirius red (PSR): staining marker for collagen fibers
GATA binding protein 4 (GATA4): protein regulating liver function such as to regulate Wnt expression
Knockout (KO): 
Knock-out the gene X in the liver sinusoidal endothelial cells (X^LSEC-KO^): 
Bile duct (BD): network of tubes responsible for transporting bile from the liver and gallbladder to the duodenum (the first part of the small intestine), where it aids in digestion, particularly in the emulsification and absorption of fats
ATP Binding Cassette Subfamily B Member 4 (Abcb4): Abcb4 transports phospholipids from hepatocytes to bile canaliculi and bile ducts, helping to dilute bile acids to physiological levels. Deletion of Abcb4 results in toxic bile acid accumulation, leading to cell injury. Abcb4*KO* mice, deficient in canalicular phospholipid flippase, spontaneously develop biliary fibrosis due to the absence of phospholipids in bile.

## Data availability

The data that support the findings of this study are available from the corresponding authors upon reasonable request.

## Code availability

The simulation source code is accessed upon request to upon signing a non-disclosure agreement since the final license of the current code has not been granted yet.

## Acknowledgements

The authors acknowledge the support of the Core Facility for LIMA Live Cell Imaging Mannheim at Microscopy Core Facility Platform Mannheim (CFPM), Medical Faculty Mannheim, Heidelberg University and the data storage service SDS@hd supported by the Ministry of Science, Research and the Arts Baden-Württemberg and the German Research Foundation.

## Author contributions

S.H conceived animal studies, blood analysis, and RNA isolation. P.E, S.H, B.H and Y.L performed RT-PCR, and IHC staining, as well as manual image analysis. M.W, S.W.K, P.S.K performed animal experiments in GATA4^LSEC-KO^ mice. M.D.L; B.H and N.G performed an automated image analysis. J.Z and D.D designed and developed the DT model. J.Z implemented the code and performed simulations and visualized the results. P.V.L and A.B provided the original code of DCM and collagen model. S.H and J.Z wrote the manuscript. M.P.E, J.G.H, S.D, S.D.W, J.B and D.D performed critical revision of the manuscript. S.D, S.H and D.D acquired funding for this study. All authors read the final version of the manuscript.

## References

1. Pinzani, M., Rombouts, K. Liver fibrosis: from the bench to clinical targets. Dig. Liver Dis. 36(4), 231–242 (2004).

2. Kisseleva, T., Brenner, D. Molecular and cellular mechanisms of liver fibrosis and its regression. Nat. Rev. Gastroenterol. Hepatol. 18(3), 151–166 (2021).

3. Moon, H., Cho, K., Shin, S., Kim, D.Y., Han, K., Ro, S.W. High risk of hepatocellular carcinoma development in fibrotic liver: role of the Hippo-YAP/TAZ signaling pathway. International Journal of Molecular Sciences 20, 581 (2019).

4. Seki, E., Schwabe, R.F. Hepatic inflammation and fibrosis: functional links and key pathways. Hepatology 61(3), 1066–1079 (2015).

5. Higashi, T., Friedman, S.L., Hoshida, Y. Hepatic stellate cells as key target in liver fibrosis. Adv Drug Deliv. Rev. 121, 27–42 (2017).

6. Ramachandran, P., Pellicoro, A., Vernon, M.A., Boulter, L., Aucott, R.L., Ali, A., Hartland, S.N., Snowdon, V.K., Cappon, A., Gordon-Walker, T.T., Williams, M.J., Dunbar, D.R., Manning, J.R., Van Rooijen, N., Fallowfield, J.A., Forbes, S.J., Iredale, J.P. Differential Ly-6C expression identifies the recruited macrophage phenotype, which orchestrates the regression of murine liver fibrosis. Proc Natl Acad Sci USA. 109(46), E3186–95 (2012).

7. Krenkel, O., Tacke, F. Liver macrophages in tissue homeostasis and disease. Nature Rev. Immunol. 17, 306–321 (2007).

8. Papachristoforou, E., Ramachandran, P. Macrophages as key regulators of liver health and disease. Int. Rev. Cell Mol. Biol. 368, 143–212 (2022).

9. Krenkel, O., Hundertmark, J., Ritz, T.P., Weiskirchen, R., Tacke, F. Single cell RNA sequencing identifies subsets of hepatic stellate cells and myofibroblasts in liver fibrosis. Cells 8(5), 503 (2019).

10. Kisseleva, T., Cong, M., Paik, Y., Scholten, D., Jiang, C., Benner, C., Iwaisako, K., Moore-Morris, T., Scott, B., Tsukamoto, H., Evans, S.M., Dillmann, W., Glass, C.K., Brenner, D.A. Myofibroblasts revert to an inactive phenotype during regression of liver fibrosis. Proc. Natl Acad Sci USA 109(24), 9448–53 (2012).

11. Filliol, A., Saito, Y., Nair, A., Dapito, D.H., Yu, L., Ravichandra, A., Bhattacharjee, S., Affo, S., Fujiwara, N., Su, H., Sun, Q., Savage, T.M., Wilson-Kanamori, J.R., Caviglia, J.M., Chin, L., Chen, D., Wang, X., Caruso, S., Kang, J.K., Amin, A.D., Wallace, S., Dobie, R., Yin, D., Rodriguez-Fiallos, O.M., Yin, C., Mehal, A., Izar, B., Friedman, R.A., Wells, R.G., Pajvani, U.B., Hoshida, Y., Remotti, H.E., Arpaia, N., Zucman-Rossi, J., Karin, M., Henderson, N.C., Tabas, I., Schwabe, R.F. Opposing roles of hepatic stellate cell subpopulations in hepatocarcinogenesis. Nature 610(7931): 356-365 (2022).

12. Huang, H., Chang, M., Chen, Y., Hsu, H., Chiang, C., Cheng, T., Wu, Y., Wu, M.Z., Hsu, Y., Shen, C., Lee, C., Chuang, Y., Hong, C., Jeng, Y., Chen, P., Chen, H., Lee, M. Persistent elevation of hepatocyte growth factor activator inhibitors in cholangiopathies affects liver fibrosis and differentiation. Hepatology 55(1), 161–172 (2012).

13. Török, N., Fan, W., Adebowale, K., Li, Y., Rabbi, M.F., Vancza, L., Chen, D., Kunimoto, K., Mozes, G., Li, Y., Tao, J., Monga, S., Charville, G., Wells, R., Dhanasekaran, R., Kim, T., Chaudhuri, O. Extracellular matrix viscoelasticity drives liver cancer progression in pre-cirrhotic NASH. preprint (2022), available at Research Square [10.21203/rs.3.rs-2087090/v1]

14. Caliari, S.R., Perepelyuk, M., Cosgrove, B.D., Tsai, S.J., Lee, G.Y., Mauck, R.L., Wells, R.G., Burdick, J.A. Stiffening hydrogels for investigating the dynamics of hepatic stellate cell mechanotransduction during myofibroblast activation. Sci. Rep. 6, 21387 (2016).

15. Hernandez-Gea, V., Friedman, S.L. Pathogenesis of liver fibrosis. Annu. Rev. Pathol. 6, 425–456 (2011).

16. Parola, M., Pinzani, M. Liver fibrosis: pathophysiology, pathogenetic targets and clinical issues. Molecular Aspects of Medicine 65, 37–55 (2019).

17. Pinzani, M., Luong, T.V. Pathogenesis of biliary fibrosis. Biochimica et Biophysica Acta - Molecular Basis of Disease 1864 (4), 1279–1283 (2018).

18. Häussinger, D. Hepatocyte heterogeneity in glutamine and ammonia metabolism and the role of an intercellular glutamine cycle during ureogenesis in perfused rat liver. European Journal of Biochemistry 133(2), 269–275 (1983).

19. Gebhardt, R., Mecke, D. Heterogeneous distribution of glutamine synthetase among rat liver parenchymal cells in situ and in primary culture. The EMBO Journal 2(4), 567–570 (1983).

20. Schliess, F., Hoehme, S., Henkel, S.G., Ghallab, A., Driesch, D., Bottger, J., Guthke, R., Pfaff, M., Hengstler, J.G., Gebhardt, R., Häussinger, D., Drasdo, D., Zellmer, S. Integrated metabolic spatial-temporal model for the prediction of ammonia detoxification during liver damage and regeneration. Hepatology 60(6), 2040–2051 (2014).

21. Ghallab, A., Celliere, G., Henkel, S.G., Driesch, D., Hoehme, S., Hofmann, U., Zellmer, S., Godoy, P., Sachinidis, A., Blaszkewicz, M., Reif, R., Marchan, R., Kuepfer, L., Häussinger, D., Drasdo, D., Gebhardt, R., Hengstler, J.G. Model-guided identification of a therapeutic strategy to reduce hyperammonemia in liver diseases. Journal of Hepatology 64(4), 860–871 (2016).

22. Ban, E., Franklin, M., Nam, S., Smith, L.R., Wang, H., Wells, R.G., Chaudhuri, O., Liphardt, J.T., Shenoy, V.B. Mechanisms of plastic deformation in collagen networks induced by cellular forces. Biophys. J. 114(2), 450–461 (2018).

23. Ronceray, P., Broedersz, C.P., Lenz, M. Fiber networks amplify active stress. Proc. Natl Acad Sci USA 113(11), 2827–2832 (2016).

24. Stein, A.M., Vader, D.A., Jawerth, L.M., Weitz, D.A., Sander, L.M. An algorithm for extracting the network geometry of three-dimensional collagen gels. Journal of Microscopy 232, 463–475 (2008).

25. Drasdo, D., Zhao, J. An integrative experimental and computational twin modeling approach to understand the clonal dynamics in normal liver. Journal of Hepatology 79(2), 273–276 (2023).

26. Giordano, C., Francone, M., Cundari, G., Pisano, A., d’Amati, G. Myocardial fibrosis: morphologic patterns and role of imaging in diagnosis and prognostication. Cardiovascular Pathology 56: 107391 (2022).

27. Lee, J.M., Kim, H.S., Lee, M., Park, H.S., Kang, S., Nahm, J.H., Park, J.S. Association between pancreatic fibrosis and development of pancreoprivic diabetes after pancreaticoduodenectomy. Scientific Reports 11: 23538 (2021).

28. Hammad, S., Cavalcanti, E., Werle, J., Caruso, M.L., Dropmann, A., Lgnazzi, A., Ebert, M.P., Dooley, S., Giannelli, G. Galunisertib modifies the liver fibrotic composition in the Abcb4KO mouse model. Archives of Toxicology 92(7), 2297--2309 (2018).

29. Hoehme, S., Brulport, M., Bauer, A., Bedawy, E., Schormann, W., Hermes, M., Puppe, V., Gebhardt, R., Zellmer, S., Schwarz, M., Bockamp, E., Timmel, T., Hengstler, J.G., Drasdo, D. Prediction and validation of cell alignment along microvessels as order principle to restore tissue architecture in liver regeneration. Proc. Natl Acad Sci USA 107(23), 10371–10376 (2010).

30. Van Liedekerke, P., Neitsch, J., Johann, T., Alessandri, K., Nassoy, P., Drasdo, D. Quantitative cell-based model predicts mechanical stress response of growing tumor spheroids over various growth conditions and cell lines. PLoS Computational Biology 15(3), e1006273 (2019).

31. Yang, L., Fitie, C.F.C., Van der Werf, K.O., Bennink, M.L., Dijkstra, P.J., Feijen, J. Mechanical properties of single electrospun collagen type I fibers. Biomaterials 29, 955–962 (2008).

32. Ferruzzi, J., Sun, M., Gkousioudi, A., Pillar, A., Roblyer, D., Zhang, Y., Zaman, M.H. Compressive remodeling alters fluid transport properties of collagen networks - implications for tumor growth. Scientific Reports 9, 17151 (2019).

33. Wen, Y. The role of immune cells in liver regeneration. Livers 3(3), 383–396 (2023).

34. Madsen, D.H., Leonard, D., Masedunskas, A., Moyer, A., Jürgensen, H.J., Peters, D.E., Amornphimoltham, P., Selvaraj, A., Yamada, S.S., Brenner, D.A., Burgdorf, S., Engelholm, L.H., Behrendt, N., Holmbeck, K., Weigert, R., Bugge, T.H. M2-like macrophages are responsible for collagen degradation through a mannose receptor-mediated pathway. Journal of Cell Biology 202(6), 951–966 (2013).

35. Winkler, M., Staniczek, T., Kuerschner, S.W., Schmid, C.D., Schoenhaber, H., Cordero, J., Kessler, L., Mathes, A., Sticht, C., Nessling, M., Uvarovskii, A., Anders, A., Zhang, X., Figura, G., Hartmann, D., Mogler, C., Dobreva, F., Schledzweski, K., Geraud, C., Koch, P., Goerdt, S. Endothelial GATA4 controls liver fibrosis and regeneration by preventing a pathogenic switch in angiocrine signaling. Journal of Hepatoycte 74, 380–393 (2021).

36. Rocha, A.S., Vidal, V., Mertz, M., Kendall, T.J., Charlet, A., Okamoto, H., Schedl, A. The angiocrine factor Rspondin3 is a key determinant of liver zonation. Cell Reports 13(9), P1757-1764 (2015).

37. Chen, Y., Li, Q., Tu, K., Wang, Y., Wang, X., Liu, D., Chen, C., Liu, D., Yang, R., Qiu, W., Kang, N. Focal adhesion kinase promotes hepatic stellate cell activation by regulating plasma membrane localization of TGFβ receptor 2. Hepatol. Commun. 4(2), 268--283 (2020).

38. Montagud, A., Beal, J., Tobalina, L., Traynard, P., Subramanian, V., Szalai, B., Alfoldi, R., Puskas, L., Valencia, A., Barillot, E., Saez-Rodriguez, J., Calzone, L. Patient-specific boolean models of signalling networks guide personalised treatments. eLife 11, e72626 (2022).

39. Henderson, N.C., Arnold, T.D., Katamura, Y., Giacomini, M.M., Rodriguez, J., McCarty, J.H., Pellicoro, A., Raschperger, E., Betsholtz, C., Ruminski, P.G., Griggs, D.W., Prinsen, M.J., Maher, J.J., Iredale, J.P., Lacy-Hulbert, A., Adams, R.H., Sheppard, D. Targeting of αv integrin identifies a core molecular pathway that regulates fibrosis in several organs. Nature Medicine 19, 1617--1624 (2013).

40. Ceci, L., Gaudio, E., Kennedy, L. Cellular interactions and crosslink facilitating biliary fibrosis in cholestasis. Cellular and Molecular Gastroenterology and Hepatology 17(4), 553--565 (2024).

41. Hoehme, S., Hammad, S., Boettger, J., Begher-Tibbe B., Bucur, P., Vibert, E., Gebhardt, R., Hengstler, J.G., Drasdo, D. Digital twin demonstrates significance of biomechanical growth control in liver regeneration after partial hepatectomy. iScience 26(1), 105714 (2023).

42. Lara, J., Lopez-Labrador, F.X., Gonzalez-Candelas, F., Berenguer, M., Khudyakov, Y.E. Computational models of liver fibrosis progression for hepatitis C virus chronic infection. BMC Bioinformatics 15(8), S5 (2014).

43. Dutta-Moscato, J., Solovyev, A., Mi, Q., Nishikawa, T., Soto-Gutierrez, A., Fox, I.J., Vodovotz, Y. A multiscale agent-based in silico model of liver fibrosis progression. Frontiers in Bioengineering and Biotechnology 2, 18 (2014).

44. Yoshizawa, M., Sugimoto, M., Tanaka, M., Sakai, Y., Nishikawa, M. Computational simulation of liver fibrosis dynamics. Scientific Reports 12, 14112 (2022).

45. Friedman, A., Hao, W. Mathematical model of liver fibrosis. Mathematical Biosciences & Engineering 14(1), 143–164 (2017).

46. Adhyapok, P., Fu, X., Sluka, J.P., Clendenon, S.G., Sluka, V.D., Wang, Z., Dunn, K., Klaunig, J.E., Glazier, J.A. A computational model of liver tissue damage and repair. PLoS ONE 15(12), e0243451 (2020).

47. Dichamp, J., Celliere, G., Ghallab, A., Hassan, R., Boissier, N., Hofmann, U., Reinders, J., Sezgin, S., Zuehlke, S., Hengstler, J.G., Drasdo, D. In vitro to in vivo acetaminophen hepatotoxicity extrapolation using classical schemes, pharmacodynamic models and a multiscale spatial-temporal liver twin. Front. Bioeng. Biotechnol. 11, 1049564 (2023).

48. Zhao, J., Ghallab, A., Hassan, R., Dooley, S., Hengstler, J.G., Drasdo, D. A digital twin of liver predicts regeneration after drug-induced damage at the level of cell type orchestration. iScience 27, 108077 (2024).

49. Seo, Y.S., Kwon, J.H., Yaqoob, U., Yang, L., De Assuncao, T.M., Simonetto, D.A., Verma, V.K., Shah, V.H. HMGB1 recruits hepatic stellate cells and liver endothelial cells to sites of ethanol-induced parenchymal cell injury. Am. J. Physiol. Gastrointest. Liver Physiol. 305, G838–G848 (2013).

50. Roohani, S., Tacke, F. Liver injury and the macrophage issue: molecular and mechanistic facts and their clinical relevance. Int. J. Mol. Sci. 22(14), 7249 (2021).

51. Van Liedekerke, P., Palm, M.M., Jagiella, N., Drasdo, D. Simulating tissue mechanics with agent-based models: concepts, perspectives and some novel results. Computational Particle Mechanics 2, 401–444 (2015).

52. Ghafoory, S., Breitkopf-Heinlein, K., Li, Q., Scholl, C., Dooley, S., Woelfl, S. Zonation of nitrogen and glucose metabolism gene expression upon acute liver damage in mouse. PLoS One 8(10), e78262 (2013).

53. Ghallab, A., Myllys, M., Holland, C.H., Zaza, A., Murad, W., Hassan, R., Ahmed, Y.A., Abbas, T., Abdelrahim, E.A., Schneider, K.M., Matz-Soja, M., Reinders, J., Gebhardt, R., Berres, M., Hatting, M., Drasdo, D., Saez-Rodriguez, J., Trautwein, C., Hengstler, J.G. Influence of liver fibrosis on lobular zonation. Cells 8(12), 1556 (2019).

54. Teutsch, H.F. The modular microarchitecture of human liver. Hepatology 42(2), 317–325 (2005).

55. Urushima, H., Yuasa, H., Matsubara, T., Kuroda, N., Hara, Y., Inoue, K., Wake, K., Sato, T., Friedman, S.L., Ikeda, K. Activation of hepatic stellate cells requires dissociation of E-cadherin-containing adherens junctions with hepatocytes. The American Journal of Pathology 191, 438--453 (2021).

56. Hammad, S., Hoehme, S., Friebel, A., Von Recklinghausen, I., Othman, A., Begher-Tibbe, B., Reif, R., Godoy, P., Johann, T., Vartak, A., Golka, K., Bucur, O.P., Vibert, E., Marchan, R., Christ, B., Dooley, S., Meyer, C., Ilkavets, I., Dahmen, U., Dirsch, O., Bottger, J., Gebhardt, R., Drasdo, D., Hengstler, J.G. Protocols for staining of bile canalicular and sinusoidal networks of human, mouse and pig livers, three-dimensional reconstruction and quantification of tissue microarchitecture by image processing and analysis. Archives of Toxicology 88, 1161–1183 (2014).

57. Hammad, S., Ogris, C., Othman, A., Erdoesi, P., Schmidt-Heck, W., Biermayer, I., Helm, B., Gao, Y., Pioronska, W., Holland, C.H., D’Alessandro, L.A., de la Torre, C., Sticht, C., Al Aoua, S., Theis, F.J., Bantel, H., Ebert, M.P., Klingmueller, U., Hengster, J.G., Dooley, S., Mueller, N.S. Tolerance of repeated toxic injuries of murine livers is associated with steatosis and inflammation. Cell Death & Disease 14, 414 (2023).

58. Duffield, J.S., Forbes, S.J., Constandinou, C.M., Clay, S., Partolina, M., Vuthoori, S., Wu, S., lang, R., Iredale, J.P. Selective depletion of macrophages reveals distinct, opposing roles during liver injury and repair. J. Clin. Invest. 115(1), 56–65 (2005).

59. Pradere, J., Kluwe, J., De Minicis, S., J, J.J., Gwak, G.Y., Dapito, D.H., Jang, M.K., Guenther, N.D., Mederacke, I., Friedman, R., Dragomir, A., Alomen, C., Schwabe, R.F. Hepatic macrophages but not dendritic cells contribute to liver fibrosis by promoting the survival of activated hepatic stellate cells in mice. Hepatology 58(4), 1461–1473 (2013).

60. Zellmer, S., Schmidt-Heck, W., Godoy, P., Weng, H., Meyer, C., Lehmann, T., Sparna, T., Schormann, W., Hammad, S., Kreutz, C., Timer, J., Weizsacker, F., Thurmann, P.A., Merfort, I., Guthke, R., Dooley, S., Hengstler, J.G., Gebhardt, R. Transcription factors ETF, E2F, and SP-1 are involved in cytokine-independent proliferation of murine hepatocytes. Hepatology 52(6), 2127–2136 (2010).

61. Hammad, S., Braeuning, A., Meyer, C., Mohamed, F., Hengstler, J.G., Dooley, S. A frequent misinterpretation in current research on liver fibrosis: the vessel in the center of CCl_4_-induced pseudolobules is a portal vein. Arch. Toxicol. 91(11):3689–3692 (2017).

62. Lopez-De Leon, A., Rojkind, M. A simple micromethod for collagen and total protein determination in formalin-fixed paraffin-embedded sections. J. Histochem Cytochem 33(8), 737–743 (1985).

