## Supplementary material for "The interplay between biomechanics and cell kinetics explains the spatial pattern in liver fibrosis": SI

[1]*School of Life Science and Engineering, Southwest Jiaotong University, Chengdu, China.* [2]*Leibniz Research Centre for Working Environment and Human Factors, Technical University of Dortmund (IfADo), Dortmund, Germany.* [3]*National Institute for Research in Computer Science and Automation (INRIA de Saclay), Palaiseau, France.* [4]*Molecular Hepatology Section, Department of Medicine II, Medical Faculty Mannheim Heidelberg University, Mannheim, Germany.* [5]*Department of Forensic Medicine and Veterinary Toxicology, Faculty of Veterinary Medicine, South Valley University, 83523-Qena, Egypt.* [6]*BioNamix group, Department Data Analysis and Mathematical Modelling, Ghent University, Gent, Belgium.* [7]*Department of Mathematics and Statistics, University of Massachusetts Amherst, Amherst, USA.* [8]*Department of Dermatology, Venereology and Allergology, University Medical Center and Medical Faculty Mannheim, Heidelberg University, Mannheim, Germany.* [9]*European Center for Angioscience, Medical Faculty Mannheim, Heidelberg University, Mannheim, Germany.* [10]*Hamamatsu Tissue Imaging and Analysis Center (TIGA), BIOQUANT, University Heidelberg, Heidelberg, Germany.* [11]*Institute of Pathology, Hannover Medical School, Hannover,*

*Germany. [12]Clinic for Gastroenterology, Hepatology and Infectious Diseases, Hospital of the Heinrich-Heine-University, Düsseldorf, Germany. [13]Department of Medicine II, Medical Faculty Mannheim Heidelberg University, Mannheim, Germany. [14]Mannheim Institute for Innate Immunoscience (MI3), Medical Faculty Mannheim, Heidelberg University, Mannheim, Germany. [15]DKFZ-Hector Cancer Institute at the University Medical Center, Mannheim, Germany.*

†These two authors contribute equally to this work

**Content**


1. Assessment of liver transaminases, page 5
2. RNA isolation and quantitative real-time (RT)-PCR, page 5
3. Imaging analysis of spatial distribution of proliferating hepatocyte and aHSC, page 6
4. Construction of the Digital twin (DT), page 8
    a. Principle, page 8
    b. Choice of model components, page 9
    c. Choice of parameters, page 9
5. Mathematical modeling of all DT components, page 10
    a. Equation of motion for CBM-based hepatocytes, page 10
    b. Equation of motion for DCM-based hepatocytes, page 12
    c. Equation of motion for macrophages, page 12
    d. Equation of motion for HSC, page 13
    e. Equation of motion for sinusoid spheres, page 14
    f. Modeling of collagen fibers, page 14
        i. Network generation approach, page 14
        ii. The elastic forces and bending forces on a collagen fiber, page 15
        iii. Equation of motion for collagen fiber nodes, page 16
        iv. Measurements of the height of the collagen network, page 17
    g. Forces between cells and elements, page 17
    h. Validation of the collagen model, page 19
        i. Plasticity test of the collagen network, page 20
    i. Organizational structure of collagen network, page 21

   j. Production and degradation of collagen fibers by HSC and Mph, page 23
   k. Signal-gradient based migration of HSC and Mph, page 24
   l. Production of collagen fibers, page 24
   m. Degradation of collagen fibers, page 25
   n. Digestion of dead hepatocytes by Mphs, page 26
6. Choosing parameters of hepatocyte death and proliferation in the simulation of biliary fibrosis, page 26
7. Constructing the heatmap of spatial-temporal distribution of bile ducts, page 27
8. Perturbation simulation of components of the DT, page 28
9. Boolean network model, page 30
   a. Algorithm 0, page 32
10. Computational scheme executed, page 33
   a. Flowcharts, page 34
   b. Algorithm 1-6, page 40
11. Supplementary Figures, page 46
12. Supplementary Tables, page 65
13. Supplementary Movies, page 70
14. References, page 72

### Supplementary Information

**Assessment of liver transaminases.** Blood is collected in Li-Heparin vials from the retrobulbar plexus and centrifuged at 13,000 rpm at 4°C for 6 min. Plasma is subsequently stored at −80°C until further analysis. Then, alanine aminotransferase (ALT), and aspartate aminotransferase (AST) are measured using a Hitachi automatic analyzer (Core facility-Medical Faculty Mannheim, Germany).

**RNA isolation and quantitative real-time (RT)-PCR.** RNA from liver tissues is isolated using the InviTrap spin kit according to the manufacturer's instructions. Liver tissues from the mice are cut into approximately 20 mg and are homogenized with a Precellys Evolution machine two times (20 s, 5 s pulse, 2s break), then the supernatants are collected. Then, 300 µL β-mercaptoethanol-containing lysis solution TR are immediately added to each sample. The resulting lysate is directly pipetted onto the DNA-Binding Spin Filter which is placed in a 2 mL Receiver Tube. Samples are incubated for 1 min at room temperature and are centrifuged at 11000 rpm for 2 min. After discarding the DNA-Binding Spin Filter, 250 µL of 70% ethanol are added to each tube and are completely mixed with the lysates. The mixtures are transferred to RNA-RTA Spin Filters, and are incubated for 1 min. Samples are centrifuged at 11000 rpm for 1 min. Then 600 µL of Wash Buffer R1 are added into the RNA-RTA Spin Filter after discarding the flow-through, then this mixture is centrifuged for 1 min at 11000 rpm. After discarding the flow-through, 700 µL of Wash Buffer R2 are added into the Filter, the same centrifuge step is performed as a wash step using buffer R1. The R2 washing step is repeated once. To eliminate any trace of ethanol in the RNA-RTA Spin Filter membrane, the tubes are centrifuged for 4 min at maximum speed. The RNA-RTA Spin Filters are transferred into RNase-free Elution Tubes and 30 µL of Elution Buffer R is pipetted. After 2 min incubation and 1 min centrifugation at 11000rpm, total RNA is collected and transferred immediately on ice. The concentration is photometrically determined by measuring the RNA solution in a nanocrystalline plate at 260 nm with a Tecan infinite M200. From 1 µg total RNA, cDNA is transcribed with RevertAid H Minus Reverse Transcriptase (Thermo Fischer Scientific, USA, cat. no. EP0451) and used for RT-PCR with PowerUP SYBR Green Master Mix (Life Technologies, USA, cat. no. A25918) in a StepOnePlus RT-PCR System (Applied Biosystems USA from Thermo Fisher Scientific, Germany, cat. no. 4376600). A list of the primer pairs used (Eurofins, Germany) is provided in **Supplementary Table 1**. *Ppia* is used as an internal reference gene. Target gene relative expression is determined with the ΔΔCt method, and a melt curve is created to ensure primer specificity. Each sample is measured in triplicate.

**Imaging analysis of spatial distribution of proliferating hepatocytes and aHSC**

The detection of the dividing cells nuclei (**Fig. 2F**) is obtained by a global thresholding on the Hue and Saturation channels[1] of the images. A post-processing step is applied to remove artifacts (too small and too large, connected components). Furthermore, touching nuclei are separated using the morphological Watershed algorithm. The segmentation function is the Euclidean distance transform on the binary image of the nuclei and the markers are obtained by computing the extended minima transform on the Euclidean transform image. These different morphological operations are standard in image analysis and further details can be found in ref[2]. A final post-processing based on a shape criterion, i.e. the eccentricity, is performed: objects with an eccentricity less than 0.9 are removed.

The stained histological images of HSC are first binarized using local Otsu thresholding to obtain the whole fibrotic pattern. In order to individualize the fibrotic streets, the fibrotic region around each central vein is computed using the so-called *rolling ball algorithm*[3]: a disk of iteratively increasing radius *r* is rolling over the central vein boundary until the area ratio of HSC reaches a certain threshold ($\leq 0.8$ in practice). The maximal radius $r_{max,i}$ returned by the algorithm defines the width of the fibrotic region around the central vein labeled $i$, a region which is called the *central fibrotic region* (**Supplementary Fig. 5A**, here the green circles do not cover all brown staining around CV, which is not needed as our purpose is not to quantify the precise width of the brownish region around the CV, but to identify the fibrotic region between the CVs and quantify it. For this an accurate segmentation of the brown staining around the CV is not required). An individual fibrotic street is therefore defined as the fibrotic pattern linking two neighboring central fibrotic regions. The second main step is the computation of the envelope image of each fibrotic street using (1) a morphological closing operation on the binary image of the street to smooth its contour, and (2) an operation to fill the holes within the street. The envelope image is the resulting binary image after these two operations (see **Supplementary Fig. 5**). The third and final step before quantifying the width distribution of the fibrotic streets is the rotation of each envelope image to align their main axis with the y-axis (see **Supplementary Fig. 5**). Consequently, the width distribution of a given street is defined as the length distribution of the intersections between a horizontal line and the envelope image, the horizontal line being translated in the y-direction. To remove noise in the computation of these lengths, a moving average technique is used. The final results are presented in **Supplementary Fig. 5** as the concatenation of all width distributions of the fibrotic streets for the three mice M1, M2 and M3. In order to check whether or not there is a significant difference between widths computed for stripes connecting close central veins and those connecting central veins further away from each other, the width distributions are presented with respect to the normalized distance to its closest central vein, hence encompassing the spatial nature of the fibrotic street formation (the maximal distance being in the middle of the given fibrotic street). The results do not show any large dependency of the fibrotic street width with respect to the central vein distance.

The computation of the spatial distribution of the dividing cells is performed using a non-parametric kernel density estimator, for which the probability that a nucleus of a dividing cell belongs to the position

$x \in R^2$ is given by:

$$f(x) = \sum_{i=1}^{n} \frac{1}{nh^2} k(\frac{\|x - x_i\|}{h}),$$

where $k$ is a symmetric kernel function (typically a Gaussian kernel is used here), $n$ is the total number of nuclei segmented and $x_i$ is the centroid location of nuclei $i$.

The polygonal contours of the necrotic regions are drawn on the image by a biologist expert using ImageJ. The mask image of the necrotic region is obtained and a morphological closing, using a small size structuring element, is performed to get smoother contours for the necrotic regions.

**Construction of the Digital twin (DT).**

***Principle:*** As the objective is to arrive at a DT of the liver that is able to consistently mimic an increasing number of liver functions inside the same DT (**Supplementary Fig. 1**), the DTs of each individual process or function must remain compatible with each other. This can be achieved by gradually building up the complexity of the liver DT by adding continuously new functionalities to the existing DT in such a way, that those additions do not harm the existing functionalities. I.e., an already existing model element is changed only if it turns out to be incompatible with experimental observations of a novel application. In case an element is modified that has been used in an earlier application, we get back to that earlier application[4,5] and study if this modification of the model affects previous conclusions. For example, the volume compression term in the cell-cell interaction force had to be modified for a DT of regeneration after partial hepatectomy to take into account the strong cell compression forces (Hoehme et. al.[6]). Subsequently it was verified that the conclusions of the DT for regeneration after drug-induced liver damage from an earlier work (Hoehme et. al.[7]) was not compromised (which itself was built upon earlier models as significant extension).

Along this line of argument, the current DT for fibrosis wall formation was constructed as an extension and refinement of our previous cell and liver models.

The physical property and the contact mechanics of the cell (modeled as an elastic object) are based on the studies in (Hoehme et. al.[7]) for the center-based model (CBM) and on studies by (Van Liedekerke et al.[8]) for the DCM. The liver architecture is based on the studies in (Hoehme et al.[6,7]). The intercellular network of signaling of liver injury and regeneration is based on the study (Zhao et al.[9]). All physical properties of the cells and elements (sinusoids and collagens) were taken from literature, the mechanical properties of hepatocytes and collagens were validated with experimental test data. All of the geometric and topological properties of the tissue architectures were based on the experimental imaging data. All of the signaling pathways were based on literature or discussion and confirmation with experts in the field of hepatology.

***Choice of model components:*** The choice of model components is guided by parsimony along the following principles: (i) Identification of those elements that are necessary to explain the question under study; (ii) inclusion of those elements that are either of particular experimental or clinical interest, for example, because these are known or hypothesized to have an important function and are therefore measured, or because these can be experimentally perturbed to test their possible role.

(iii) representing the components by model elements that are, whenever possible, measurable. E.g., interaction forces, cell material properties etc. can in principle be measured. Among those model elements possible we tend to choose first the simplemost. E.g., we approximate cells by the CBM whenever cell deformations can be assumed to be minimal, and by the DCM whenever they may be non-negligible. This is why we chose the DCM only for hepatocytes in the liver damage region, where hepatocytes interact with collagen fibers or bundles (explained below).

***Choice of parameters***: The model parameter choice is guided by the principles explained in refs. (Drasdo et al.[10], Hoehme et al.[7]). In brief: Wherever possible, the model is parameterized by measurable quantities as this permits to either measure those parameters directly or to identify their physiological ranges, and the importance of the parameter is studied by varying it and observing its effect on certain observables, which ideally can be experimentally measured (simulated parameter sensitivity analysis). As the model is stepwise built up, earlier parameter sensitivity analyses still apply to later DT-model applications.

We described further the details of our DT and the new components and physical properties we added in the following.

**Mathematical modeling of all DT components.** To capture the approximate shape of cells and elements (e.g. sinusoids and collagen fibers), they are represented as individual objects with different geometric shapes. The cell surface of a hepatocyte close to the lesion is represented by a triangulated structure consisting of nodes connected by viscoelastic elements (DCM model, **Fig. 2G**) while a hepatocyte far away from the lesion is represented by a homogeneous isotropic elastic adhesive sphere (CBM model)[11]; a macrophages is represented by a homogeneous isotropic elastic adhesive spheres; an HSC is represented as a sphere forming the cell's core body, connected to four semi-flexible chains of spheres[9]; sinusoids are represented as semi-flexible chains of spheres[7]; collagen fibers are represented as elastic springs that can resist stretching and bending.

High resolution cell models as the DCM permit a higher accuracy of biomechanics at the expense of longer simulation times[12-15], while low resolution models as the CBM are less accurate, but computationally more efficient[11].

**Equation of motion for CBM-based hepatocytes**. Each CBM (center-based model) for a hepatocyte is represented as a homogeneous isotropic, elastic, adhesive sphere. It can migrate, grow, divide, and interact with other cells or elements. The position of a hepatocyte $i$ is updated from a force balance equation:

$$\Gamma_{env,i}\vec{v}_i + \sum_j \Gamma_{i,j}(\vec{v}_i - \vec{v}_j) = \sum_j \vec{F}_{ij} + \vec{F}_{mig,i}, \qquad (1)$$

where $\Gamma_{env,i}$ is the $3 \times 3$ - friction matrix with the environment, which we here approximate by a scalar, $\Gamma_{i,j} = \gamma_{\perp}(\vec{e}_{ij} \otimes \vec{e}_{ij}) + \gamma_{\parallel}(I - \vec{e}_{ij} \otimes \vec{e}_{ij})$ is the friction tensor between cell $i$ and sphere (another cell, a sinusoidal element, or a cellular element) $j$, $\vec{e}_{ij}$ is the unit vector from $i$ towards $j$, $\vec{F}_{ij}$ is the corresponding central repulsion/adhesion interaction force, and $\vec{F}_{mig,i}$ is force mimicking the (active) migration force of cell $i$. The central interaction force between cells is computed by[11]:

$$\vec{F}_{ij} = (\frac{4\hat{E}}{3\hat{R}}[a(\delta_{ij})]^3 - \sqrt{8\pi\sigma\hat{E}[a(\delta_{ij})]^3})\vec{e}_{ij}, \qquad (2)$$

where $a(\delta_{ij})$ is the radius of the hepatocyte-hepatocyte contact area, which is related to the indentation of the neighboring deformed cells via $\delta_{ij} = \frac{a^2}{\hat{R}} - \sqrt{\frac{2\pi\sigma}{\hat{E}}}$. $\hat{E}$ and $\hat{R}$ are defined as $\hat{E} = (\frac{1-\nu_i^2}{E_i} + \frac{1-\nu_j^2}{E_j})^{-1}$ and $\hat{R} = (\frac{1}{r_i} + \frac{1}{r_j})^{-1}$, with $E_i$ and $E_j$ being the Young's moduli, $\nu_i$ and $\nu_j$ the Poisson ratios, and $r_i$ and $r_j$ the radii of $i$ and $j$. To consider the limited cell volume compressibility in a pairwise cell-cell interaction force, we further fitted $\hat{E}$ by a function that depends on the local average distance $\hat{d}_{ij} = 1 - d_{ij}/(r_i + r_j)$ for a bulk cell in the simulated experiment: $\hat{E} = \{\hat{E}, 0 \leq \hat{d}_{ij} \leq 0.08 \; a_0 + a_1\hat{d}_{ij} + \cdots + a_6\hat{d}_{ij}^6, 0.08 < \hat{d}_{ij}$, where $d_{ij}$ is the distance between $i$ and $j$ (see more details of choosing the values of $a_i, i = 0, \ldots, 6$ in Van Liedekerke et al.[8]. The (active) migration force is approximated by $\vec{F}_{mig,i} = f_{dir} \cdot \vec{e}_i + \sqrt{2{\Gamma_{env,i}}^2 D_i} \cdot \vec{\eta}_i$, where $f_{dir}$ has been chosen as a constant force magnitude, $\vec{e}_i$ is the unit vector of migration direction, $D_i$ is the diffusion constant of $i$, $\vec{\eta}_i$ is an uncorrelated noise term with amplitude $\langle \eta_{in}(t)\eta_{jm}(t') \rangle = \delta_{ij}\delta_{mn}\delta(t - t')$, $t$ and $t'$ denote time, and $m, n \in (x, y, z)$ denote the coordinates[7].

**Equation of motion for DCM-based hepatocytes**. Each DCM (deformable cell model)-based hepatocyte is represented by nodes that are connected by parametrized viscoelastic elements. The nodes constitute a flexible triangulated surface which models the boundary (membrane and actin cortex) of the cell. To keep the model computationally tractable, the inside of the cell is assumed to be homogeneous and slightly compressible. The adhesion between two cells is determined by the attraction forces between those nodes of the cells that are in each other's neighborhood. To update the configuration for each cell, a force-balance equation for every of its nodes has to be solved. For a hepatocyte, the position of each of its nodes $i$ can be updated by solving the following equation of motion:

$$\Gamma_{env,i}\vec{v}_i + \sum_j \Gamma_{i,j}(\vec{v}_i - \vec{v}_j) = \sum_j \vec{F}_{e,ij} + \sum_m \vec{F}_{m,i} + \vec{F}_{vol,i} + \sum_T \vec{F}_{T,i} + \vec{F}_{rep,i} + \vec{F}_{adh,i}, \quad (3)$$

where $\Gamma_{env,i}$ is the friction coefficient with the environment, $\Gamma_{i,j} = \gamma_{\perp}(\vec{e}_{ij}\otimes\vec{e}_{ij}) + \gamma_{\parallel}(I - \vec{e}_{ij}\otimes\vec{e}_{ij})$ is the friction tensor between $i$ and another node $j$, and $\vec{e}_{ij}$ is the unit vector from $i$ towards $j$ (whereby $j$ can be either a node of the hepatocyte or a neighboring structure the hepatocyte interacts with), $\vec{F}_{e,ij}$ is the in-plane elastic force with node $j$, $\vec{F}_{m,i}$ is the bending force with node $m$, $\vec{F}_{vol,i}$ is the volume force controlled by the cell compressibility, $\sum_T \vec{F}_{T,i}$ is the force that avoids excessive triangle distortion of triangle $T$ which contains $i$, $\vec{F}_{rep,i}$ is the repulsion force on the local surface, $\vec{F}_{adh,i}$ is the adhesion force with nearby objects (for more details on the DCM and equation (3), see Van Liedekerke et al.[8]).

**Equation of motion for macrophages**. Each macrophage is represented as a homogeneous isotropic, elastic, adhesive sphere. The position of macrophage $i$ is updated by solving the following equation:

$\Gamma_{env,i}\vec{v}_i + \sum_j \Gamma_{i,j}(\vec{v}_i - \vec{v}_j) = \sum_j \vec{F}_{ij} + \vec{F}_{mig,i}$, (4)

where $\Gamma_{env,i}$ is the friction coefficient with the environment, $\Gamma_{i,j}$ is the friction tensor between $i$ and $j$ (same as equation (1)), $\vec{F}_{ij}$ is the corresponding central repulsion/adhesion interaction force, $\vec{F}_{mig,i} = f_{dir} \cdot (\vec{e}_i + \vec{\theta}_i)/(\|\vec{e}_i + \vec{\theta}_i\|)$ is the migration force, where $f_{dir}$ has been chosen as a constant force magnitude, $\vec{e}_i$ is the unit vector of migration direction, $\vec{\theta}_i$ is a random direction unit vector of $i$, where $< \vec{\theta}_i >= \vec{e}_i$.

**Equation of motion for HSC**. The core body of an HSC is modeled as a homogeneous isotropic, elastic, adhesive sphere (mainly representing the HSC's nucleus) with four semi-flexible chains of spheres as "arms" (to mimic the long HSC's protruding branches). The position of an HSC sphere $i$ is updated by solving the following equation of motion[7]:

$\Gamma_{env,i}\vec{v}_i + \sum_j \Gamma_{i,j}(\vec{v}_i - \vec{v}_j) = \sum_j \vec{F}_{ij} + \vec{F}_{mig,i} + \sum_k \vec{F}_{ela,ik}$, (5)

where $\Gamma_{env,i}$ is the friction coefficient with the environment, $\Gamma_{i,j}$ is the friction tensor between $i$ and another cell or element $j$ (same as equation (1)), $\vec{F}_{ij}$ is the interaction force between $i$ and another cell or element $j$ (same as equation (2)). $\vec{F}_{ela,ik}$ represents elastic force between the body center $i$ and the connected spheres $k$ of its arms. $\vec{F}_{mig,i} = f_{dir} \cdot (\vec{e}_i + \vec{\theta}_i)/(\|\vec{e}_i + \vec{\theta}_i\|)$ is the migration force to drive $i$ to migrate, where $f_{dir}$ has been chosen as a constant force magnitude, $\vec{e}_i$ is the unit vector of migration direction, $\vec{\theta}_i$ is a random direction unit vector of $i$, where $< \vec{\theta}_i >= \vec{e}_i$. The equation of motion for a sphere representing element $i$ of an HSC's arm is approximated by:

$\Gamma_{env,i}\vec{v}_i + \sum_j \Gamma_{i,j}(\vec{v}_i - \vec{v}_j) = \sum_j \vec{F}_{ij} + \sum_k \vec{F}_{ela,ik}$, (5b)

where $j$ represents the cells or elements interacting with $i$, and $k$ represents the spheres in the same

HSC's arm of element $i$ and connecting to $i$.

**Equation of motion for the sinusoid spheres**. Sinusoids are modeled as chains of semi-flexible spheres[6]. For each sinusoid sphere $i$, the position of $i$ is updated by solving the following equation of motion[7]:

$$\Gamma_{env,i}\vec{v}_i = \sum_j \left(\Gamma_{\parallel,SE}(\vec{w}_{ij} - \vec{e}_{ij}(\vec{w}_{ij} \cdot \vec{e}_{ij})) + \vec{F}_{ij}\right) + \sum_k \left(\Gamma_{\parallel,SS}\big(\vec{w}_{ik} - \vec{e}_{ik}(\vec{w}_{ik} \cdot \vec{e}_{ik})\big) + \vec{F}_{ik}\right) + \vec{F}_{i,ela}, \qquad (6)$$

where $\Gamma_{env,i}$ is the friction coefficient with the environment, $\Gamma_{\parallel,SE}$ denotes the longitudinal friction between sinusoid sphere $i$ and its interacting cells or elements, for example hepatocyte $j$, $\vec{w}_{ij} = \vec{v}_j - \vec{v}_i$ is the difference of velocity between $i$ and $j$, $\vec{e}_{ij}$ is the unit direction vector from $i$ towards $j$, $\vec{F}_{ij}$ is the interaction force between $i$ and $j$ (same as equation (2)), $\Gamma_{\parallel,SS}$ denotes the longitudinal friction between two sinusoid spheres $i$ and $k$, $\vec{w}_{ik} = \vec{v}_k - \vec{v}_i$ is the difference of velocity between $i$ and $k$, $\vec{e}_{ik}$ is the unit direction vector from $i$ towards $k$, $\vec{F}_{ik}$ is the interaction force between $i$ and $k$, $\vec{F}_{i,ela}$ is the spring force that arises from the chain connections between spheres belonging to the same sinusoid.

**Modeling of collagen fibers**

In the fibrosis development simulations, the collagen fibers are generated by the aHSCs. In order to calibrate the network parameters, we perform a simulated compression test and compare the simulation results with experimental data. The network is generated as follows.

**Network generation approach**. In a given gel space $\Omega_g$ (volume of $\Omega_g$ is $V_g$), the total volume of the collagen fiber $V_c$ is determined by $V_c = V_g \rho_c \upsilon_c$, where $\upsilon_c = 1.89\, ml/g$ is the specific volume of collagen[16], $\rho_c = 1.0\, mg/ml$ is the mass density of collagen[17]. The total length of collagen fiber, $L_{tot}$ deposited in the given space is given by $L_{tot} = V_c/\pi r_c^2$, where $r_c$ is the radius of collagen fiber. A network is grown by one fiber at a time until the network reaches the total fiber length $L_{tot}$. The fiber network is generated in the following steps:

(1) Insert a fiber $i$ in $\Omega_g$. It is assigned an initial point $u_0^i$ and an initial tangent direction $\vec{\theta}_0^i$, both are chosen from a uniform distribution. A fiber length $L_f$ is also assigned to $i$, where $< L_f > \sim 6\mu m$ (see ref[9]). Grow the fiber after $n$ extensions of segments in particular direction $\vec{\theta}_{j+1}^i = \vec{\theta}_j^i + \Delta\vec{\theta}_j^i, j = 0, \ldots, n-1$ until $L_f$ is reached. The length of each extended segment is $L_c = 3\ \mu m$ (see ref[9]). Stop until the target total fiber length $L_{tot}$ is reached.

(2) Apply a random force onto each of the fibers and update their positions. Crosslinks are added if any two fiber segments are closer than a distance threshold of 1.5 μm (**Supplementary Fig. 6A**).

(3) Repeat step (2) until all fibers in $\Omega_g$ are connected into one network through crosslinks.

**The elastic forces and bending forces on a collagen fiber**. Suppose a collagen fiber consists of several nodes and edges (**Supplementary Fig. 6B**). The elastic force at node $x_n$ can be written as $\vec{f_e}(x_n) = k(\Delta l_{n-1}\frac{\vec{l}_{n-1}}{|\vec{l}_{n-1}|} + \Delta l_n \frac{\vec{l}_n}{|\vec{l}_n|})$, where $k$ is the elasticity constant, $\Delta l_n$ is the difference between the current length $|\vec{l}_n|$ and the equilibrium length $|\vec{l}_{n,0}|$ of the spring connecting the vertices (nodes) $n+1$ and $n$ (see ref[18]).

The bending energy at $x_n$ can be written as $U(x_n) = \frac{1}{2}\int_{x_0}^{x_n} EI\kappa(s)^2 ds$, where $E$ is the Young's modulus, $I$ is the second moment of area, $\kappa$ is the local curvature at $x_n$. The bending energy due to the bending angle $\theta_n$ at $x_n$ can be approximated as:

$\Delta U(x_n) = U(x_n) - U(x_{n-1}) \approx \frac{1}{2}EI\kappa(x_n)^2|l_n| \approx \frac{1}{2}EI(\theta_n - \theta_{n,0})^2/|l_n|$ (for $\kappa(x_n) \approx \theta_n/l_n$), where $\theta_{n,0}$ is a constant equilibrium angle for $x_n$. The bending force is evaluated by derivative of involved $\Delta U(x_{n+i})$ over $x_n$:

$$\vec{f}_b(x_n) = \sum_{i=-1}^{1} \nabla_{x_n} \Delta U(x_{n+i}) = \sum_{i=-1}^{1} \partial \Delta U(x_{n+i})/(\partial cos\,(\theta_{n+i} - \theta_{n,0}))\nabla_{x_n} cos(\theta_{n+i} - \theta_{n,0}), \quad (7)$$

We denote $\vec{l}_n = \vec{x}_{n+1} - \vec{x}_n$ and $\vec{l}_{n-1} = \vec{x}_n - \vec{x}_{n-1}$, we can get $cos\,(\theta_n - \theta_{n,0}) = \frac{(R_{\theta_1}\vec{l}_n)^T (R_{\theta_2}\vec{l}_{n-1})}{|l_n||l_{n-1}|}$, where $R_{\theta_1}$ and $R_{\theta_2}$ are two rotational matrices satisfying $\theta_1 - \theta_2 = \theta_{n,0}$. It is easy to get $R_{\theta_2}^T R_{\theta_1} = R_{\theta_1-\theta_2} = R_{\theta_{n,0}}$, which is also a constant matrix. In addition, we also have $\frac{\partial \Delta U(x_n)}{\partial cos\,(\theta_n - \theta_{n,0})} = -EI \frac{\theta_n - \theta_{n,0}}{sin\,(\theta_n - \theta_{n,0})}$. Using the property of vector calculus identities $\nabla(A \cdot B) = B \cdot (\nabla A) + A \cdot (\nabla B)$, we have:

$$\nabla_{x_n} cos(\theta_n - \theta_{n,0}) = \nabla_{x_n} \frac{(R_{\theta_1}\vec{l}_n)^T (R_{\theta_2}\vec{l}_{n-1})}{|l_n||l_{n-1}|} = \frac{\nabla_{x_n}\left((R_{\theta_1}\vec{l}_n)^T (R_{\theta_2}\vec{l}_{n-1})\right)|l_n||l_{n-1}| - (R_{\theta_1}\vec{l}_n)^T (R_{\theta_2}\vec{l}_{n-1})\nabla_{x_n}(|l_n||l_{n-1}|)}{|l_n|^2 |l_{n-1}|^2} =$$

$$\frac{R_{\theta_{n,0}}\vec{l}_n - R_{-\theta_{n,0}}\vec{l}_{n-1}}{|l_n||l_{n-1}|} + \frac{\vec{l}_n^{\,T}(R_{-\theta_{n,0}}\vec{l}_{n-1})}{|l_n||l_{n-1}|}\left(\frac{\vec{l}_n}{|l_n|^2} - \frac{\vec{l}_{n-1}}{|l_{n-1}|^2}\right), \quad (8)$$

Accordingly, we also have

$$\nabla_{x_n} cos(\theta_{n-1} - \theta_{n,0}) = \frac{-1}{|l_{n-1}||l_{n-2}|}\left(\frac{\vec{l}_{n-1}^{\,T} R_{-\theta_{n,0}}\vec{l}_{n-2}}{|l_{n-1}|^2}\vec{l}_{n-1} - R_{-\theta_{n,0}}\vec{l}_{n-2}\right) \text{ and}$$

$$\nabla_{x_n} cos(\theta_{n+1} - \theta_{n,0}) = \frac{1}{|l_{n+1}||l_n|}\left(\frac{\vec{l}_{n+1}^{\,T} R_{-\theta_{n,0}}\vec{l}_n}{|l_n|^2}\vec{l}_n - R_{\theta_{n,0}}\vec{l}_{n+1}\right).$$

We denote $\nabla_{x_n} cos(\theta_{n-1} - \theta_{n,0})$ as a vector $\vec{d}_1(x_n)$, $\nabla_{x_n} cos(\theta_n - \theta_{n,0})$ as a vector $\vec{d}_2(x_n)$, and $\nabla_{x_n} cos(\theta_{n+1} - \theta_{n,0})$ as a vector $\vec{d}_3(x_n)$,. Thus, for each node $x_n$ in the fiber, the bending force is obtained as:

$$\vec{f}_b(x_n) = \frac{-EI}{|l_{n-1}|}\frac{\theta_{n-1} - \theta_{n,0}}{sinsin\,(\theta_{n-1} - \theta_{n,0})}\vec{d}_1(x_n) + \frac{-EI}{|l_n|}\frac{\theta_n - \theta_{n,0}}{sinsin\,(\theta_n - \theta_{n,0})}\vec{d}_2(x_n) + \frac{-EI}{|l_{n+1}|}\frac{\theta_{n+1} - \theta_{n,0}}{sinsin\,(\theta_{n+1} - \theta_{n,0})}\vec{d}_3(x_n), \quad (9)$$

**Equation of motion for collagen fiber nodes**. The center of mass position of a collagen fiber node $i$ is obtained by an overdamped Langevin equation of motion, which summarizes all forces exerted on $i$:

$$\Gamma_{c,i}\vec{v}_i + \sum_j \Gamma_{i,j}(\vec{v}_i - \vec{v}_j) = \sum_j \vec{F}_{c,el,ij} + \sum_j \vec{F}_{c,be,ij} + \sum_j \vec{F}_{cc,ij}, \qquad (10)$$

where $\Gamma_{c,i} = \frac{1}{2}\sum_j \gamma_\perp^{c,j}(\vec{e}_{ij}\otimes\vec{e}_{ij}) + \gamma_\parallel^{c,j}(I - \vec{e}_{ij}\otimes\vec{e}_{ij})$ is the collagen-environment friction tensor, $\vec{e}_{ij}$ is the unit orientation vector of fiber $j$ connecting node $i$. $\gamma_\perp^{c,j}$ and $\gamma_\parallel^{c,j}$ are the perpendicular and parallel components of the friction of fiber $j$. $\gamma_\perp^{c,j}$ and $\gamma_\parallel^{c,j}$ can be approximated as $\gamma_\perp^{c,j} = 4\pi\eta_0 L_j/(ln\left(\frac{L_j}{2r_j}\right) + \rho_\perp)$ and $\gamma_\parallel^{c,j} = 2\pi\eta_0 L_j/(ln\left(\frac{L_j}{2r_j}\right) + \rho_\parallel$ (see ref[19]), where $L_j$ and $r_j$ are the length and radius of fiber $j$. $\rho_\perp$ and $\rho_\parallel$ are two correction factors for approximating friction coefficient for rod-like objects[20]. $\Gamma_{i,j}$ is the friction tensor between the spring containing $i$ and other cell or element $j$ (for example, a sinusoidal element) in contact with the spring (same as in equation (3)). The first term of the right-hand side (rhs) of equation (10) represents the elastic force exerted on $i$ from elastic springs connecting $i$. The second term of rhs of equation (10) represents the bending force exerted on $i$ from bending springs connecting $i$. The third term of rhs of equation (10) represents the interaction force between the spring containing $i$ and other elements (e.g. hepatocytes or sinusoids).

**Measurement of the height of the collagen network**

The height of the collagen network is measured by the gyration radius in $y$-axis (the direction of pressure from other cells) of the collagen network by $r_{gyr} = \sqrt{\frac{1}{N}\sum_i (r_{i,y} - \frac{1}{N}\sum_j r_{j,y})^2}$, where $r_{i,y}$ is the $y$-coordinate of collagen node $i$. This gyration radius is a measure for the spatial spread of the network in y-direction.

**Forces between cells and elements**. The total force between cell/element $i$ and cell/element $j$ can be approximated by the sum of a repulsive and an adhesive force, which are characterized by a function of the geometrical overlap $\delta_{ij}$. Since the cellular adhesion force is due to the specific ligand-receptor on cell surface, we assume that there is only adhesive force between two hepatocytes, which can be approximated by the Johnson-Kendal-Roberts (JKR) model following a previous study[11]. For two hepatocytes $i$ and $j$, the interaction force is computed using equation (2). For the other cells and elements (e.g. collagen fiber), we use Hertz model to approximate the repulsive force between them[21].

For two spherical cell objects $i$ and $j$, the Hertz repulsive force is computed by

$$\vec{F}_{rep,ij} = \frac{4\hat{E}}{3}\sqrt{\hat{R}}\delta_{ij}{}^{3/2}\frac{\vec{e}_{ij}}{|\vec{e}_{ij}|}, \qquad (11)$$

where $\hat{E}$ and $\hat{R}$ are as defined in equation (2). $\vec{e}_{ij}$ is the unit vector from $i$ towards $j$. $\hat{R} = (\frac{1}{r_i} + \frac{1}{r_j})^{-1}$, where $r_i$ and $r_j$ are the radii of cell objects $i$ and $j$.

For two collagen fiber springs $i$ and $j$, the Hertz repulsive force is computed by

$$\vec{F}_{rep,ij} = \frac{4\hat{E}}{3}\sqrt{\hat{R}}\delta_{ij}{}^{3/2}\frac{\vec{e}_{ij}}{|\vec{e}_{ij}|}, \qquad (12)$$

where $r_i$ and $r_j$ are the radii of springs $i$ and $j$, which are associated with the radii of the two interacting collagen filament bundles. In our model, $r_i = r_j$, hence $\hat{R} = r_i$.

For a spherical cell object $i$ and a collagen fiber $j$, the Hertz force repulsive force can be shown to be well approximated by (Buttenschoen et. al., in preparation):

$$\vec{F}_{rep,ij} = \frac{2\hat{E}}{3}(\sqrt{\hat{R}} + \sqrt{r_i})\delta_{ij}{}^{3/2}\frac{\vec{e}_{ij}}{|\vec{e}_{ij}|}, \qquad (13)$$

where $r_j$ is the radius of the fiber spring $j$.

For a triangle of DCM-based hepatocyte $i$ and fiber spring $j$, we apply the algorithm of triangle-cylinder collision detection[22] to check the collision type between $i$ and $j$. If $i$ and $j$ contact with parallel axes, the Hertz force repulsive force is computed by[23]

$$\vec{F}_{rep,ij} = \frac{\pi}{4}\hat{E}L_{ij}\delta_{ij}\frac{\vec{e}_{ij}}{|\vec{e}_{ij}|}, \qquad (14)$$

where $L_{ij}$ is the length of the overlapped section between $i$ and $j$ (see ref[23] for the calculation of $L_{ij}$). If $i$ and $j$ contact with non-parallel axes, the Hertz force repulsive force is computed by[23]

$$\vec{F}_{rep,ij} = 2\hat{E}r_j\delta_{ij}\frac{\vec{e}_{ij}}{|\vec{e}_{ij}|}, \qquad (15)$$

where $r_j$ is the radius of $j$.

**Validation of the collagen model**

We compare simulations of our collagen model directly with experimental data of two types of mechanical tests. This permits us to validate the structure of the model and to fine-tune the model parameters. We first apply the model to simulate the deformation of a single collagen fiber upon external force as depicted in ref[24]. A fiber with length of $3\mu m$ consisting of 20 segments is fixed at its two ends. A force is applied perpendicularly on the fiber on three different positions (we assume y-axis is along the direction of the applied force, **Supplementary Fig. 6C**). The displacement of the fiber is computed as the largest displacement of the fiber node on the y-axis. The simulated relationship between fiber displacement and magnitude of applied force can perfectly fit the experimental observation (**Supplementary Fig. 6D**).

We then apply the model to simulate the stress of the collagen network under compression (**Supplementary Fig. 6E**). A fiber network with size of 42.8×42.8×42.8 $\mu m^3$ is generated. The collagen nodes in the top layer (< 3 $\mu m$ from the top) is fixed. The collagen network is compressed by pushing the nodes in the bottom layer (< 3 $\mu m$ from the bottom) upwards (y-axis is along the up-bottom direction). The strain is calculated as $(\sum_{i\in top} y_i - \sum_{i\in bottom} y_i)/(\sum_{i\in top} y_{i,0} - \sum_{i\in bottom} y_{i,0})$, where $y_i$ and $y_{i,0}$ are the y-coordinate and initial y-coordinate of collagen node $i$. We use two approaches to measure stress.

**Approach one**. Stress is calculated as $\sum_{i \in top} f_{y,i}/A$, where $f_{y,i}$ is the projection of force on the y-axis of collagen node $i$, $A = (x_{max} - x_{min}) \times (z_{max} - z_{min})$, $x_{max}$, $x_{min}$, $z_{max}$ and $z_{min}$ are the maximum and minimum x-coordinates and z-coordinates of all collagen nodes in the network, respectively.

**Approach two**. This approach is based on the concept of virial stress, a well-known concept in particle based simulations, whereby the volume averaged stress tensor reads $\sigma = \frac{1}{V} \sum_{i,j:i>j} \vec{f}_{ij} \otimes \vec{e}_{ij}$, where $\vec{e}_{ij}$ is the string vector between collagen nodes $i$ and $j$, $\vec{f}_{ij}$ is the force exerted on $\vec{e}_{ij}$, $V$ is the volume of the cubic space occupied by the network, that is 42.8×42.8×42.8 $\mu m^3$. Then the stress is taken as the y-element of the diagonal of the virial stress tensor $\sigma$ (compression is along the y-axis). The strain-stress curve of our simulation can fit the experimental data[25] (**Supplementary Fig. 6G**).

**Plasticity test of the collagen network**. During the progression of liver fibrosis, HSCs produce collagen fibers while macrophages degrade collagen fibers simultaneously. The dynamic of production and degradation of collagen fibers might change the mechanical property of the collagen network (e.g. plasticity instead of elasticity). Here we do one plasticity test by rebuilding the collagen network which we use for the compression test (**Supplementary Fig. 6F**):

(1) We compress the collagen network until it experiences a strain of 15%.

(2) We randomly remove some collagen fibers from the network (the removed number of fibers is calculated according to a given fraction, e.g. if the given fraction is 5%, then 5% of the fibers would be removed. Any crosslink node joint with the removed fiber would be removed as well) and add the same number of new collagen fibers back.

(3) The newly added collagen fibers are cross-linked within the network.

(4) We release the compressed network and measure the stress-strain curve and compare it with the

curve of the pure compression test.

Such rebuilding process (removing and adding fibers) is to mimic the process that Mphs degrade while HSCs produce collagen fibers simultaneously during fibrosis progression. As shown in **Supplementary Fig. 6G**, after removing and adding new collagen fibers, the stress of the collagen network drops significantly, and it displays the property of plasticity (permanent deformation after stress reaches to zero) (**Supplementary Fig. 6G**).

**Organizational structure of collagen network**

The spatial organization of the collagen fibers during the formation of a fibrotic wall is still unknown. Therefore, we assume six different scenarios (S) of the organizational structure of collagen network: the collagen fibers are deposited as single fibers and not anchoring to the nearest sinusoids (S1); as single fibers and anchoring to the nearest sinusoids (S2); as single fibers and crosslinked to form a network, but not anchoring to the nearest sinusoids (S3); as single fibers and crosslinked to form a network, and anchoring to the nearest sinusoids (S4); as single fibers firstly and gradually crosslinked to form a network, but not anchoring to the nearest sinusoids (S5); as single fibers firstly and gradually crosslink to from a network, and anchoring to the nearest sinusoids (S6) (**Supplementary Fig. 10A**). One simplified section of the two-lobules system without HSC and Mph in DT (the lesion region of the section is set to locate in the middle of the section, with the width of 4 hepatocytes' diameter) is run for 21 days to test these scenarios. The initial lesion size is set to 45 μm (in y-axis). Six rounds of $CCl_4$ are injected at day 0, 3, 7, 10, 14, 17, respectively. At the moment of the second injection, namely, day 3, all collagen fibers are generated homogeneously distributed in the lesion. The total length of generated collagen fibers are calculated according to the given collagen density (see section of **Network generation approach** for details of transforming the density into the total length of fibers).

Hepatocytes outside of the lesion region proliferate to fill up the lesion and compress the fibers (the proliferation rate data is from ref[7]. For each scenario, we study three different collagen densities: 0.4 mg/mL, 1.0 mg/mL, and 2.0 mg/mL. The deformation of the collagen network is measured by its gyration radius in y-axis (see section of **Measurement of the height of the collagen network**). For each density within each scenario, three simulation runs are performed to calculate the average and standard deviation.

Our results show that in S1 the proliferating hepatocytes result in a displacement of the fibers, or pass through the fibers. As a consequence, part of the fibers remain between the hepatocytes (scenario 1 in **Supplementary Fig. 10B-D**). In S2, fibers are assumed to anchor to the sinusoids. They can partially resist hepatocyte movement as they bend upon contact with the hepatocytes (scenario 2 in **Supplementary Fig. 10B-D**, gyration radius curve in **Supplementary Fig. 10E-G**). The difference between S3 and S4 is similar to the difference between S1 and S2. This can be attributed to the fact that the crosslinked fibers are integrated into one scaffold that is much more difficult to be displaced or compressed than individual fibers as long as cells cannot push or pass through the network. In addition, in S4, fibers are assumed to anchor to the sinusoids, so it is even more difficult for hepatocytes to deform them (reflected by the gyration radius curve in **Supplementary Fig. 10E-G**, scenarios 3 & 4 in **Supplementary Fig. 10B-D**). Moreover, as we increase the collagen density from 0.4 mg/mL to 2.0 mg/mL, the fiber network is more resistant to the penetration of proliferating hepatocytes. In the case of the highest collagen density (2.0 mg/mL), the network is barely deformed compared to those with lower densities (**Supplementary Fig. 10D** and gyration radius curve in **Supplementary Fig. 10G**). In S5, fibers are compressed and concentrated in the middle of the lesion region and form a wall-like structure (scenario 5 in **Supplementary Fig. 10B-D**). This can be attributed to the fact that the fiber network is formed through crosslinks gradually. So compared to the network formed immediately through crosslinks (in S3), it is easier to be deformed. In addition, since all fibers are eventually crosslinked, they are not distributed scatteredly as shown in S1 and S2. Moreover, under each collagen density, the thickness of the fiber network in S5 is among the thinnest (gyration radius curve in **Supplementary Fig. 10E-G**). In S6, the fiber network is more resistant to the penetration of proliferating hepatocytes than that in S5 (scenario 6 in **Supplementary Fig. 10B-D**). This is because the fibers are anchored to the nearest sinusoids in S6.

These simulation results suggest that the crosslinks between collagen fibers are important to maintain the elastic property of the collagen network. Otherwise, fibers without crosslinks cannot remain as integrity against the penetration of proliferating hepatocytes (**Supplementary Fig. 10B-D**). However, if the crosslinks are formed too early or if the fibers can anchor to the sinusoids, the network is highly resistant to the compression from hepatocytes. Among all six scenarios of organizational structure of collagens, scenario (5), namely, collagen initiated as single fibers and gradually crosslinked but not anchoring to the closest sinusoids seems to be the optimal scenario for our simulation of fibrosis progression. In addition, the collagen network under low density (0.4 mg/mL) is too sparse to form a thick wall-like structure as observed in experiments (**Fig. 1A**) while the network under high density (2.0 mg/mL) is too dense for hepatocytes to deform (**Supplementary Fig. 10D**). Therefore, we take the median density 1.0 mg/mL for our next simulation.

**Production and degradation of collagen fibers by HSC and Mph**. During liver fibrosis, excessive amounts of ECM proteins are secreted by activated HSC[26]. Collagen fibers are degraded by macrophages[43]. We model the whole procedure of the migration of HSC and macrophages into the lesion, producing and degrading collagen fibers, respectively. In addition, we assume that if the local collagen density at the location of an aHSC (calculated as the density of collagen fibers within the sphere whose radius is equal to HSC's branch) is above a threshold, it would stop producing collagen fiber.

**Signal-gradient based migration of HSC and Mph**. Previous studies have reported that injured hepatocytes can produce DAMP signals to activate macrophages[27,28]. Here we assume that DAMP is produced by necrotic hepatocytes. HSC and Mph migrate following the gradient of DAMP (**Supplementary Fig. 8A**). The concentration of DAMP $\phi_{DAMP}$ is updated by solving a partial differential equation (PDE)[9]:

$$\frac{\partial \phi_{DAMP}}{\partial t} = \nabla(D_{DAMP} \nabla \phi_{DAMP}) + s_{DAMP}(\sum_j \delta(x - x_j)) - \gamma_{DAMP}\phi_{DAMP}, \qquad (16)$$

where $D_{DAMP}$, $s_{DAMP}$, $\gamma_{DAMP}$ are the diffusion coefficient, production rate, decay rate of DAMP, respectively; $x_j$ denotes the position of necrotic hepatocyte $j$, which produces DAMP. The first term of the rhs of equation (16) denotes the diffusion of DAMPs, the second term denotes the production of DAMPs (produced by HSC which is localized at positions $x_j$, where index $j$ enumerates the HSCs, the center of mass of the cell body of HSC $j$ is set as the origin of the source). The simulation domain $\Omega$ is set as a box large enough to contain the entire lobule. For a macrophage or HSC $i$, if the concentration of DAMP at the location of $i$, $\phi_{DAMP,i}$ becomes higher than a threshold, a migration force $\vec{F}_{mig,i} = f_{mig,i}(\frac{\nabla \phi_{DAMP,i}}{\|\nabla \phi_{DAMP,i}\|} + \vec{\theta}_i)/(\|\frac{\nabla \phi_{DAMP,i}}{\|\nabla \phi_{DAMP,i}\|} + \vec{\theta}_i\|), f_{mig,i} \sim N(F_{mig,i,mean}, F_{mig,i,sd})$ is applied on $i$, where $F_{mig,i,mean}$ and $F_{mig,i,sd}$ are the mean and standard deviation of migration force magnitude calibrated from the mean and standard deviation of migration speed of $i$ [29,30], $\vec{\theta}_i$ is a random direction unit vector of $i$, where $<\vec{\theta}_i> = \frac{\nabla \phi_{DAMP,i}}{\|\nabla \phi_{DAMP,i}\|}$.

**Production of collagen fibers**. Previous studies have reported that HSCs proliferate and become activated. Activated HSCs are the major source of collagens and other matrix proteins[31]. It is also documented that the recovery from liver fibrosis is accompanied by apoptosis of activated HSC[32]. Here we assume that after each dose of $CCl_4$ injection, HSCs in the lesion are killed, activated and induced to proliferate. The fraction of dead and proliferating HSCs after the injection of $CCl_4$ are taken from the data of Lee et al.[33]. The HSC proliferation is taken at 32 hours after the dose injection while the death of HSC is gradually induced after the dose injection until 64 hours after the dose injection[33]. The activated HSC can produce collagen fibers. We observe that after the second dose of $CCl_4$, collagen fibers gradually accumulate within one day after the injection. We denote the coordinates of the head spheres of all activated HSCs as $V_{aHSC} \equiv \{x_i\}$. For a given collagen mass density $\rho_c$, the total length of deposited collagen fiber is $L_{tot}$. The fibers are produced equally by all aHSCs within one day after the injection of $CCl_4$. For a deposited fiber $j$, the initial position $x_{j,0}$, is sampled randomly from $V_{aHSC}$ by HSC $i$, which satisfies $\|x_{j,0} - x_i\| < L_{i,HSCB}$, where $L_{i,HSCB}$ is the length of the HSC branch of $i$. The choice of initial tangent direction and direction of extending segment of $j$ is described in the section of **Network generation approach**. One day after the injection of $CCl_4$, the rest half of aHSCs are switched to quiescence (if it is activated before, it is switched to reverted phenotype, according to the report that 50% of the aHSCs are switched to quiescence during fibrosis[32]).

**Degradation of collagen fibers**. In our model, we have two types of macrophages: resident and infiltrating macrophages. The resident macrophages are those in the liver while the infiltrating macrophages enter the liver after each injection dose. The life of infiltrating macrophages is 120 hours following previous study[9]. We observe that F4/80 positive macrophages only start to appear 12 hours after the injection of $CCl_4$. Therefore, we assume that both resident and infiltrating macrophages are initially F4/80 negative ($Ly6C^{high}$) phenotypes. They maintain the F4/80 negative ($Ly6C^{high}$) phenotype within 1-12 hours after the injection of $CCl_4$ and switch to a F4/80 positive ($Ly6C^{low}$) phenotype from 12 hours on after the injection. Macrophages with $Ly6C^{low}$ phenotype can revert aHSCs to quiescent

HSCs and digest collagen fibres. To mimic the digestion process, for each F4/80 positive (Ly6C$^{low}$) phenotype macrophage $i$, if the distance between $i$ and a collagen fiber segment $j$ satisfies $d_{ij} < r_i + r_j$, where $r_i$ is the radius of $i$ and $r_j$ is the radius of $j$, we remove $j$ from the system. Since a previous study has reported that the engulfment time one macrophage takes to digest an object is about 3 hours[34], we assume that a F4/80 positive (Ly6C$^{low}$) macrophage is only able to digest the next fiber 3 hours after it engulfs the previous fiber.

**Digestion of dead hepatocytes by macrophages**. Only CYP2E1+ hepatocytes die upon $CCl_4$-injection. As shown in **Supplementary Fig. 7B**, the AST level (partially correlated with the number of necrotic cells) after the second dose of $CCl_4$ is only 1/10 of the AST level after the first dose of $CCl_4$. As shown in **Fig. 2A** (HE & IgG staining), at the time point of the first dose of $CCl_4$, there are about 8 layers of hepatocytes dying due to intoxication (these are CYP2E1 positive hepatocytes before they are killed by a first dose of $CCl_4$). Under the hypothesis that the AST level correlates linearly with the number of necrotic hepatocytes, we assume that there are only two layers of CYP2E1+ - hepatocytes along the CV-CV-connection leading to two layers of necrotic hepatocytes in the middle of the lesion ($1/10 \times 8 \approx 1$) after the second dose of $CCl_4$ . The F4/80-positive macrophage can digest the necrotic hepatocytes. To mimic the digestion process, for a F4/80-positive macrophage $i$, if the distance between $i$ and a necrotic hepatocyte $j$ satisfies $d_{ij} < r_i + r_j$, where $r_i$ is the radius of $i$ and $r_j$ is the radius of $j$, we remove $j$ from the system. We also assume that the F4/80 positive macrophage is only able to digest a necrotic hepatocyte 3 hours after it engulfs a previous necrotic hepatocyte.

**Choosing parameters of hepatocyte death and proliferation in the simulation of biliary fibrosis**

As described in the main text, the heatmap of the spatial-temporal distribution of the bile ducts is measured by analyzing the staining of CK19 (marker for bile duct) in the Abcb4*KO* mice after 8, 16, and 28 weeks (3 mice in each case). The number of bile ducts and their distribution along the PV-PV axis over time are sampled according to the heatmap. The bile leaking from the bile ducts induce cell death in the surrounding hepatocytes. In our model, each bile duct is modeled as one sphere with the same size and physical property of hepatocyte (Young's modulus and Poisson ratio). We assumed the hepatocytes surrounding each bile duct would be chosen stochastically to be killed. Here is also one upper bound of the total number of killed hepatocytes. This upper bound of the fraction of dying hepatocytes is estimated by the expression level of AST, which is around 48, 198, 161, 286 U/l at control and after 8, 16, 28 weeks, respectively. In our septal fibrosis simulation, the AST level at control and under repeated injection of $CCl_4$ is around 50 and 200 U/l, respectively. Since AST level at 200 U/l in septal fibrosis case corresponds to around 4% of hepatocyte death, here we assumed similar relationship of AST level and hepatocyte death fraction in the biliary fibrosis case, where the hepatocyte death fraction would be no more than 4%, 3.2%, 6% after 8, 16, 28 weeks, respectively.

In addition to hepatocyte death, there is also hepatocyte proliferation. The liver/weight ratio over time is 6.3±0.1%, 6.0±0.1%, 6.5±0.3% after 8, 16, 28 weeks, respectively. Since there is no significant change in the liver/weight ratio (all around 6%), we assume that the number of newly proliferating hepatocytes equals the number of dead hepatocytes. Therefore, the liver can still maintain its total mass over time.

**Constructing the heatmap of spatial-temporal distribution of bile ducts**

In the staining images of bile ducts (as shown in **Fig. 6B**), each pair of PV-PV is taken out and one line is added to represent the PV-PV axis. The neighboring region surrounding the PV-PV axis is divided into subregions. For each bile duct, its distance to the PV-PV axis and its relative position along the PV-PV axis were calculated to determine it was located in which subregion. The numbers of bile ducts in all subregions were normalized to count the heatmap of probability of bile duct distribution according to their relative position to the PV-PV axis.

**Perturbation simulation of components of the DT**

In addition to the 2 perturbation tests shown in **Fig. 5A**, we also conduct *in silico* perturbations targeting hepatocyte proliferation, HSC activation, and Mph phagocytic activity, comparing the outcomes with the reference DT model (**Supplementary Fig. 11A**). To assess the impact of biomechanical pressure from dividing hepatocytes on collagen organization, we run the DT simulation by excluding hepatocyte proliferation for 21 days **(P3**). The model predicts that fewer hepatocytes would be present in the lesion compared to the reference model. Interestingly, the same amount of deposited ECM in the injured regions fails to form a fibrotic wall, as observed in the reference model after 6 doses of $CCl_4$. The collagen network diameter is much larger compared to the reference case (**P3; Supplementary Fig. 11B, C, Supplementary Movie 6**). The relationship suggesting that fewer hepatocytes lead to a wider ECM network supports the conclusion that hepatocyte division is a significant driver of the experimentally observed septal fibrosis pattern, likely through biomechanically displacing and compressing the ECM fibers, which is absent in absence of hepatocyte proliferation (**Supplementary Fig. 10B, C, D**).

In another scenario, we investigate the influence of HSC phenotypes and activities on fibrosis pattern development and shaping. We conduct three independent perturbations. In the reference model, 50% of aHSCs undergo apoptosis while 50% revert to a quiescent state during regeneration (as determined in previous experimental studies[32]). In one perturbation, aHSC migration is disabled (**P4**), and in the other perturbation, HSCs are initiated as inverted phenotype (**P5**). These perturbations are simulated with the DT for 3 weeks, involving 6 $CCl_4$ injections. In **P4**, there are fewer collagen fibers distributed along the lesion boundary, because HSC lacking migratory activity remains in their original position (**P4, Supplementary Fig. 11C; Supplementary Movie 7**).

As a consequence, fewer HSCs accumulate in the center of the lesion compared to the reference. In **P5**, collagen fiber deposition begins earlier compared to the reference model, around day 1.5 versus day 3 (**Supplementary Fig. 11B, Supplementary Movie 8**). This occurs because reverted HSCs are more responsive to DAMP-mediated activation. In summary, the HSC-directed model perturbations suggest that the HSC phenotype primarily impacts the absolute amount of ECM deposition, while HSC migration is crucial for the formation of the septal fibrosis pattern.

Furthermore, we assess the phagocytic and migratory effects of Mph and conducted two separate Mph related perturbations with the DT. Mphs are transitioned to Ly6C$^{low}$ after 5 days following each $CCl_4$ injection, instead of the 12 hours as evident in the reference model (**P6**), or they are assumed to be deficient in migration ability (**P7; Supplementary Fig. 11A**). Simulation P6 results in increased collagen fiber presence in the injured regions without altering the wall structure formation pattern (**Supplementary Fig. 11C; Supplementary Movie 9**). Simulating Mph migration deficiency (**P7**) leads to the formation of a looser and more broadly distributed collagen fiber network (**Supplementary Fig. 11C, Supplementary Movie 10**). Furthermore, the amount of deposited collagen increases, because the number of Mphs available in the lesion to degrade collagen fibers is decreased (**Supplementary Fig. 11C**). The gyration radius of the collagen network in simulations **P6** and **P7** is slightly larger than that in the reference model (**Supplementary Fig. 11**). Perturbations **P6** and **P7** have no impact on the number of hepatocytes in the lesion (**Supplementary Fig. 11B**). The perturbation results indicate that Mph activity has a minor effect on the development of the septal fibrosis pattern.

Finally, we assess the effects of hepatocyte pressure on collagen and the existence of sinusoids, therefore, we conducted two separate perturbations with the DT. Hepatocyte-collagen contact is disabled (**P8; Supplementary Fig. 11A**) to check the effect of the biomechanical pressure from proliferating hepatocytes on the collagen network and sinusoids are removed (**P9; Supplementary Fig. 11A**). Simulation P8 results in a much larger gyration diameter of the collagen network (**Supplementary Fig. 11B and C; Supplementary Movie 11**). This indicates that biomechanics of the cell-collagen contact plays significant role in shaping the fibrosis pattern. Simulation P9 results in a little bit increase of the gyration diameter of the collagen network (**Supplementary Fig. 11B and C; Supplementary Movie 12**). This partially indicates that the presence of the sinusoids play a geometric role to regulate the movement of the hepatocytes in shaping the fibrosis pattern.

**Boolean network model**

When we performed the perturbation test to predict the outcome of a drug treatment on the activity of FAK (focal adhesion kinase) on HSC, we applied a Boolean network model (Montagud et al.[35]) to model the probability of HSC activation upon activation or inactivation of FAK over time. As shown in **Fig. 5G**, a simplified intracellular network regarding FAK-promoted HSC activation was used for the Boolean network. In this network, TGFβ binds to the TGFβ receptor to trigger the activation of SMAD, which then triggers the HSC activation. The presence of FAK promotes the activity of TGFβ. In the Boolean network, we assigned two states for each of these species: 1 for presence and 0 for absence. For example, if FAK is present to bind TGFβ receptor, the state of FAK is 1, otherwise if FAK is absent to bind TGFβ receptor, the state of FAK is 0. For the species TGFβ receptor, SMAD and HSC activation, each has two rates: rate of upregulation and rate of downregulation. The values of the two rates of each of these species (TGFβ receptor, SMAD and HSC activation) would be determined by its binding or triggering partner. The state of each of these species (TGFβ receptor, SMAD and HSC activation) would be determined stochastically according to its two rates. For example, the rates of upregulation and downregulation of SMAD would be determined by the state of TGFβ receptor. The state of SMAD would then be determined stochastically by the rates of upregulation and downregulation of SMAD. Here we provide the pseudo code of our Boolean network model (Algorithm 0) below:

**Algorithm 0**. The pseudo code of the Boolean network model:

---

1 {***Determine the state of one species stochastically according to its rates of upregulation and downregulation***}:
2 For species $S$ and its rates of upregulation $rate_{up,S}$ and downregulation $rate_{down,S}$
3     generate one random number $r \sim U(0,1)$
4     if state $S == 0$
5         if $r < rate_{up,S}$
6             state $S = 1$
7         else
8             state $S = 0$
9         end if
10     else
11         if $r < rate_{down,S}$
12             state $S = 0$
13         else
14             state $S = 1$
15         end if
16     end if
17 {***Initiate the states of all species and simulation time***}:
18 Initiate
19     $TGF\beta = 1, TGF\beta R = 0, SMAD = 0, HScActivation = 0, FAK = 0|1$
20     Starting time $t = t_0$
21 {***Main loop***}:
22     repeat
23         {***Determine the rates of upregulation and downregulation of TGF$\beta$ receptor according to the state of FAK***}:
24         if state $FAK == 0$
25             $rate_{up,TGF\beta} = rate_{up,TGF\beta,0}$
26             $rate_{down,TGF\beta} = rate_{down,TGF\beta,0}$
27         else
28             $rate_{up,TGF\beta R} = rate_{up,TGF\beta R,1}$
29             $rate_{down,TGF\beta R} = rate_{down,TGF\beta R,1}$
30         end if
31         {***Determine the state of TGF$\beta$ receptor stochastically***}:
32         determine the state of TGF$\beta$R → go to line 1-16
33         {***Determine the rates of upregulation and downregulation of SMAD according to the state of TGF$\beta$ receptor***}:
34         if state $TGF\beta R == 0$
35             $rate_{up,SMAD} = rate_{up,SMAD,0}$
36             $rate_{down,SMAD} = rate_{down,SMAD,0}$
37         else
38             $rate_{up,SMAD} = rate_{up,SMAD,1}$
39             $rate_{down,SMAD} = rate_{down,SMAD,1}$
40         end if
41         {***Determine the state of SMAD stochastically***}:
42         determine the state of SMAD → go to line 1-16
43         {***Determine the rates of upregulation and downregulation of HSC activation according to the state of SMAD***}:
44         if state $SMAD == 0$
45             $rate_{up,HSCActivation} = rate_{up,HSCActivation,0}$
46             $rate_{down,HSCActivation} = rate_{down,HSCActivation,0}$
47         else
48             $rate_{up,HSCActivation} = rate_{up,HSCActivation,1}$
49             $rate_{down,HSCActivation} = rate_{down,HSCActivation,1}$
50         end if
51         {***Determine the state of HSC activation stochastically***}:
52         determine the state of HSC activation → go to line 1-16
53         {***Update the time***}:
54         $t = t + \Delta t$
55     until $t > t_{end}$

---

**Sub-section** (**lines 1 to 16**): Determine the state of one species according to its rates of upregulation and downregulation stochastically.

**Initiate the states of all species** (**lines 17 to 20**): TGFβ=1,TGFβR=0,SMAD=0,HSCActivation=0, FAK=0 or 1.

**Main loop** (**lines 21 to 55**): A step-by-step time evolution is proceeded within a loop. The following steps are repeated until the system reaches the end time $t_{end}$.

**Step 0** (**lines 23 to 30**): Determine the rates of upregulation and downregulation of TGFβR according to the state of FAK.

**Step 1** (**lines 31 to 32**): Using sub-section (lines 1 to 16) to determine the state of TGFβR.

**Step 2** (**lines 33 to 40**): Determine the rates of upregulation and downregulation of SMAD according to the state of TGFβR.

**Step 3** (**lines 41 to 42**): Using sub-section (lines 1 to 16) to determine the state of SMAD.

**Step 4** (**lines 43 to 50**): Determine the rates of upregulation and downregulation of HSCActivation according to the state of SMAD.

**Step 5** (**lines 51 to 52**): Using sub-section (lines 1 to 16) to determine the state of HSCActivation.

**Step 6** (**lines 53 to 54** – Update time): Increase t by Δt.

**Computational scheme executed**

The simulations of the DT were performed using TiSim with customized additions. The formal version of TiSim will be released in the near future.

In every time step, the new positions of all cells and elements (collagen fibers and sinusoids) are updated by solving the force balance equation, given the forces applied on them. The interaction force either between objects either from the different entities (e.g. the adhesive and repulsive force between two hepatocytes or between hepatocyte and sinusoidal sphere, equation (2)) or from the same entity (e.g. the spring force between two sinusoidal spheres in the same sinusoidal edge, $\vec{F}_{i,ela}$ in equation (6)) are based on the overlap of the two objects which is identified by collision detection. The system of force balance equation is a large linear system with size increased as more elements are added into the system (e.g. the production of collagen fibers). This system can be solved efficiently by using a conjugate gradient method. The signaling dynamic is updated by solving corresponding PDE of each signal using finite difference method. The time for a full round of simulation takes approximately 720 hours on a workstation (Intel(R) Xeon(R) Gold 6136 CPU).

The implementation of the above computational scheme is shown and described in the following flowcharts and corresponding piece of pseudo codes.

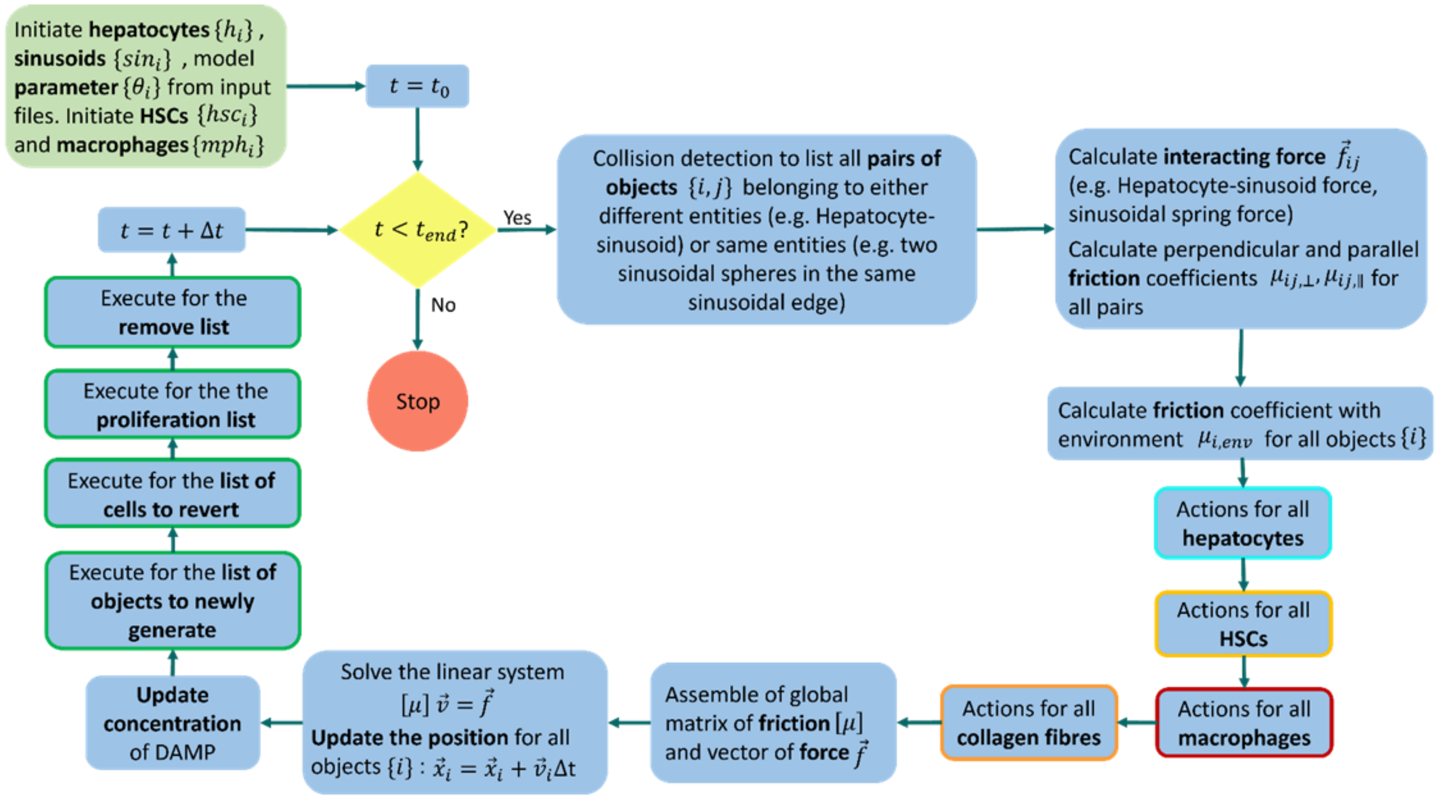

**Flowchart of the main loop of the scheme execution**. Schematic representation of the main loop of our code implementation of the liver fibrotic pattern formation process. The details of implementation of actions for all hepatocytes, HSCs, macrophages and collagen fibers and executes for the list of objects to newly generate, the list of cells to revert, the proliferation list, and the remove list are represented in individual flowcharts below.

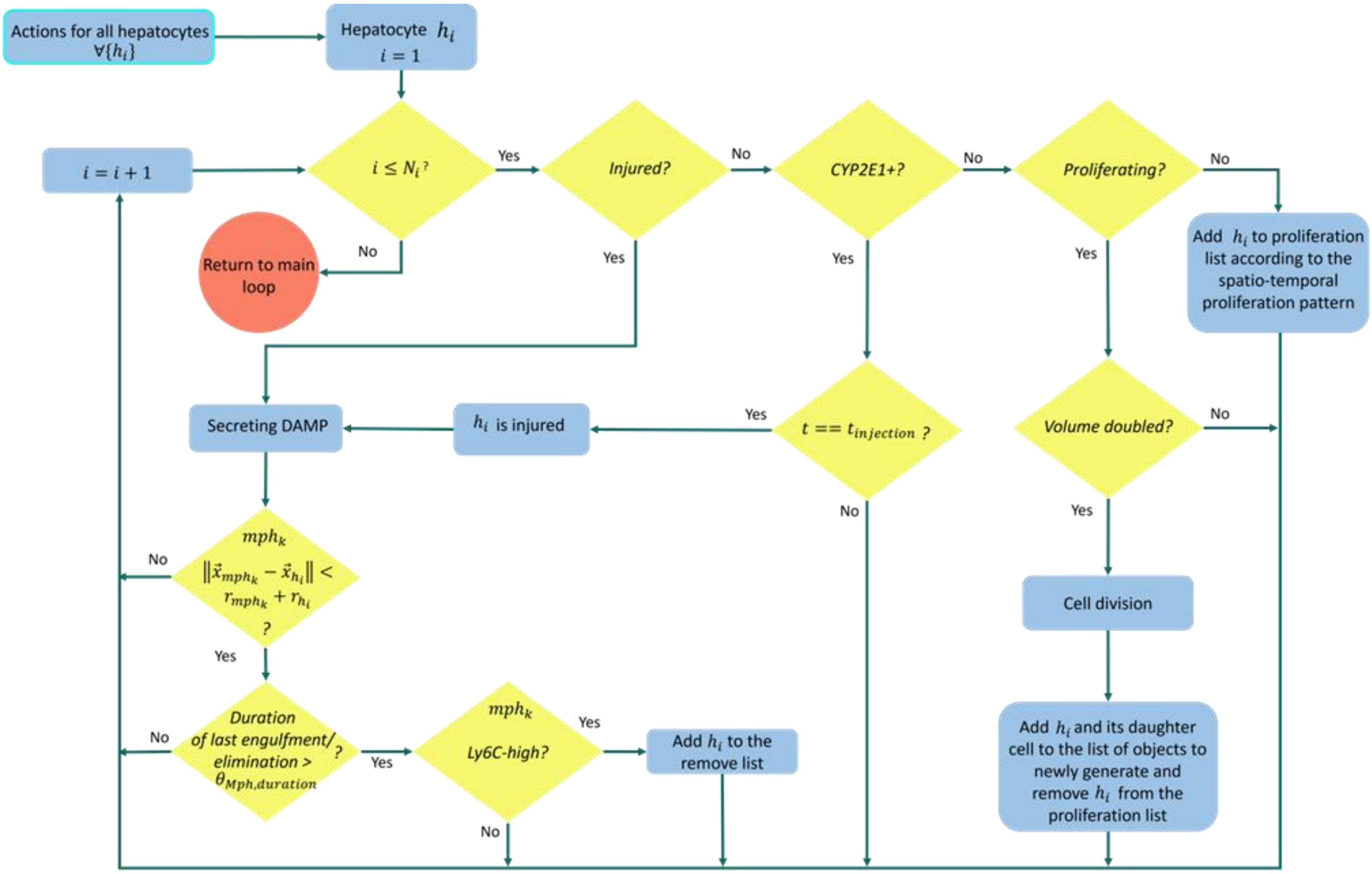


**Flowchart of the implementation of actions for all hepatocytes.**

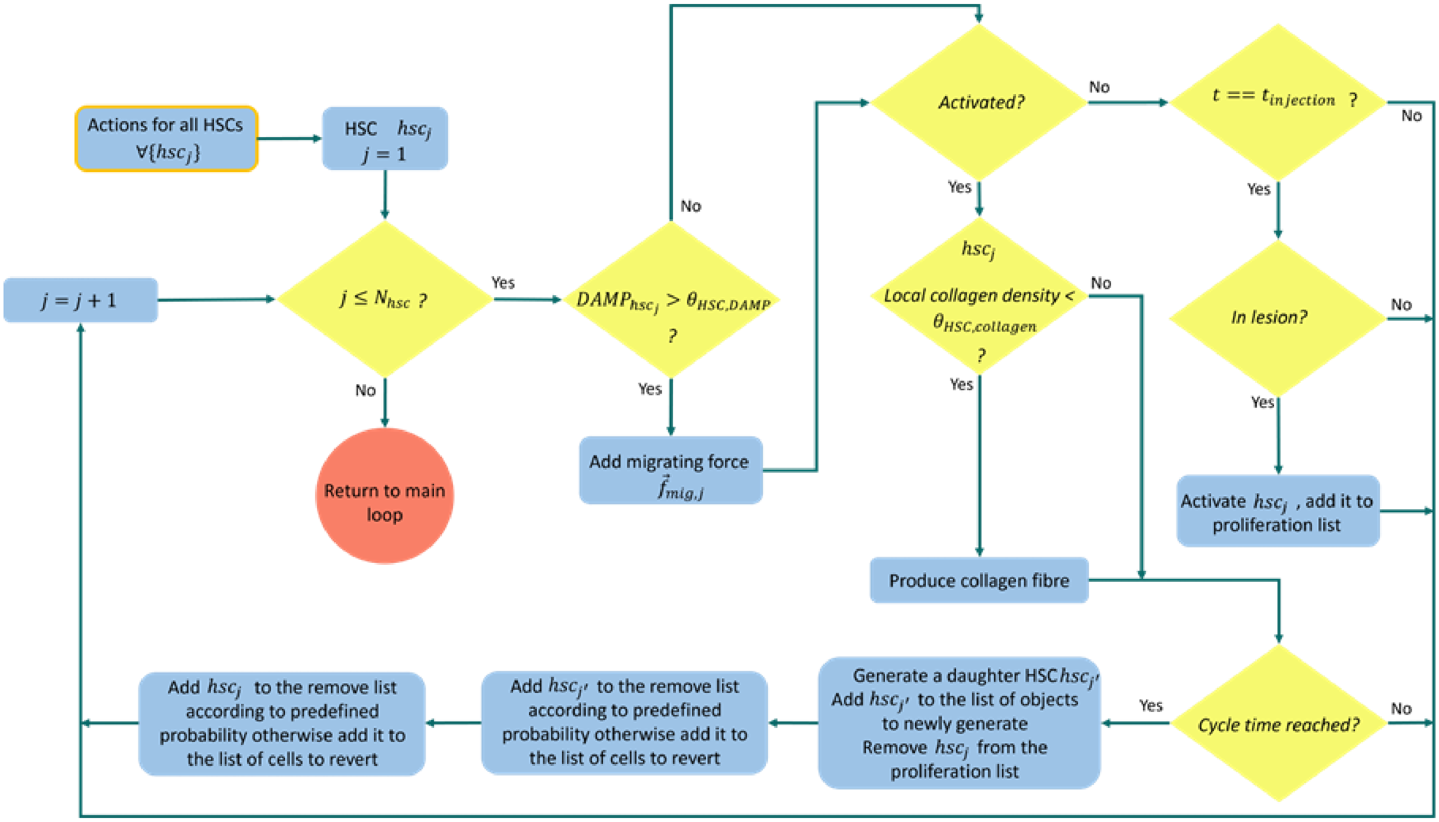


**Flowchart of the implementation of actions for all HSCs. The fate decisions of HSCs after cell division were implemented to mimic the observed conservation of HSC population size in ref.[24] (The precise mechanism of population size conservation is not expected to influence the results.)**

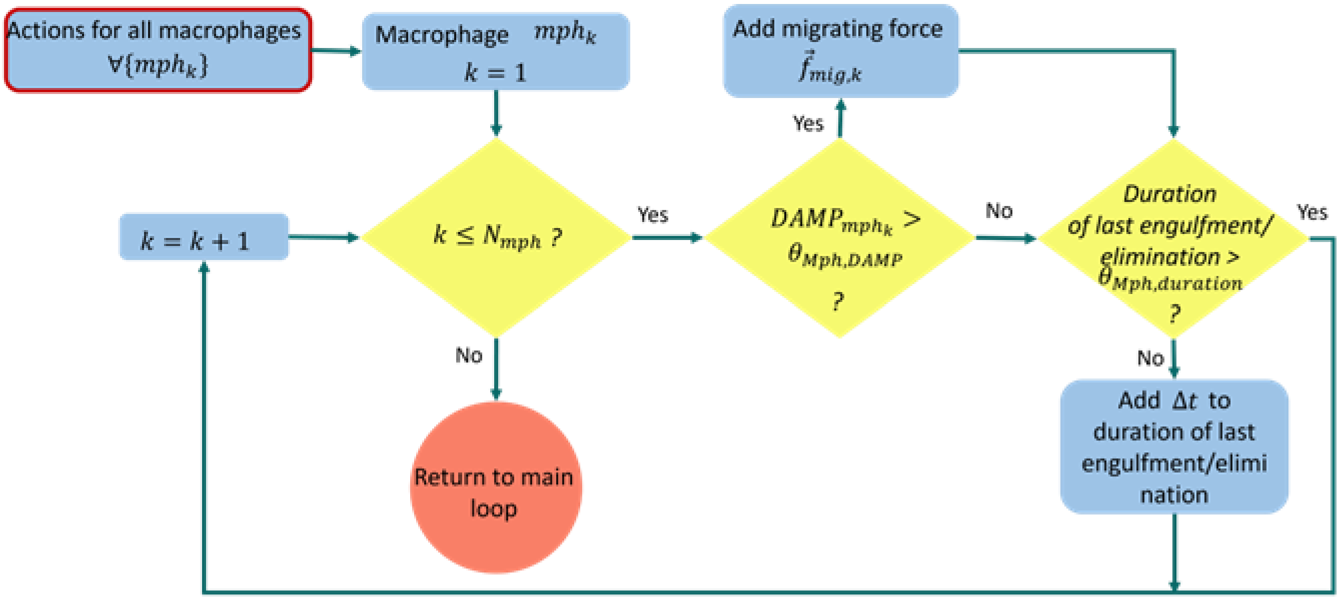


**Flowchart of the implementation of actions for all macrophages.**

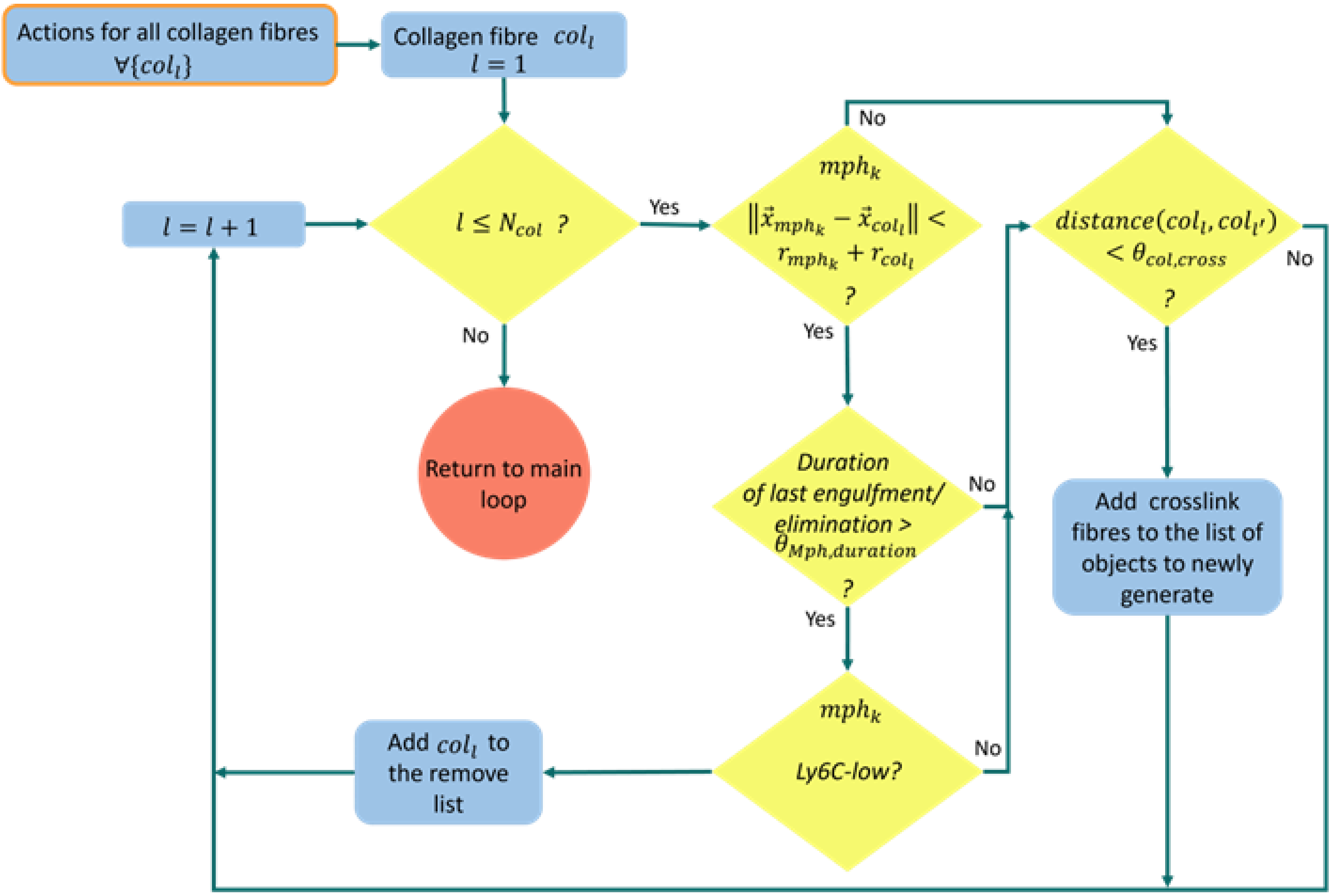

**Flowchart of the implementation of actions for all collagen fibers.**

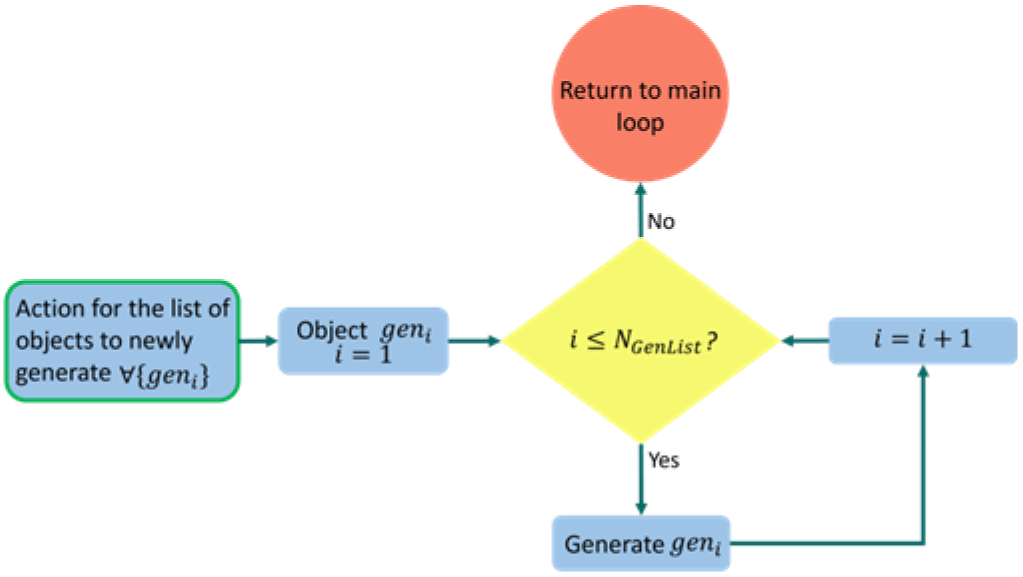


**Flowchart of the implementation of execute for the list of objects to newly generate.**

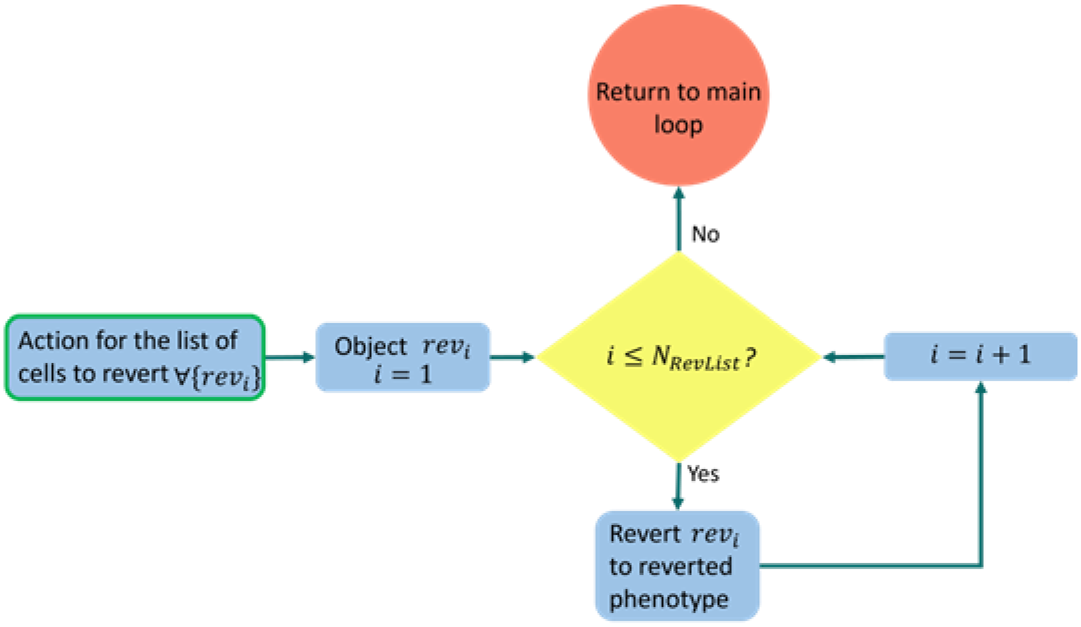


**Flowchart of the implementation of execute for the list of cells to revert.**

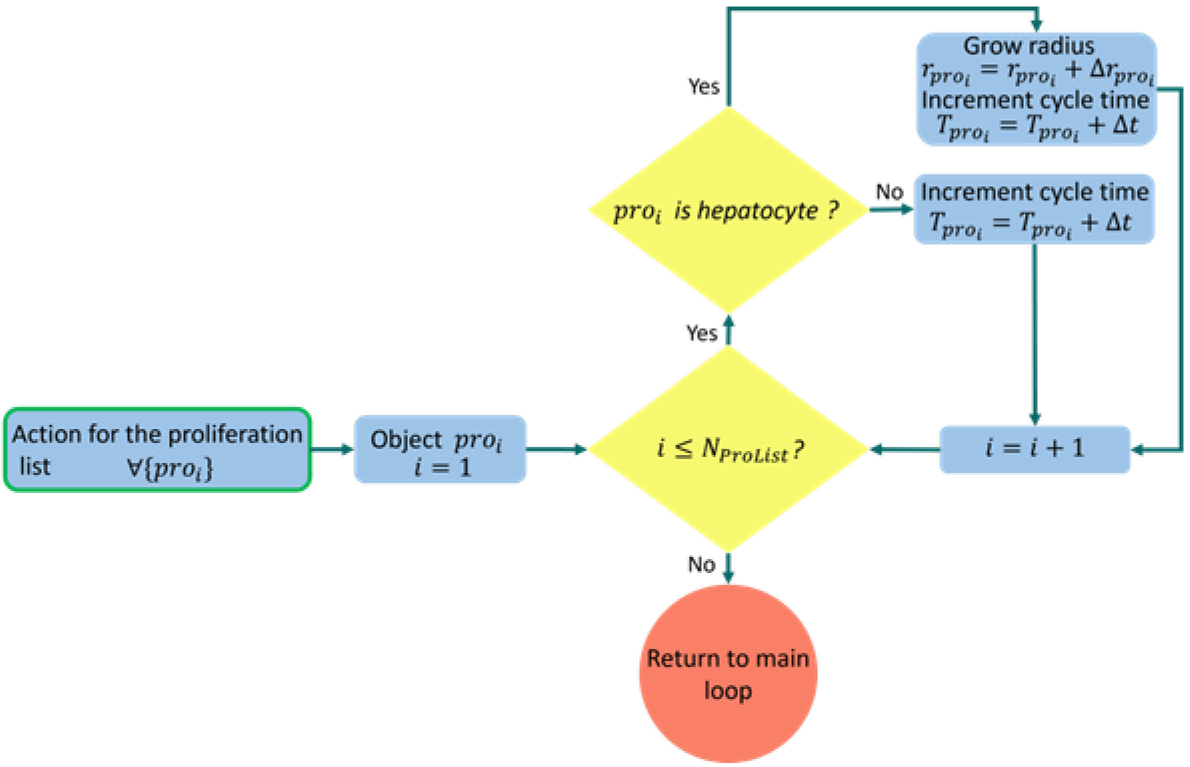


**Flowchart of the implementation of execute for the proliferation list.**

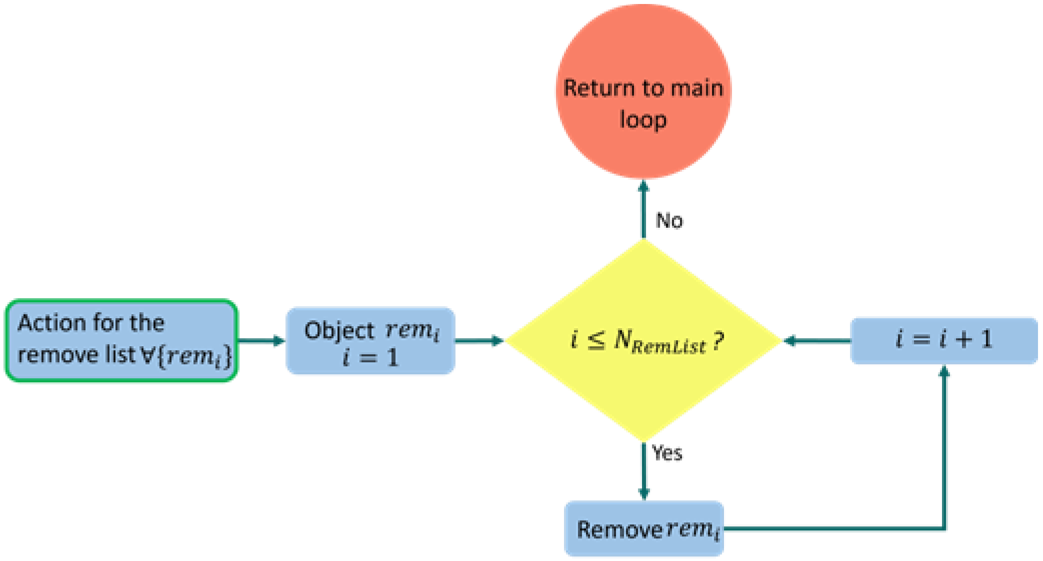


**Flowchart of the implementation of execute for the remove list.**

**Algorithm 1**. The pseudo code of the main loop:

---

1 ***{Read the structures of hepatocytes and sinusoids:}***
2 Initiate the
3 set of sinusoids $SinList \coloneqq \{sin_i, sin_i.objectType = Sinusoid | i: sinusoid\ sphere\ index\}$
4 set of hepatocytes $HepList \coloneqq \{h_i, h_i.objectType = Hepatocyte, h_i.daughter = FALSE | i: hepatocyte\ index\}$
5 ***{Read model parameters:}***
6 Initiate the set of parameters $\Theta List \coloneqq \{\theta_i | i: parameter\ index\}$
7 ***{Initiate HSCs and macrophages:}***
8 Initiate the
9 set of HSCs $HSCList \coloneqq \{hsc_i, hsc_i.objectType = HSC | i: HSC\ index\}$
10 set of macrophages $MphList \coloneqq \{mph_i, mph_i.objectType = Macrophage | i: macrophage\ index\}$
11 ***{Initiate other data sets and simulation time:}***
12 Initiate the
13 set of collagen fibres $ColList \coloneqq \{col_i, col_i.objectType = Collagen, col_i.toRemove = FALSE | i: collagen\ fibre\ index\}$ as empty: $Col \coloneqq \emptyset$
14 set of objects $ObjList \coloneqq \{obj_i | obj_i \in SinList \parallel obj_i \in HepList \parallel obj_i \in HSCList \parallel obj_i \in MphList \parallel obj_i \in ColList\}$
15 set of pairs of interacting objects $PairList \coloneqq \{(i,j) | i,j \in ObjList\}$
16 list of objects to newly generate $GenList \coloneqq \{gen_i | gen_i \in ObjList\}$ as empty: $GenList \coloneqq \emptyset$
17 list of cells to revert $RevList \coloneqq \{rev_i | rev_i \in HSCList\}$ as empty: $RevList \coloneqq \emptyset$
18 proliferation list $ProList \coloneqq \{pro_i | pro_i \in HepList \parallel pro_i \in HSCList\}$ as empty: $ProList \coloneqq \emptyset$
19 remove list $RemList \coloneqq \{rem_i | rem_i \in ObjList\}$ as empty: $RemList \coloneqq \emptyset$
20 starting time: $t := t_0$
21 repeat
22 ***{perform collision detection to update the set of pairs of interacting objects and calculate the interacting force and perpendicular and parallel friction coefficients:}***
23 for all $(i,j) \in PairList$
24 calculate $\vec{f}_{ij}$
25 calculate $\mu_{ij,\perp}$ and $\mu_{ij,\parallel}$
26 end for
27 ***{calculate the friction with the environment:}***
28 for all $obj_i \in ObjList$
29 calculate $\mu_{i,env}$
30 end for
31 ***{actions for different cells and collagen fibres:}***
32 actions for
33 $HepList \coloneqq \{h_i\}$ → **Algorithm 2**
34 $HSCList \coloneqq \{hsc_j\}$ → **Algorithm 3**
35 $MphList \coloneqq \{mph_k\}$ → **Algorithm 4**
36 $ColList \coloneqq \{col_l\}$ → **Algorithm 5**
37 ***{assemble friction matrix and force vector:}***
38 assemble the global matrix of friction: $[\mu]$, the vector of force: $\vec{f}$
39 ***{solve the linear system and update the position:}***
40 solve $[\mu]\vec{v} = \vec{f}$
41 for all $obj_i \in ObjList$
42 $\vec{x}_i := \vec{x}_i + \vec{v}_i \Delta t$
43 end for
44 ***{update the DAMP concentration:}***
45 solve eqn(16)
46 ***{executes for the list of objects to newly generate, the list of cells to revert, the proliferation list, and the remove list:}***
47 executes for *GenList*, *RevList*, *ProList*, *and* *RemList* → **Algorithm 6**
48 ***{update the time:}***
49 $t := t + \Delta t$
50 until $t > t_{end}$

---

**Step 0** (**lines 1 to 20 – Read the input files to initiate the data sets and variables for simulation**):

Read the input lobule architecture file for the spatial positions of sinusoids and hepatocytes to initiate the sets of sinusoids $SinList \coloneqq \{sin_i\}$ and hepatocytes $HepList \coloneqq \{h_i\}$. For any non-injured hepatocyte, if its distance to the nearest lesion boundary is smaller than 4 hepatocytes' diameters, it is modeled using DCM otherwise it is modeled using CBM. The injured hepatocytes upon the first injection of $CCl_4$ are modeled using CBM while the injured hepatocytes from the second injection of $CCl_4$ are modeled using DCM (because there is no collagen generated after the first injection of $CCl_4$, the high-resolution model of mechanical contact between collagen and hepatocyte is therefore not needed during that stage). The set of HSCs $HSCList \coloneqq \{hsc_i\}$ and macrophages $MphList \coloneqq \{mph_i\}$ are initiated in the space of simulation according to their corresponding cell density (see more details in Zhao et al.[6]). The set of collagen fibres is initiated as empty. There is one concatenated set of all these cells and elements as a set of objects $ObjList \coloneqq \{obj_i\}$. There is a set of all pairs of interacting objects $PairList \coloneqq \{(i,j)|i,j \in ObjList\}$, which will be updated each time step by collision detection. There are four additional sets recording cells or objects with morphological change during the simulation, all initiated as empty: the list of objects to newly generate $GenList \coloneqq \{gen_i|gen_i \in ObjList\}$, which records the objects to be generated into the system; the list of cells to revert $RevList \coloneqq \{rev_i|rev_i \in HSCList\}$, which records the cells to be reverted to inverted phenotype; the proliferation list $Pro \coloneqq \{pro_i|pro_i \in HepList \ \| \ pro_i \in HSCList\}$, which records the proliferating cells; the remove list $RemList \coloneqq \{rem_i|rem_i \in ObjList\}$, which records the objects to be removed from the system. The executes of these four lists are performed individually after the position of all objects are updated during each time step.

**Steps 1 to 7** (**lines 21 to 50**): A step-by-step time evolution is proceeded within a loop. The following steps are repeated until the system reaches the end time $t_{end}$.

**Step 1** (**lines 22 to 26 – Calculate the interacting force and friction coefficients**): Perform the collision detection to update the set of interacting objects $PairList \coloneqq \{(i,j)|i,j \in ObjList\}$, where $i$ and $j$ are two interacting objects from either different entities (e.g. two interacting hepatocytes or hepatocyte-sinusoid interaction) or the same entity (e.g. two sinusoid spheres from the same sinusoid edge). For all interacting pairs $(i,j) \in PairList$, calculate the interacting force $\vec{f}_{ij}$ and the perpendicular and parallel friction coefficients $\mu_{ij,\perp}$ and $\mu_{ij,\parallel}$.

**Step 2** (**lines 27 to 30 – Calculate the friction coefficient with the environment**): For all objects $obj_i \in ObjList$, calculate its friction coefficient with the environment $\mu_{i,env}$.

**Step 3** (**lines 31 to 36 – Actions for cells and collagen fibres**): For different types of cells (hepatocytes, HSCs and macrophages) and collagen fibres, their behaviors and fates are modeled individually in each section of actions. See **Algorithms 2 to 5** for details.

**Step 4** (**lines 37 to 43 – Update the positions for all objects**): Assemble the global matrix of friction coefficients $[\mu]$ and vector of forces $\vec{f}$ of all objects . Solve the linear system $[\mu]\vec{v} = \vec{f}$ to obtain the velocity vector of all objects. The new positions of all objects $obj_i \in ObjList$ are updated as $\vec{x}_i := \vec{x}_i + \vec{v}_i\Delta t$.

**Step 5** (**lines 44 to 45 – Update the DAMP concentration**): Solve the PDE (eqn(16)) to update the concentration of DAMP (see more details in Zhao et al.[6]).

**Step 6** (**lines 46 to 47 – Executes for the list of objects to newly generate, the list of cells to revert, the proliferation list, and the remove list**): Executes for objects and cells in the lists $GenList$, $RevList$, $ProList$, and $RemList$. See **Algorithm 6** for details.

**Step 7** (**lines 48 to 49 – Update time**): Increase $t$ by $\Delta t$.

**Algorithm 2**. The pseudo code of actions for all hepatocytes:

```
$\forall h_i \in HepList$:  $i := 1$
repeat
   if $!h_i.injured$
      if $h_i.CYP2E1+$
         if $t == t_{injection}$
            $h_i.injured = TRUE$
         end if
      else
          if $h_i.proliferating$
             if $h_i.volume > 2 \times h_i.initialVolume$
                remove $h_i$ from $ProList$
                create daughter cell: $h_{i'}$
                $h_{i'}.daughter = TRUE$
                add $h_i$ and $h_{i'}$ to $GenList$
             end if
          else
             generate $u_1$ from $U(0,1)$
             if $u_1 < \theta_{Hep,proliferation}(\vec{x}_{h_{i'}}, t)$
                add $h_i$ to $ProList$
             end if
          end if
       end if
   end if
   if $h_i.injured$
      $h_i.secretingDAMP = TRUE$
      if $mph_k$ satisfying $\left\|\vec{x}_{mph_k} - \vec{x}_{h_i}\right\| < r_{mph_k} + r_{h_i}$
         if $mph_k.durationEngulfmentElimination > \theta_{Mph,duration}$
            if $mph_k.Ly6CHigh$
              add $h_i$ to $RemList$
            end if
         end if
      end if
   end if
   $i = i + 1$
until $i > N_h$
```

**Algorithm 3**. The pseudo code of actions for all HSCs:

---

```
∀$hsc_j \in HSCList$: $j := 1$
repeat
   if $DAMP_{hsc_j} > \theta_{HSC,DAMP}$
      $\vec{f}_{hsc_j} := \vec{f}_{hsc_j} + \vec{f}_{mig,j}$
   end if
   if $hsc_j.activated$
      if $hsc_j.localCollagenDensity < \theta_{HSC,collagen}$
         $ProduceCollagen(hsc_j)$
      end if
      if $hsc_j.reachCycleTime$
         remove $hsc_j$ from $ProList$
         generate a daughter HSC $hsc_{j'}$
         add $hsc_{j'}$ to $GenList$
         generate $u_1$ from $U(0,1)$
         if $u_1 < \theta_{HSC,remove}$
            add $hsc_j$ to $RemList$
         else
            add $hsc_j$ to $RevList$
         end if
         generate $u_2$ from $U(0,1)$
         if $u_2 < \theta_{HSC,remove}$
            add $hsc_{j'}$ to $RemList$
         else
            add $hsc_{j'}$ to $RevList$
         end if
      end if
   else
      if $t == t_{injection}$
         if $hsc_j.inLesion$
            $hsc_j.activated = TRUE$
            add $hsc_j$ to the $ProList$
         end if
      end if
   end if
   $j = j + 1$
until $j > N_{hsc}$
```

---

**Algorithm 4**. The pseudo code of actions for all macrophages:

```
$\forall mph_k \in MphList$:  $k := 1$
repeat
   if $DAMP_{mph_j} > \theta_{Mph,DAMP}$
      $\vec{f}_{mph_k} \coloneqq \vec{f}_{mph_k} + \vec{f}_{mig,k}$
   end if
   if $mph_k.durationEngulfmentElimination < \theta_{Mph,duration}$
      $mph_k.durationEngulfmentElimination \coloneqq mph_k.durationEngulfmentElimination + \Delta t$
   end if
   $k = k + 1$
until $k > N_{mph}$
```

**Algorithm 5**. The pseudo code of actions for all collagen fibres:

```
$\forall col_l \in ColList$:  $l := 1$
repeat
   if $mph_k$ satisfying $\left\|\vec{x}_{mph_k} - \vec{x}_{col_l}\right\| < r_{mph_k} + r_{col_l}$
      if $mph_k.durationEngulfmentElimination < \theta_{Mph,duration}$
         if $mph_k.Ly6CLow$
            $col_l.toRemove = TRUE$
         end if
      end if
   end if
   if $!col_l.toRemove$
      if $col_{l'}$ satisfying $distance(col_l, col_{l'}) < \theta_{col,cross}$
         add $crosslinkCollagenFibres(col_l, col_{l'})$ to $GenList$
      end if
   else
      add $col_l$ to $RemList$
   end if
   $l = l + 1$
until $l > N_{col}$
```

**Algorithm 6**. The pseudo code of actions for the list of objects to newly generate, the list of cells to revert, the proliferation, and the remove list:

---

```
∀gen_i ∈ GenList: i := 1
repeat
   if gen_i.objectType == Hepatocyte
      if gen_i.daughter
         add gen_i to HepList
         add gen_i to ObjList
         gen_i.daughter = FALSE
      else
         update states of gen_i
      end if
   else if gen_i.objectType == Collagen
      add gen_i to ColList
      add gen_i to ObjList
      gen_i.toRemove = FALSE
   else
      add gen_i to HSCList
      add gen_i to ObjList
   end if
   i = i + 1
until i > N_GenList
∀rev_i ∈ RevList, i := 1
repeat
   rev_i.activated = FALSE
   rev_i.reverted = TRUE
   i = i + 1
until i > N_RevList
∀pro_i ∈ ProList: i := 1
repeat
   if pro_i.objectType == Hepatocyte
      r_pro_i = r_pro_i + Δr_pro_i
      T_pro_i = T_pro_i + Δt
   else
      T_pro_i = T_pro_i + Δt
   end if
   i = i + 1
until i > N_ProList
∀rem_i ∈ RemList: i := 1
repeat
   if rem_i.objectType == Hepatocyte
      remove rem_i from HepList
   else if rem_i.objectType == HSC
      remove rem_i from HSCList
   else
      remove rem_i from ColList
   end if
   delete rem_i from ObjList
   i = i + 1
until i > N_RemList
clear RemList, GenList, RevList
```

---

**Supplementary Figures**

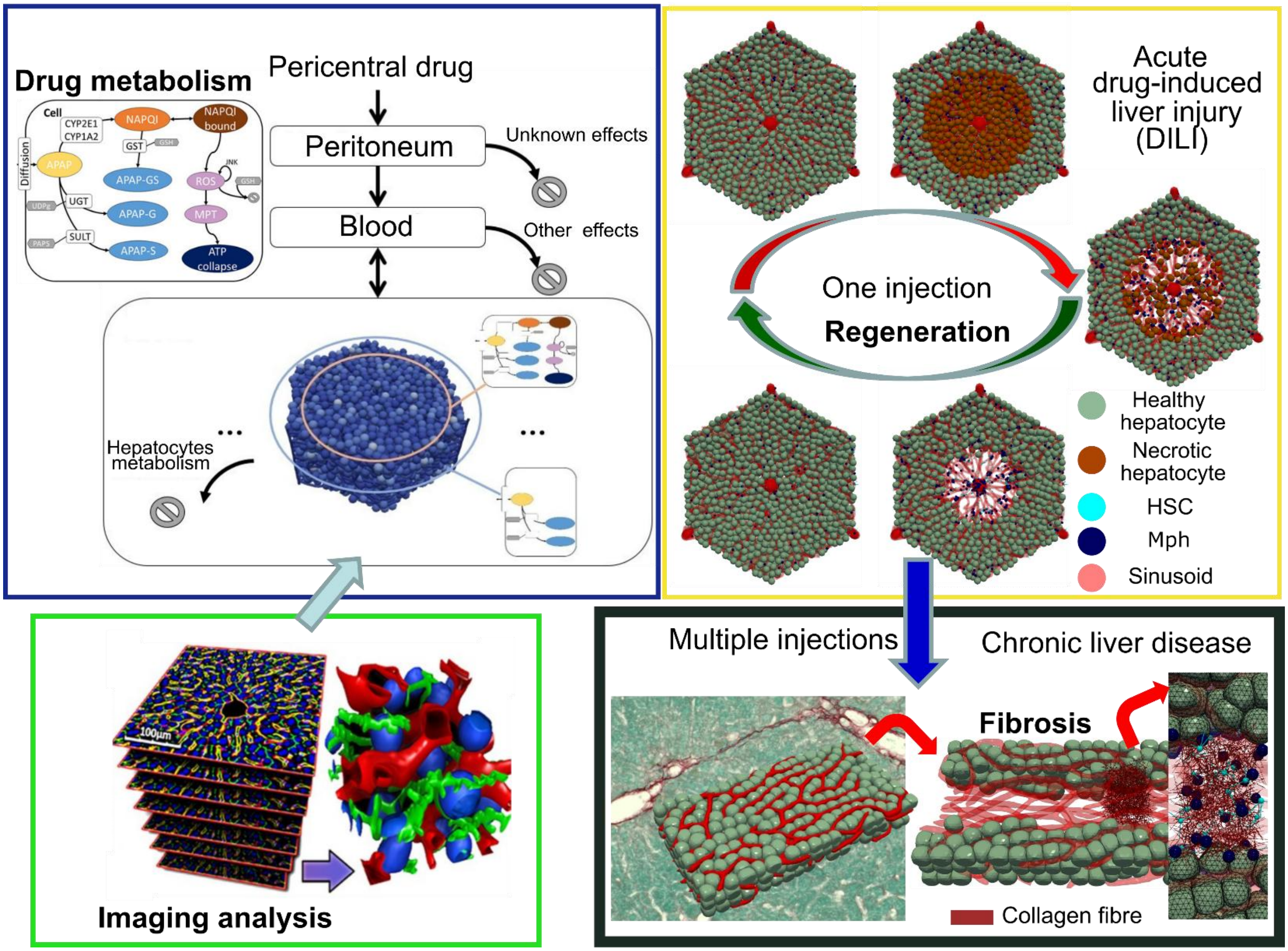


**Supplementary Figure 1. Scheme of the digital twin concept in studying liver related disease.** Construction of the liver microarchitecture of the digital twin from imaging analysis (green box, Hoehme et al.[7] (2010)). Modeling the drug metabolism in the liver cell (blue box, Dichamp et al.[36] (2023)). Modeling the liver regeneration upon one injection of drug-induced injury (yellow box, Zhao et al.[9] (2024)). Modeling septal liver fibrosis (black box, the focus of this study). (The PV-PV-bridging biliary fibrosis found in Abcb4-*KO*-mice and onion-shaped fibrosis found in humans simulated in this study as well are not shown here.)

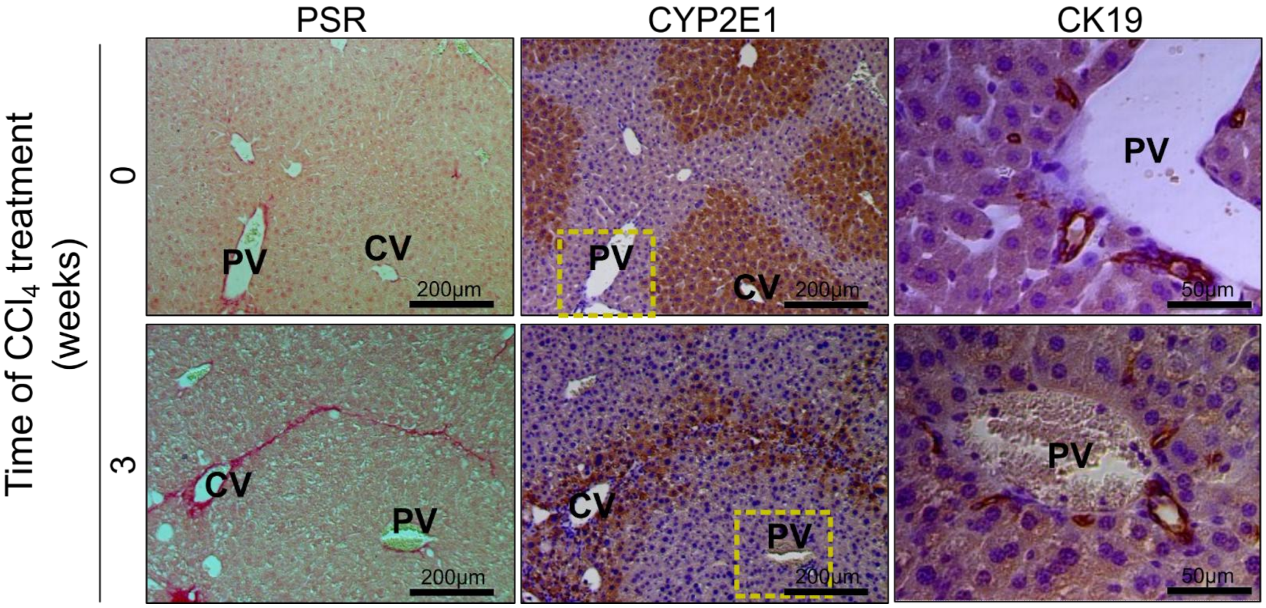


**Supplementary Figure 2. CYP2E1 expressing hepatocytes guide ECM deposition upon repetitive $CCl_4$ injections.** Serial sections from control and fibrotic mouse liver (3 weeks of $CCl_4$) are stained with PSR (ECM deposition; red), CYP2E1 (CCl4 metabolizing enzyme; brown) and CK19 (bile duct epithelial cell marker in periportal compartments). Scale bars are 200μm (PSR and CYP2E1 images) and 50μm (CK19 images).

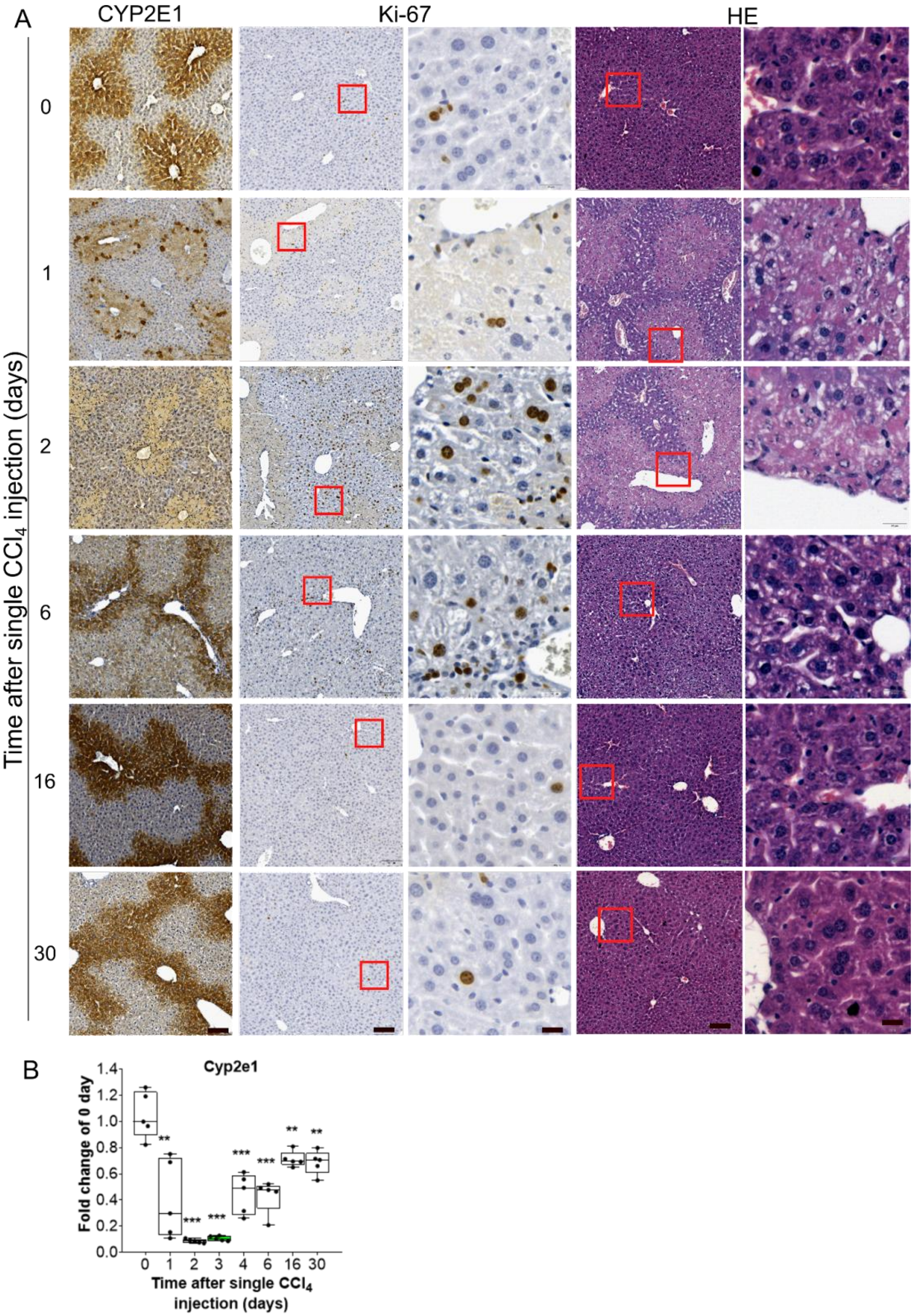
A
CYP2E1
Ki-67
HE
Time after single $CCl_4$ injection (days)
0
1
2
6
16
30
B
Cyp2e1
Fold change of 0 day
1.4
1.2
1.0
0.8
0.6
0.4
0.2
0.0
**
***
***
***
***
**
**
0 1 2 3 4 6 16 30
Time after single $CCl_4$ injection (days)

**Supplementary Figure 3. Regeneration upon acute liver injury in mice from one $CCl_4$ injection.** (A) Representative time-resolved images of CYP2E1, Ki-67 and HE stained livers upon one dose of $CCl_4$. Scale bars are 100µm and 20µm for overviews and closeups, respectively. (B) mRNA level of Cyp2e1, determined at different time points after administration of $CCl_4$ as indicated. Data are shown as the means ± SD of 5 mice per group. The values at day 0 represent the controls. ** P <0.01, *** P <0.001.

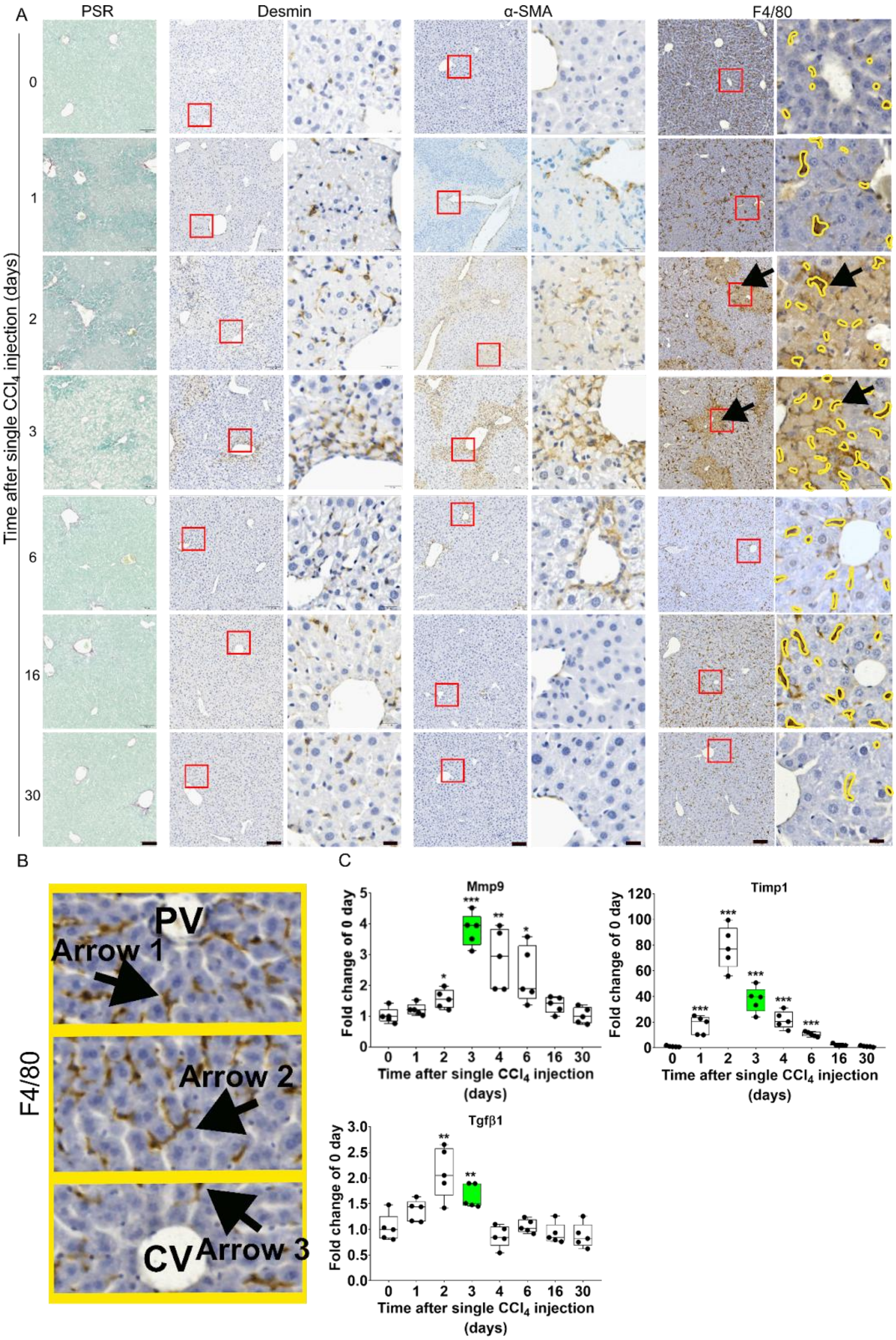
A
PSR
Desmin
α-SMA
F4/80
Time after single CCl4 injection (days)
0
1
2
3
6
16
30
B
F4/80
PV
Arrow 1
Arrow 2
Arrow 3
CV
C
Mmp9
Timp1
Tgfβ1
Fold change of 0 day
Time after single CCl4 injection (days)
0 1 2 3 4 6 16 30

**Supplementary Figure 4. ECM, HSC and macrophages during liver regeneration upon one $CCl_4$ injection.** (A) Time-resolved histopathological data after a single $CCl_4$ injection, showing PSR (ECM), desmin (quiescent and activated HSC), α-SMA (activated HSC), F4/80 (resident macrophages, highlighted in yellow). Desmin, α-SMA and F4/80 are each displayed at overview-magnification (left, scale bar 100μm) and high magnification for closeup (right, scale bar 20μm). (B) The close view of Fig. 3A, F4.80. (C) Time resolved RT-PCR data for Tgfβ1, Mmp9 and Timp1. Data are shown as the means ± SD of 5 mice per group. * $P < 0.05$, ** $P < 0.01$, *** $P < 0.001$.

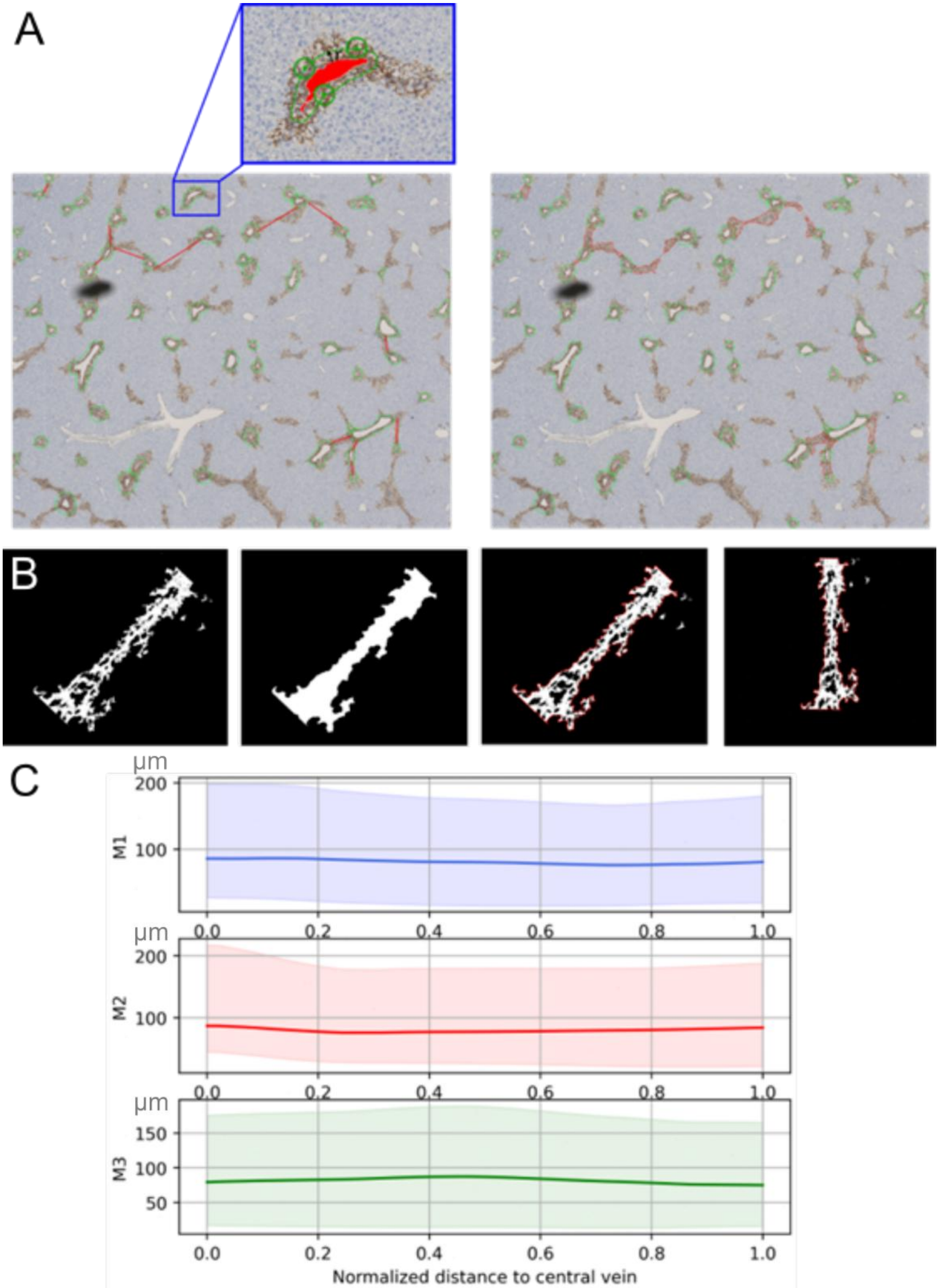
A
B
C
µm
200
100
M1
0.0
0.2
0.4
0.6
0.8
1.0
µm
200
100
M2
µm
150
100
50
M3
Normalized distance to central vein

**Supplementary Figure 5**. **Computation of the distribution of the spanning width of α-SMA positive cells.** (A) Example of a region of interest showing the activated HSCs in brown; brown regions with void representing CVs are marked by green circles. In the left subfigure, the red lines are drawn manually to identify neighboring CVs connected by areas occupied by activated HSCs; in the right subfigure the red curved lines demarcate the borders of these areas ("stripes"). (B) The different steps to obtain the envelope (border) of an aHSC positive area (stripe) connecting two neighboring CVs. Each stripe is finally oriented vertically before computing its spanning width along the axis starting at one CV and ending at the neighboring CV. (C) The resulting values for the width of the stripe along the CV-CV connection is then normalized by division of the length of the CV-CV connection to permit accumulation of the data for CV-CV connections of different lengths (shown are accumulated data, each from multiple CV-CV connections from different mice M1, M2, and M3). Interestingly, the width of the stripes does not vary along the CV-CV connection.

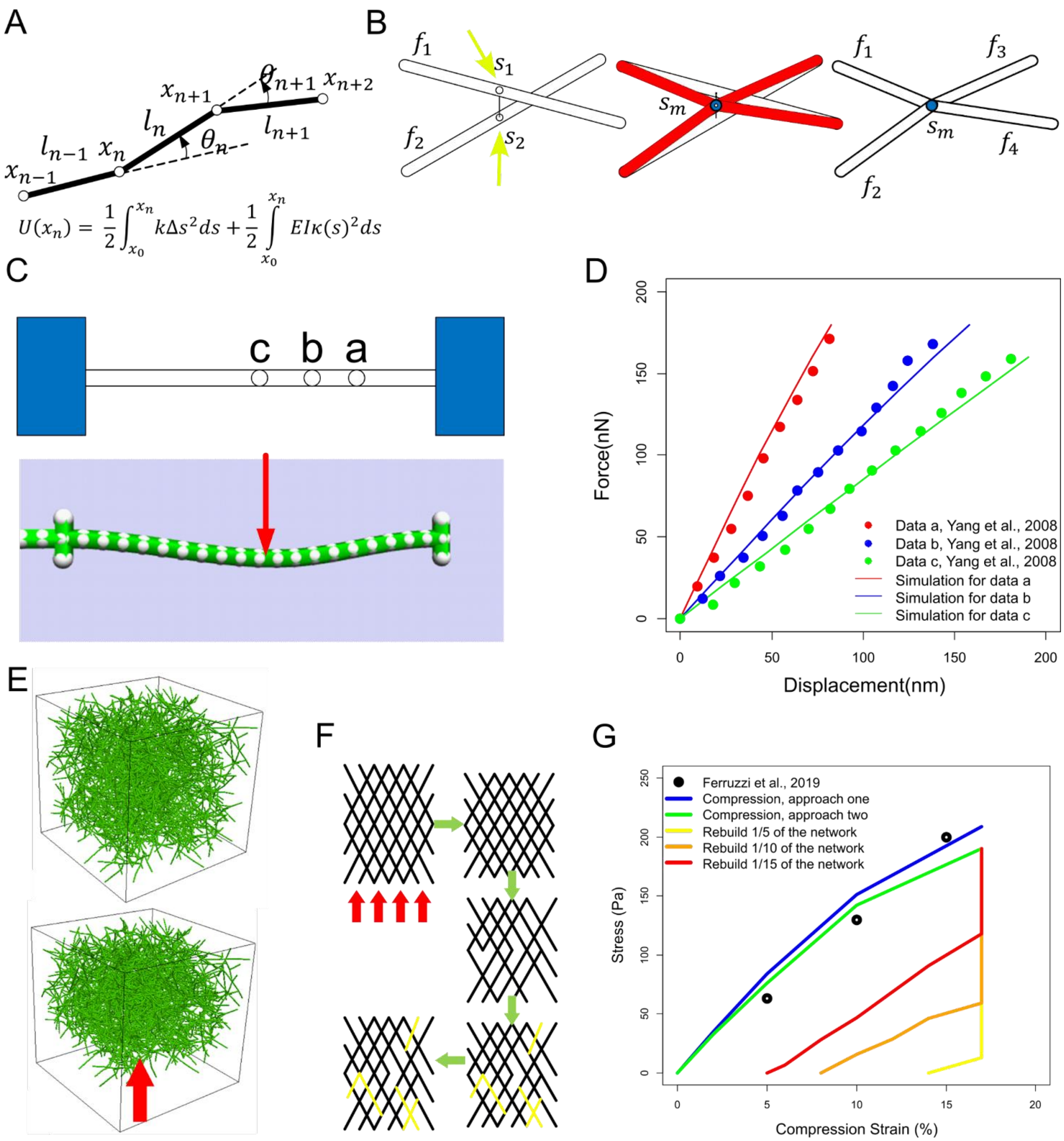


**Supplementary Figure 6**. **Construction and validation of the collagen (ECM) model**. (A) Elastic and bending energy of the collagen fiber network. (B) Addition of crosslink nodes: If the distance between two fiber segments $f_1$ and $f_2$ (between two perpendicular nodes $s_1$ and $s_2$ on $f_1$ and $f_2$, respectively, indicated by yellow arrows) is smaller than a predefined threshold, a crosslink node sm (blue) is added, where $f_1$ and $f_2$ would crosslink (red). After crosslinking, the segments are reenumerated: $f_1$ and $f_2$ are divided generating two additional segments ($f_3$ and $f_4$), which are all attached at $s_m$ (blue). (C) Calibration of mechanical parameters for the bundles mimicking the bending test by Yang et al.[16]. ‘a’, ‘b’, and ‘c’ indicate the corresponding positions where the force is applied on the fiber. The simulated fiber (length of 3 μm) bends upon the exerted force (red arrow). (D) Simulation result and experimental data from Yang et al.[24]. (E) The modeled compression of the collagen network. The size of the network is 42.8×42.8×42.8μm3. The network is compressed by fixing the collagen nodes on the top layer and moving the bottom layer of collagen nodes upwards. (F) Rebuild the network for plasticity test: After the 3D network of cross-linked fibers (from D, E) is compressed, some fibers are randomly removed from and subsequently randomly added back to the network; subsequently, the compressed network is relaxed. This algorithm mimics a visco-elastic network with a plastic response. (G) Strain-stress curve of simulation and experimental data from Ferruzzi et al.[25]. Different fractions of total fibers rebuilt (removed first and then added back) in the network are tested.

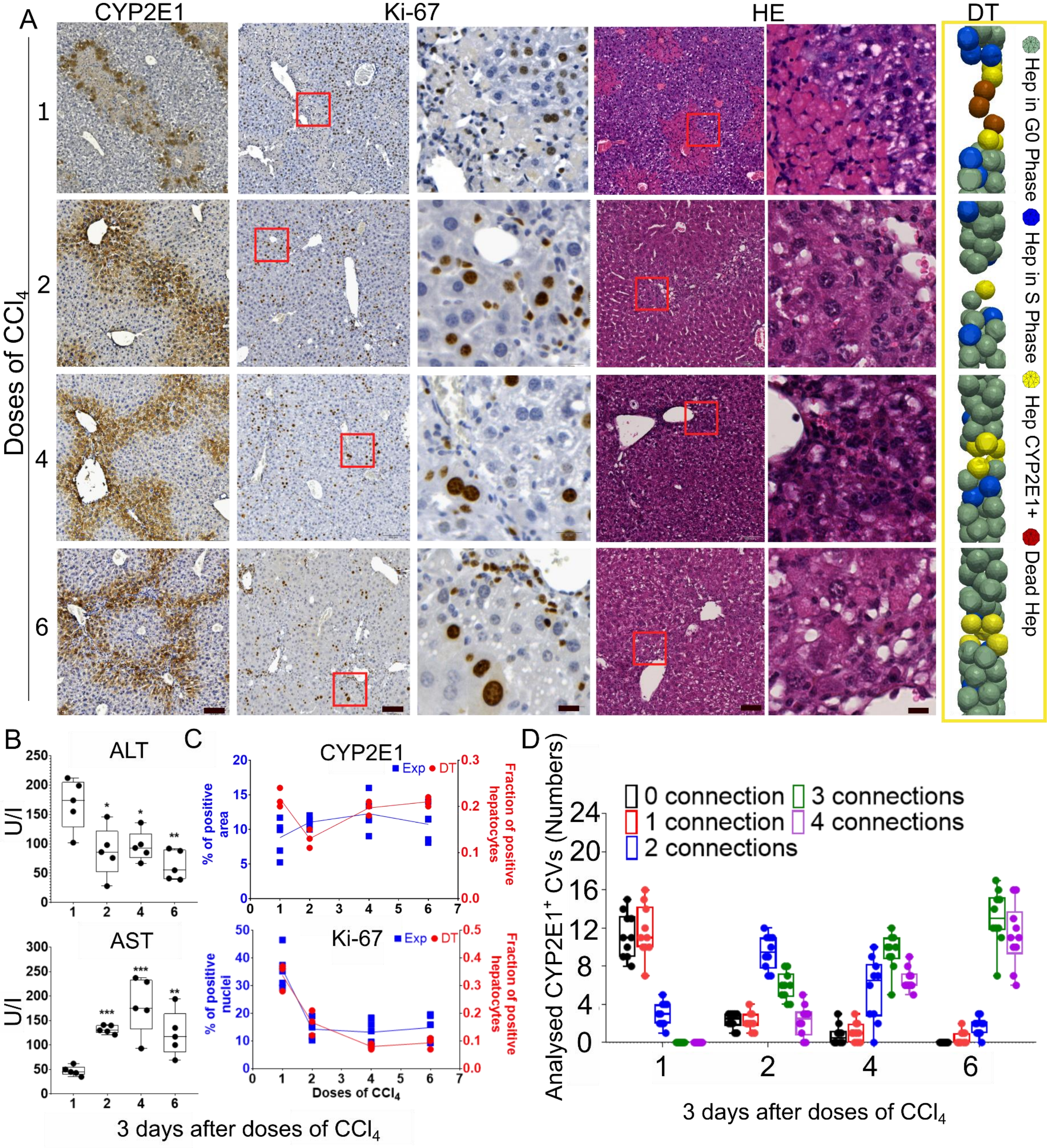


**Supplementary Figure 7. Chronic liver injury in mice from 6 injections of $CCl_4$.** (A) Representative images of CYP2E1, Ki-67 and HE stained livers plus snapshots of the DT for repeated doses of $CCl_4$ (scale bars are 100µm for overview and 20µm for closeup images). The corresponding days are day 3 (3 days after dose 1), 7 (4 days after dose 2), 14 (4 days after dose 4), 21 (4 days after dose 6). (B) Blood levels of liver transaminases (AST and ALT), upon repetitive $CCl_4$ injections (compare Fig. 2D for the control). (C) (Top) Quantification of the experimentally determined CYP2E1 positive area and fraction of CYP2E1 positive cells in the DT. (Bottom) Quantification of Ki-67 positive nuclei and fraction of S-phase hepatocytes in the DT (for both fractions of CYP2E1 positive and S-phase hepatocytes, the number of cells is taken power of 2/3 to compare the simulation result to the experimentally analyzed area $A \sim V^{2/3}$). (D) Quantification of CVs connected by CYP2E1 positive hepatocytes. Experimental data are shown as the means ± SD of 5 mice per group. * $P < 0.05$, ** $P < 0.01$, *** $P < 0.001$. DT simulation data are shown as the means ± SD of 3 independent simulation runs. The CYP2E1+ cells are chosen according to the experimental observation of CYP2E1+ cells and those cells dying upon $CCl_4$-administration (see "**Digestion of dead hepatocytes by macrophages"** above). At day 0 before injection, the cells located in the region spanning 8 layers of the cells in the central are chosen as CYP2E1+ cells (corresponding to a radius of about 4 cell layers around the CV in a liver lobule (see Hoehme et. al.[7]). From the 2nd injection on the experimental stainings and the concentration of AST suggest that after regeneration, the CYP2E1+ cell layer is only two cells thick (see also **Fig. 2**). Hence the two innermost cell layers along the CV-CV-axis are marked as CYP2E1+.

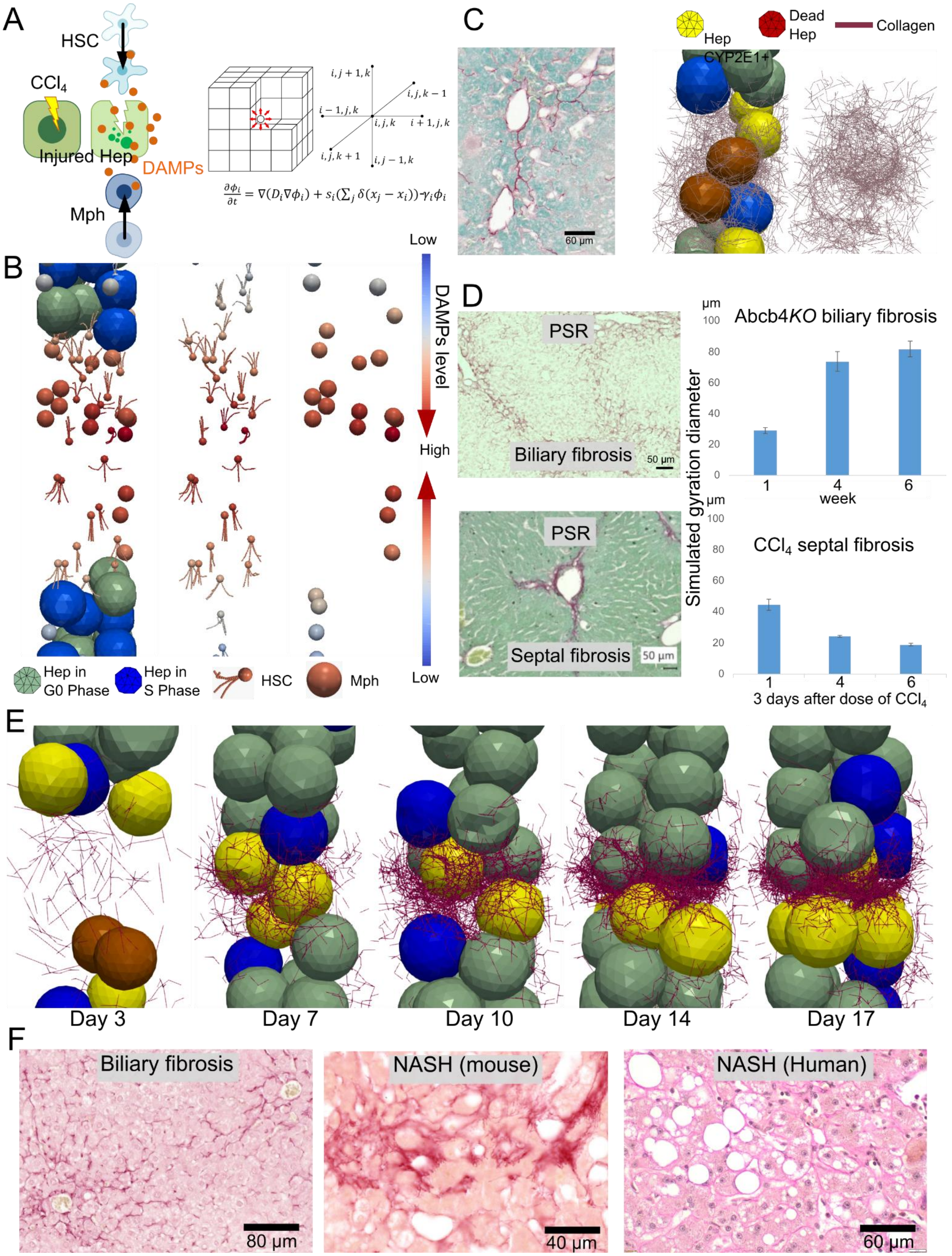


**Supplementary Figure 8. Migration of HSC and Mph to mediate liver fibrosis formation.** (A)

Scheme of DAMPs gradient in the model (DT). Injured Hep produces DAMPs. DAMPs can activate HSCs and macrophage (Mph) to migrate towards the largest DAMP gradient. The spatio-temporal DAMP concentration is calculated by solving a differential equation that takes into account synthesis, diffusion, and decay of DAMPs. The differential equations are solved in a discretized cubic system. (B) Snapshots representative for migrating HSCs and Mphs from low (blue) to high (red) DAMP concentrations (**Supplementary Movie 3**). (C) Chicken wire pattern of liver fibrosis (collagen fibers wrap liver cells), as observed in $CCl_4$ mouse model and in DT simulation. (D) The staining of collagens (PSR) in both biliary and septal fibrosis at the same scale (scale bar: 50 μm) were compared along with the simulation result of gyration diameter of the collagen network of both biliary and septal fibrosis. (E) The snapshots of collagen fibers at days 3, 7, 10, 14, 17. (F) The chicken wire pattern observed in biliary fibrosis (Abcb4*KO* mouse model), and NASH (mouse model and human patient).

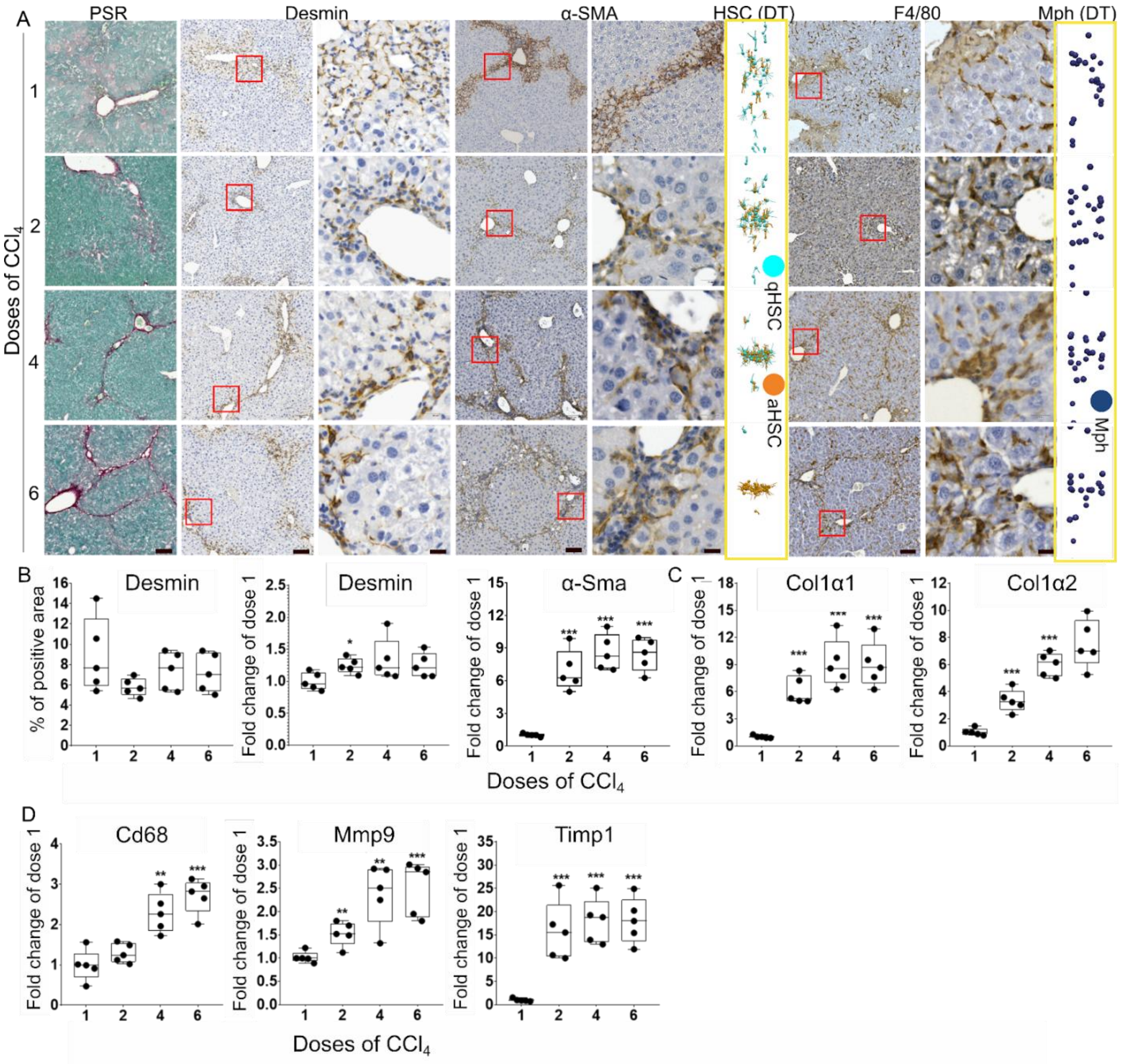


**Supplementary Figure 9. HSC, Mph and matrisome analysis in the dynamics of $CCl_4$ mediated chronic liver disease. (**A) Representative images of PSR, desmin, α-SMA, and F4/80 stained livers and snapshots of HSC and Mph in the DT simulation. (B-D) Protein/mRNA levels of desmin, α-SMA, Collagen1α1, Collagen1α2, Cd68, Mmp9, and Timp1 determined by IHC or RT-PCR, as indicated. Scale bars are 100μm for overview and 20μm for closeup images. Experimental data are shown as the means ± SD of 5 mice per group. * P <0.05, ** P <0.01, *** P <0.001. DT simulation data are shown as the means ± SD of 3 independent simulation runs.

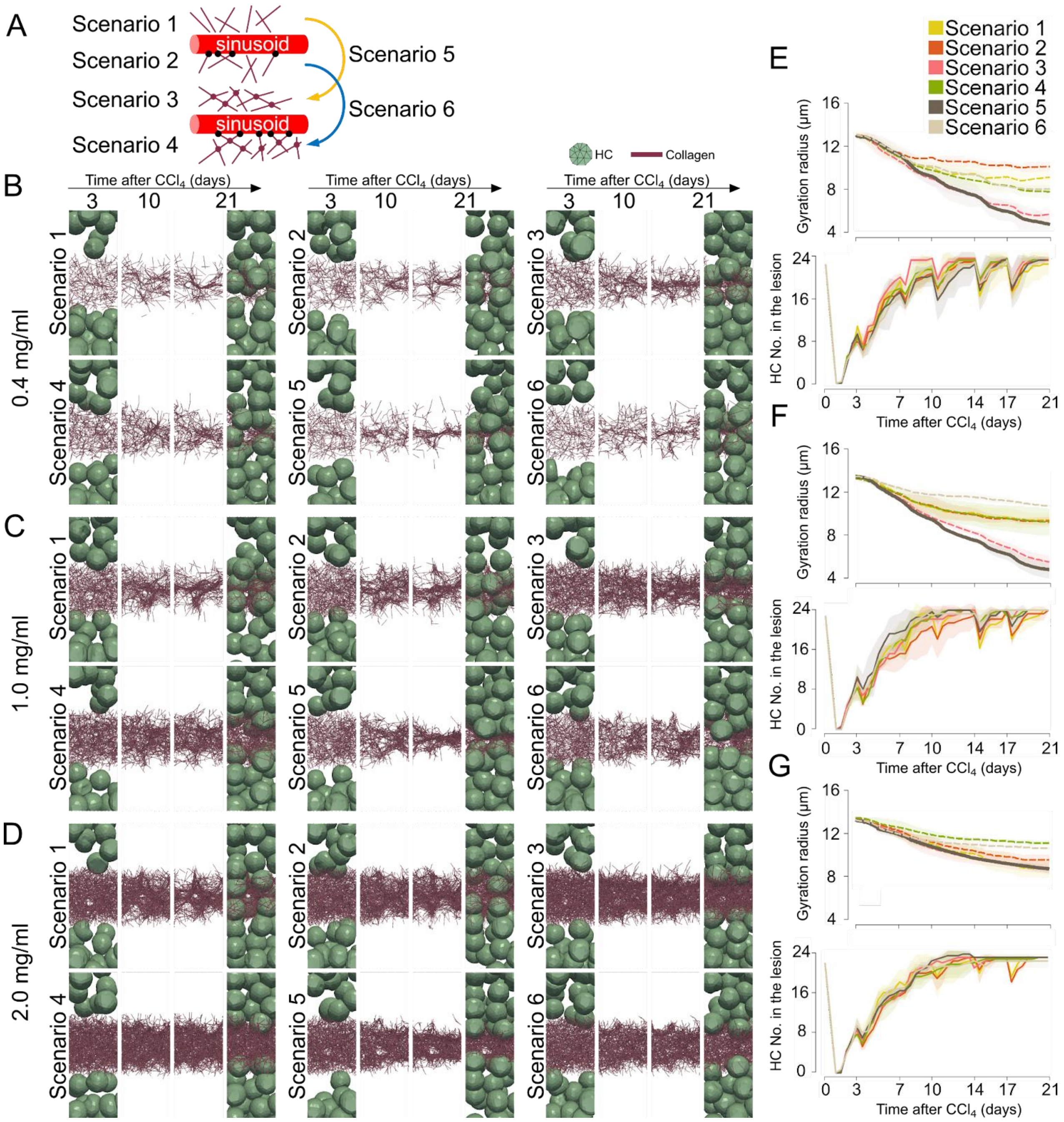


**Supplementary Figure 10**. **Organizational structure of the collagen network**. (A) There are six scenarios (S) for the structure of collagen deposition. Collagen fibers are deposited as single fibers in S1, single fibers anchored to the nearest sinusoids in S2, crosslinked single fibers in S3, crosslinked single fibers anchored to the nearest sinusoids in S4. In S5, collagen fibers are firstly deposited as single fibers and then gradually crosslinked while in S6, collagen fibers are firstly deposited as single fibers anchored to the nearest sinusoids and then gradually crosslinked. (B-D) Snapshots of collagen fiber organization as resulting from the different simulation scenarios with the DT based on assuming collagen densities of 0.4 mg/mL, 1.0 mg/mL and 2.0 mg/mL, respectively. (E-G) Gyration radius of the fibers and number of hepatocytes in the lesion in the different scenarios. Scenario 5 in the graph depicting the gyration radius is highlighted in a solid line, while the other scenarios are presented in dashed lines. The error bars represent the standard deviation of 3 simulation runs.

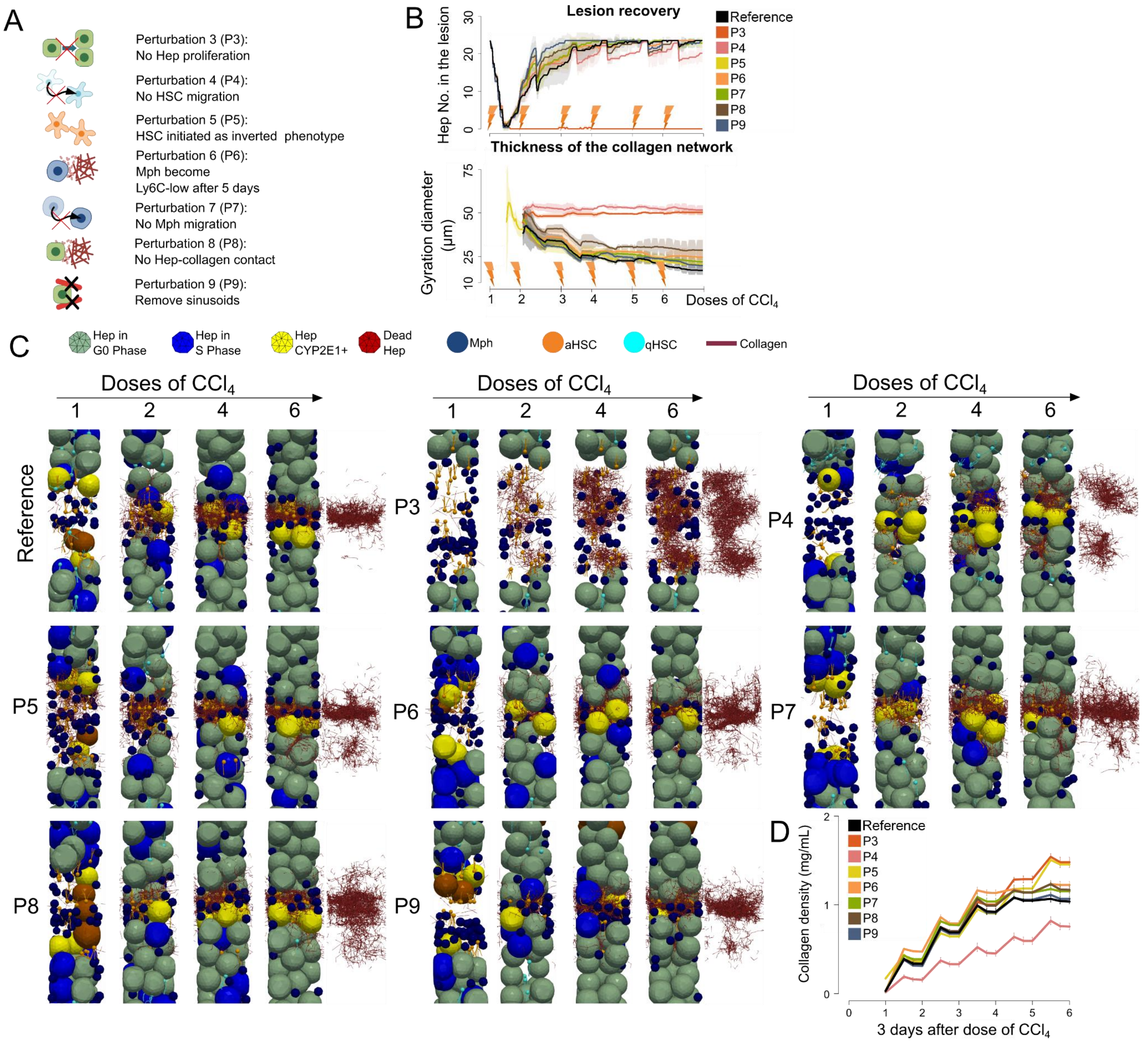


**Supplementary Figure 11**. **DT simulation of different perturbation cases**. (A) Seven more perturbation scenarios in addition to those in **Fig. 5** are tested. P3, Disable hepatocyte proliferation; P4, Disable HSC migration; P5, HSCs are initiated as inverted phenotype after $CCl_4$ injections; P6, macrophages become Ly6C$^{low}$, 5 days after $CCl_4$ injections; P7, Disable macrophage migration; P8, disable hepatocyte-collagen interaction; P9, remove sinusoids. (B) Number of hepatocytes in the lesion and thickness of the collagen network during 3 weeks of chronic $CCl_4$ injections for all perturbation scenarios. Thickness is determined by the gyration diameter of the network. The error bars represent the standard deviation of 3 simulation runs. (C) Snapshots of the simulation of all perturbation scenarios at different time spots. Typical time-lapse DT simulations are shown as **Supplementary Movies 6-12**. (D) Plot of collagen density in the lesion region at the days of injection of $CCl_4$ (day 3, 7, 10, 14, 17, 21), and at days between injections (day 5, 6, 8.5, 9, 12, 13, 15.5, 16, 19, 20) for all perturbation scenarios.

### Supplementary Tables

#### Supplementary table 1. Sequences of primer pairs used for real time PCR

| Gene | Species | Forward | Reverse |
|---|---|---|---|
| Acta2 (α-Sma) | Mouse | TTCGCTGTCTACCTTCCAGC | GAGGCGCTGATCCACAAAAC |
| Timp1 | Mouse | GGCATCTGGCATCCTCTTGT | ACTCTTCACTGCGGTTCTGG |
| Cd68 | Mouse | GGCGGTGGAATACAATGTGTCC | AGCAGGTCAAGGTGAACAGCTG |
| Col1α1 | Mouse | ACGTGGAAACCCGAGGTATG | TTGGGTCCCTCGACTCCTAC |
| Col1α2 | Mouse | AGTCGATGGCTGCTCCAAAA | AGCACCACCAATGTCCAGAG |
| Cyp2e1 | Mouse | CGTTGCCTTGCTTGTCTGGA | AAGAAAGGAATTGGGAAAGGTCC |
| Desmin | Mouse | TACACCTGCGAGATTGATGC | ACATCCAAGGCCATCTTCAC |
| Mmp9 | Mouse | GCAGAGGCATACTTGTACCG | TGATGTTATGATGGTCCCACTTG |
| Ppia | Mouse | GAGCTGTTTGCAGACAAAGTC | CCCTGGCACATGAATCCTGG |
| Tgfβ1 | Mouse | AGGGCTACCATGCCAACTTC | CCACGTAGTAGACGATGGGC |

**Supplementary Table 2. DT (digital twin) parameters**.

| Description | Value | Reference |
|---|---|---|
| Hepatocyte radius | ~10.7 μm | Estimated from data |
| Sinusoid radius | ~2.1 μm | Hoehme et al.[7] (2010) |
| HSC radius | ~2 μm | Wake[37] (2006) |
| HSC branch length | ~12 μm | Wake[37] (2006) |
| Mph radius | ~6 μm | Shi et al.[38] (2011) |
| Collagen fiber radius | ~0.1 μm | Wenger et al.[39] (2007) |
| Collagen fiber segment length | ~3 μm | Stein et al.[17] (2008) |
| Young's modulus of hepatocyte | ~400 Pa | Hoehme et al.[7] (2010) |
| Young's modulus of sinusoid | ~600 Pa | Hoehme et al.[7] (2010) |
| Young's modulus of HSC | ~700 Pa | Estimated from fibroblast, Yang et al.[40] (2012) |
| Young's modulus of macrophage | ~1400 Pa | Estimated from immune macrophage, Bufi et al.[41] (2015) |
| Young's modulus of collagen fiber | ~50 MPa | Manssor et al.[42] (2016) |
| Poisson ratio of all components | 0.4 | Hoehme et al.[7] (2010) |

| | | |
|---|---|---|
| Cell cycle time of hepatocyte | 24 hours | Hoehme et al.[7] (2010) |
| Cell cycle time of HSC | 24 hours | Estimated |
| Medium friction for hepatocyte, sinusoid and macrophage | $10^8$ Ns/m$^3$ | Estimated |
| Medium friction for HSC | $10^{10}$ Ns/m$^3$ | Estimated |
| Medium friction for collagen fiber | $10^{11}$ Ns/m$^3$ | Estimated |
| Friction between all components | $10^8$ Ns/m$^3$ | Estimated |
| HSC density in the liver | ~1/70 μm of the sinusoid | Wake[37] (2006) |
| Mph density in the liver | ~2×$10^4$/mm$^3$ | Bouwens et al.[43] (1986) |
| Collagen density | ~1 mg/ML | Stein et al.[17] (2008) |
| Local collagen density restriction of further collagen production by aHSC | ~10 mg/ML | Estimated |
| Mean of migration speed of HSC | ~2.1 μm/hour | Tangkijvanich et al.[30] (2001) |
| SD of migration speed of HSC | ~0.1 μm/hour | Tangkijvanich et al.[30] (2001) |
| Mean of migration speed of Mph | ~5 μm/min | Grabher et al.[29] (2007) |

| | | |
|---|---|---|
| SD of migration speed of Mph | ~1.8 μm/min | Grabher et al.[29] (2007) |
| Diffusion rate of DAMPs, estimated from its molecular weight, 28 kDa | ~$2.5\times10^{-11}$ $m^2/s$ | Davies et al.[44] (2018) |
| Concentration of DAMPs to trigger Mph to migrate | 0.1 ng/ml | estimated |
| Concentration of DAMPs to trigger HSC to migrate | 0.1 ng/ml | estimated |
| Decay rate of DAMPs, estimated from its half-life, ~1000 seconds | ~$5.7\times10^{-4}$ /s | Zandarashvili et al.[45] (2013) |
| Fraction of HSC switching to inverted phenotype | 0.5 | Kisseleva et al.[32] (2012) |
| Fraction of apoptosis HSC after each injection of $CCl_4$ | 0.5 | Kisseleva et al.[32] (2012) |
| Time of macrophage switching to $Ly6C^{low}$ phenotype after each injection of $CCl_4$ | 12 hours | Estimated from data |
| Engulfment and elimination duration of Mph | ~3 hours | Estimated from Haecker et al.[34] (2002) |

| | | |
|---|---|---|
| The width of the lesion size | ~80 μm | Estimated from data |

**Supplementary Movies**

**Supplementary Movie 1.** 3D reconstruction of a hepatocyte (yellow), nucleus (blue) and wrapped by sinusoids (red). This cell reconstruction is generated from confocal images obtained from healthy mouse liver[46].

**Supplementary Movie 2.** Time-Lapse video of a typical model simulation of the formation of fibrotic pattern in the reference model (Light green: healthy hepatocyte; Yellow: CYP2E1+ hepatocyte; Brown: dead hepatocyte; Blue: dividing hepatocyte; Dark Blue: Mph; Cyan: qHSC; Orange: aHSC; Dark purple: collagen fiber. This color set is consistent across subsequent movies).

**Supplementary Movie 3.** Time-Lapse video of a typical model simulation showing migration of HSC and Mph towards the gradient of DAMPs. Mph is represented as a sphere and HSC is represented as a sphere attached with several chains.

**Supplementary Movie 4.** Time-Lapse video of a typical model simulation of the perturbation 1 (randomly distributed CYP2E1+ cells) upon the reference model. collagen networks are randomly generated in the regions where hepatocytes are killed.

**Supplementary Movie 5.** Time-Lapse video of a typical model simulation of the perturbation 2 (inhibit the FAK signal) upon the reference model. Fewer activated HSCs and collagen fibers are generated compared to the reference model.

**Supplementary Movie 6.** Time-Lapse video of a typical model simulation of the perturbation 3 (no hepatocyte proliferation) upon the reference model. No hepatocyte proliferation occurs, and collagen networks are generated in the lesion site without mechanical contacts with hepatocytes.

**Supplementary Movie 7.** Time-Lapse video of a typical model simulation of the perturbation 4 (no HSC migration) upon the reference model. The collagen fibers are generated close to the border of the lesion, not in the center of the lesion.

**Supplementary Movie 8.** Time-Lapse video of a typical model simulation of the perturbation 5 (HSC is initiated as inverted phenotype) upon the reference model. Collagen fibers are generated earlier but the general pattern remains the same as that in the reference.

**Supplementary Movie 9.** Time-Lapse video of a typical model simulation of the perturbation case 6 (macrophage becomes Ly6C$^{low}$ after 5 days) upon the reference model. There are more collagen fibers without being digested by macrophages, therefore the gyration diameter of the collagen network is larger than that in the reference.

**Supplementary Movie 10.** Time-Lapse video of a typical model simulation of the perturbation 7 (no macrophage migration) upon the reference model. There are more collagen fibers without being digested by macrophages, therefore the gyration diameter of the collagen network is larger than that in the reference.

**Supplementary Movie 11.** Time-Lapse video of a typical model simulation of the perturbation 8 (no hepatocyte-collagen interaction force) upon the reference model. Collagen fibers are expanding a much wider range than that in the reference, therefore the gyration diameter of the collagen network is larger.

**Supplementary Movie 12.** Time-Lapse video of a typical model simulation of the perturbation 9 (no sinusoids) upon the reference model. The gyration diameter of the collagen network is larger than that in the reference.

### References


1. Gonzalez, R.C., Woods, R.E. *Digital Image Processing*. (Pearson/Prentice Hall, 2010).
2. Soille, P. *Morphological image analysis: principles and applications. Vol. 2, No. 3.* (Berlin: Springer,1999).
3. Sternberg, S.R. Grayscale morphology. *Computer Vision, Graphics, and Image Processing* 35(3), 333-355 (1986).
4. Drasdo, D., Hoehme, S. A single-cell-based model of tumor growth in vitro: monolayers and spheroids. *Phys. Biol.* 2, 133 (2005).
5. Drasdo, D., Hoehme, S. Modeling the impact of granular embedding media and pulling versus pushing cells on growing cell clones. *New J. Phys.* 14, 055025 (2012).
6. Hoehme, S., Hammad, S., Boettger, J., Begher-Tibbe, B., Bucur, P., Vibert, E., Gebhardt, R., Hengstler, J.G., Drasdo, D. Digital twin demonstrates significance of biomechanical growth control in liver regeneration after partial hepatectomy. *iScience* 26, 105714 (2023).
7. Hoehme, S., Brulport, M., Bauer, A., Bedawy, E., Schormann, W., Hermes, M., Puppe, V., Gebhardt, R., Zellmer, S., Schwarz, M., Bockamp, E., Timmel, T., Hengstler, J.G., Drasdo, D. Prediction and validation of cell alignment along microvessels as order principle to restore tissue architecture in liver regeneration. *Proc. Natl Acad Sci USA* 107(23), 10371-10376 (2010).
8. Van Liedekerke, P., Neitsch, J., Johann, T., Alessandri, K., Nassoy, P., Drasdo, D. Quantitative cell-based model predicts mechanical stress response of growing

tumor spheroids over various growth conditions and cell lines. *PLoS Computational Biology* 15(3), e1006273 (2019).

9. Zhao, J., Ghallab, A., Hassan, R., Dooley, S., Hengstler, J.G., Drasdo, D. A digital twin of liver predicts regeneration after drug-induced damage at the level of cell type orchestration. *iScience* 27, 108077 (2024).
10. Drasdo, D., Hoehme, S., Hengstler, J.G. How predictive quantitative modelling of tissue organisation can inform liver disease pathogenesis. J. Hepatol. 61, 9151--956 (2014).
11. Van Liedekerke, P., Palm, M.M., Jagiella, N., Drasdo, D. Simulating tissue mechanics with agent-based models: concepts, perspectives and some novel results. *Computational Particle Mechanics* 2, 401-444 (2015).
12. Odenthal, T., Smeets, B., Van Liedekerke, P., Tijskens, E., Van Oosterwyck, H., Ramon, H. Analysis of initial cell spreading using mechanistic contact formulations for a deformable cell model. *PLoS Computational Biology* 9(10), e1003267 (2013).
13. Kim, M., Siberberg, Y.R., Abeyaratne, R., Asada, H.H. Computational modeling of three-dimensional ECM-rigidity sensing to guide directed cell migration. *Proc. Natl Acad Sci USA* 115(3), E390-399 (2018).
14. Winkler, B., Aranson, I.S., Ziebert, F. Confinement and substrate topography control cell migration in a 3D computational model. *Communications Physics* 2, 82 (2019).
15. Van Liedekerke, P., Neitsch, J., Johann, T., Warmt, E., Gonzalez-Valverde, I., Hoehme, S., Grosser, S., Kaes, J., Drasdo, D. A quantitative high-resolution

computational mechanics cell model for growing and regenerating tissues. *Biomechanics and Modeling in Mechanobiology* 19, 189-220 (2020).

16. Levick, J.R. Flow through interstitium and other fibrous matrices. *Quarterly Journal of Experimental Physiology* 72(4), 409-437 (1987).
17. Stein, A.M., Vader, D.A., Jawerth, L.M., Weitz, D.A., Sander, L.M. An algorithm for extracting the network geometry of three-dimensional collagen gels. *Journal of Microscopy* 232, 463-475 (2008).
18. Allens, M.P., Tildesley, D.J. *Computer Simulation of Liquids*. (Oxford University Press, 1987).
19. Yang, K., Lu, C., Zhao, X., Kawamura, R. From bead to rod: Comparison of theories by measuring translational drag coefficients of micron-sized magnetic bead-chains in Stokes flow. *PLoS ONE* 12(11), e0188015 (2017).
20. Cox, R.G. The motion of long slender bodies in a viscous fluid Part 1. General theory. *Journal of Fluid Mechanics* 44, 791-810 (1970).
21. Popov, V.L. *Contact mechanics and friction*. (Springer, 2010).
22. Karabassi, E., Papaioannou, G., Theoharis, T., Boehm, A. Intersection test for collision detection in particle systems. *Journal of Graphics Tools* 4(1), 25-37 (1999).
23. Sneddon, I.N. The relation between load and penetration in the axisymmetric boussinesq problem for a punch of arbitrary profile. *International Journal of Engineering Science* 3(1), 47-57 (1965).
24. Yang, L., Fitie, C.F.C., Van der Werf, K.O., Bennink, M.L., Dijkstra, P.J., Feijen, J. Mechanical properties of single electrospun collagen type I fibers. *Biomaterials*

29, 955-962 (2008).

25. Ferruzzi, J., Sun, M., Gkousioudi, A., Pillar, A., Roblyer, D., Zhang, Y., Zaman, M.H. Compressive remodeling alters fluid transport properties of collagen networks - implications for tumor growth. *Scientific Reports* 9, 17151 (2019).
26. Ghosh, S., Kaplan, K.J., Schrum, L.W., Bonkovsky, H.L. Chapter Five – cytoskeletal proteins: shaping progression of Hepatitis C virus-induced liver disease. *International Review of Cell and Molecular Biology* 302, 279-319 (2013).
27. Calderwood, S.K., Gong, J., Murshid, A. Extracellular HSPs: the complicated roles of extracellular HSPs in immunity. *Frontiers in Immunology* 7, 159 (2016).
28. Martin-Murphy, B.V., Holt, M.P., Ju, C. The role of damage associated molecular pattern molecules in acetaminophen-induced liver injury in mice. *Toxicology Letters* 192: 387-394 (2010).
29. Grabher, C., Cliffe, A., Miura, K., Hayflick, J., Pepperkik, R., Rørth, P., Wittbrodt, J. Birth and life of tissue macrophages and their migration in embryogenesis and inflammation in medaka. *Journal of Leukocyte Biology* 81(1), 263-271 (2007).
30. Tangkijvanich, P., Tam, S.P., Yee, J.F. Wound-induced migration of rat hepatic stellate cells is modulated by endothelin-1 through Rho-kinase-mediated alterations in the acto-myosin cytoskeleton. *Hepatology* 3(1), 74-80 (2001).
31. Iredale, J.P., Benyon, R.C., Pickering, J., McCullen, M., Northrop, M, Pawley, S., Hovell, C., Arthur, M.J. Mechanisms of spontaneous resolution of rat liver fibrosis. Hepatic stellate cell apoptosis and reduced hepatic expression of metalloproteinase inhibitors. *The Journal of Clinical Investigation* 102(3), 538-549 (1998).

32. Kisseleva, T., Cong, M., Paik, Y., Scholten, D., Jiang, C., Benner, C., Iwaisako, K., Moore-Morris, T., Scott, B., Tsukamoto, H., Evans, S.M., Dillmann, W., Glass, C.K., Brenner, D.A. Myofibroblasts revert to an inactive phenotype during regression of liver fibrosis. *Proc. Natl Acad Sci USA* 109(24), 9448-53 (2012).
33. Lee, J.I., Lee, K.S., Paik, Y.H., Park, Y.N., Han, K.H., Chon, C.Y., Moon, Y.M. Apoptosis of hepatic stellate cells in carbon tetrachloride induced acute liver injury of the rat: analysis of isolated hepatic stellate cells. J. Hepatol. 39, 960--966 (2003).
34. Haecker, H., Fuermann, C., Wagner, H., Haecker, G. Caspase-9/-3 activation and apoptosis are induced in mouse macrophages upon ingestion and digestion of Escherichia coli bacteria. *J. Immunol.* 169, 3172-3179 (2002).
35. Montagud, A., Beal, J., Tobalina, L., Traynard, P., Subramanian, V., Szalai, B., Alfoldi, R., Puskas, L., Valencia, A., Barillot, E., Saez-Rodriguez, J., Calzone, L. Patient-specific boolean models of signalling networks guide personalised treatments. eLife 11, e72626 (2022).
36. Dichamp, J., Celliere, G., Ghallab, A., Hassan, R., Boissier, N., Hofmann, U., Reinders, J., Sezgin, S., Zuehlke, S., Hengstler, J.G., Drasdo, D. In vitro to in vivo acetaminophen hepatotoxicity extrapolation using classical schemes, pharmacodynamic models and a multiscale spatial-temporal liver twin. *Front. Bioeng. Biotechnol.* 11, 1049564 (2023).
37. Wake, K. Hepatic stellate cells: Three-dimensional structure, localization, heterogeneity and development. *Proceedings of the Japan Academy, Series B* 82(4), 155-164 (2006).

38. Shi, C., Jia, T., Mendez-Ferrer, S., Hohl, T.M., Serbina, N.V., Lipuma, L., Leiner, I., Li, M.O., Frenette, P.S., Pamer, E.G. Bone marrow mesenchymal stem and progenitor cells induce monocyte emigration in response to circulating Toll-like receptor ligands. *Immunity* 34(4), 590-601 (2011).
39. Wenger, M.P., Bozec, L., Horton, M.A., Mesquida, P. Mechanical properties of collagen fibrils. *Biophysical Journal* 93(4), 1255-1263 (2007).
40. Yang, Y., Liao, J., Lin, C., Chang, C., Wang, S., Ju, M. Characterization of cholesterol-depleted or -restored cell membranes by depth-sensing nano-indentation. *Soft Matter* 8(3): 682-687 (2012).
41. Bufi, N., Saitakis, M., Dogniaux, S., Buschinger, O., Bohineust, A., Richert, A., Maurin, M., Hivroz, C., Asnacios, A. Human primary immune cells exhibit distinct mechanical properties that are modified by inflammation. *Biophysical Journal* 108(9), 2181-2190 (2015).
42. Manssor, N.A.S., Radzi, Z., Yahya, N.A., Yusof, L.M., Hariri, F., Khairuddin, N.H., Kasim, N.H.A., Czernuszka, J.T. Characteristics and Young's modulus of collagen fibrils from expanded skin using anisotropic controlled rate self-inflating tissue expander. *Skin Pharmacology and Physiology* 29(2), 55--62 (2016).
43. Bouwens, L., Knook, D.L., Wisse, E. Local proliferation and extrahepatic recruitment of liver macrophages (Kupffer cells) in partial-body irradiated rats. *Journal of Leukocyte Biology* 39, 687-697 (1986).
44. Davies, J.E., Apta, B.H.R., Harper, M.T. Cross-reactivity of anti-HMGB1 antibodies for HMGB2. Journal of Immunological Methods 456: 72—76 (2018).
45. Zandarashvili, L., Sahu, D., Lee, K., Lee, Y.S., Singh, P., Rajarathnam, K.,

Iwahara, J. Real-time kinetics of high-mobility group box 1 (HMGB1) oxidation in extracellular fluids studied by in Situ protein NMR spectroscopy. Journal of Biological Chemistry 288(17): 11621—11627 (2013).

46. Hammad, S., Hoehme, S., Friebel, A., Von Recklinghausen, I., Othman, A., Begher-Tibbe, B., Reif, R., Godoy, P., Johann, T., Vartak, A., Golka, K., Bucur, O.P., Vibert, E., Marchan, R., Christ, B., Dooley, S., Meyer, C., Ilkavets, I., Dahmen, U., Dirsch, O., Bottger, J., Gebhardt, R., Drasdo, D., Hengstler, J.G. Protocols for staining of bile canalicular and sinusoidal networks of human, mouse and pig livers, three-dimensional reconstruction and quantification of tissue microarchitecture by image processing and analysis. *Archives of Toxicology* 88, 1161-1183 (2014).
